# A 3D human iPSC-derived multi-cell type neurosphere system to model cellular responses to chronic amyloidosis

**DOI:** 10.1101/2024.07.01.601570

**Authors:** Stefan Wendt, Ada J. Lin, Sarah N. Ebert, Declan J. Brennan, Wenji Cai, Yanyang Bai, Da Young Kong, Stefano Sorrentino, Christopher J. Groten, Christopher Lee, Jonathan Frew, Hyun B. Choi, Konstantina Karamboulas, Mathias Delhaye, Ian R Mackenzie, David R. Kaplan, Freda Miller, Brian A. MacVicar, Haakon B. Nygaard

**Author notes:** Corresponding authors: Haakon B. Nygaard and Stefan Wendt.

## Abstract

**Background:** Alzheimer’s Disease (AD) is characterized by progressive amyloid beta (Aβ) deposition in the brain, with eventual widespread neurodegeneration. While the cell-specific molecular signature of end-stage AD is reasonably well characterized through autopsy material, less is known about the molecular pathways in the human brain involved in the earliest exposure to Aβ. Human model systems that not only replicate the pathological features of AD but also the transcriptional landscape in neurons, astrocytes and microglia are crucial for understanding disease mechanisms and for identifying novel therapeutic targets.

**Methods:** In this study, we used a human 3D iPSC-derived neurosphere model to explore how resident neurons, microglia and astrocytes and their interplay are modified by chronic amyloidosis induced over 3 to 5 weeks by supplementing media with synthetic Aβ1-42 oligomers. Neurospheres under chronic Aβ exposure were grown with or without microglia to investigate the functional roles of microglia. Neuronal activity and oxidative stress were monitored using genetically encoded indicators, including GCaMP6f and roGFP1, respectively. Single nuclei RNA sequencing (snRNA-seq) was performed to profile Aβ and microglia driven transcriptional changes in neurons and astrocytes, providing a comprehensive analysis of cellular responses.

**Results:** Microglia efficiently phagocytosed Aβ inside neurospheres and significantly reduced neurotoxicity, mitigating amyloidosis-induced oxidative stress and neurodegeneration following different exposure times to Aβ. The neuroprotective effects conferred by the presence of microglia was associated with unique gene expression profiles in astrocytes and neurons, including several known AD-associated genes such as *APOE*. These findings reveal how microglia can directly alter the molecular landscape of AD.

**Conclusions:** Our human 3D neurosphere culture system with chronic Aβ exposure reveals how microglia may be essential for the cellular and transcriptional responses in AD pathogenesis. Microglia are not only neuroprotective in neurospheres but also act as key drivers of Aβ-dependent *APOE* expression suggesting critical roles for microglia in regulating *APOE* in the AD brain. This novel, well characterized, functional *in vitro* platform offers unique opportunities to study the roles and responses of microglia to Aβ modelling key aspects of human AD. This tool will help identify new therapeutic targets, accelerating the transition from discovery to clinical applications.

**Highlights:**

- Well-characterized functional human iPSC-derived 3D neurospheres (hiNS) consisting of neurons and astrocytes can be supplemented with microglia/macrophages (hiMG)
- Chronic amyloidosis in the presence of hiMG recapitulate key features and gene expression profiles of AD
- hiMG within the model phagocytose Aβ and mitigate Aβ-induced neurotoxicity, reducing oxidative stress and neuronal damage
- hiMG are essential for Aβ to upregulate AD-like gene expression signatures in astrocytes
- Immunohistochemical analysis reveals hiMG-dependent colocalization of Aβ and APOE

## Introduction

Alzheimer’s disease (AD) remains inextricably linked to progressive aggregation of Aβ and Tau in the brain. Both are prime targets for therapeutic interventions, with two monoclonal antibodies against Aβ recently approved by the US Food and Drug Administration for use in AD. The mechanisms by which Aβ and Tau cause neurodegeneration are not fully understood. Aβ plaques initiate a significant neuroinflammatory reaction including robust microglial activation. However, the detrimental versus neuroprotective effects of microglia activation remains controversial. For example, a mutation in *TREM2*, which reduces the efficiency of microglial Aβ phagocytosis confers significant risk for developing sporadic AD, suggesting an important protective role of microglia[1]. Microglial phagocytosis is also the principal mechanism by which monoclonal antibodies against Aβ, such as lecanemab and donanemab, remove brain Aβ leading to improvement in clinical function and AD biomarkers[2, 3]. However, preclinical models have shown that microglial activation can also lead to excessive synaptic pruning which can accelerate neurodegeneration[4]. While there is broad agreement on the potential importance of resolving how glia contribute to sporadic AD pathophysiology[5–7], finding the most relevant preclinical system to best model AD pathophysiology remains a significant challenge. Murine models of AD have been used to investigate cellular mechanisms in AD for several decades, but clinical trials based on results from these models have rarely been successful[8]. The establishment of cerebral organoid and neurosphere 3D human cell culture systems sparked widespread interest in developing 3D cell culture models that more faithfully resemble human AD, to achieve better translation of pre-clinical studies to novel therapies[9–12]. Most of these approaches rely on the expression of familial AD mutations in *APP* or *PSEN1*[13–16]. Mimicking the AD-associated transcriptional changes of glial cells in a human 3D sporadic AD model system could help to unravel the impact of neuron-glia-Aβ interactions in AD.

Here we describe the utility of using human induced pluripotent stem cell (hiPSC)-derived 3D neurospheres (hiNS) to determine the impact of chronic Aβ exposure, a characteristic feature of AD, on neuronal and glial function. Introducing hiPSC-derived microglia (hiMG) into hiNS during chronic Aβ treatment triggered striking neuroprotection, by preventing Aβ neuronal activity, oxidative stress and reducing neuronal death. While hiMG displayed a remarkable phagocytic activity towards Aβ, single nuclei RNA sequencing revealed that the addition of hiMG also had significant effects on astrocyte and neuronal transcriptional profiles. For example, many AD-associated genes of interest in astrocytes, such as *APOE*, *CLU*, *LRP1* and *VIM*, were upregulated by Aβ only in the presence of hiMG. This study sheds light on the critical timing and multifaceted role of microglia in neuroprotection against Aβ, offering new insights into potential therapeutic strategies for AD.

## Results

### Formation of hiPSC-derived neurospheres as a 3D tissue culture system includes neurons, astrocytes and microglia

Blood-derived hiPSCs were generated from a healthy individual and subsequently differentiated into neural progenitor cells (NPCs) (see Suppl. Table 1 for media compositions). NPCs were plated in an AggreWell™ 800 plate to initiate the formation of hiNS. After 7 days *in vitro* (DIV), hiNS were transferred to 10 cm petri dishes on an orbital shaker for subsequent differentiation and maturation before plating into 48-well plates. While NPCs spontaneously differentiated into neurons and astrocytes, microglia did not differentiate in the 3D neurospheres. Therefore, hiMG were cultured in parallel to hiNS and added to a subset of mature neurospheres with neurons and astrocytes/NPCs to determine the impact of microglia. Neurospheres without added hiMG are termed hiNS(-) whereas neurospheres with hiMG infiltration are termed hiNS(+) (Figure 1A). hiNS were readily transduced using adeno-associated viruses (AAVs) encoding cell type specific expression with the neuronal-selective promoter *hSyn* or the astrocyte-selective promoter *gfaABC1D*. In contrast to certain 3D cerebral organoid models [17], our hiNS did not develop necrotic regions, likely because they remain a smaller size, rarely exceeding a diameter of 500 μm. To monitor tissue health of hiNS, we transduced neurospheres with *hSyn*-roGFP1, a redox-sensitive green fluorescent protein that detects cellular oxidative stress associated with reactive oxygen species production and, typically, necrosis[18, 19]. We found that neurons in hiNS displayed a largely reduced cytosol throughout the neurosphere, confirming the absence of necrotic tissue (N = 4 each, Figure 1B).

**Figure 1:**
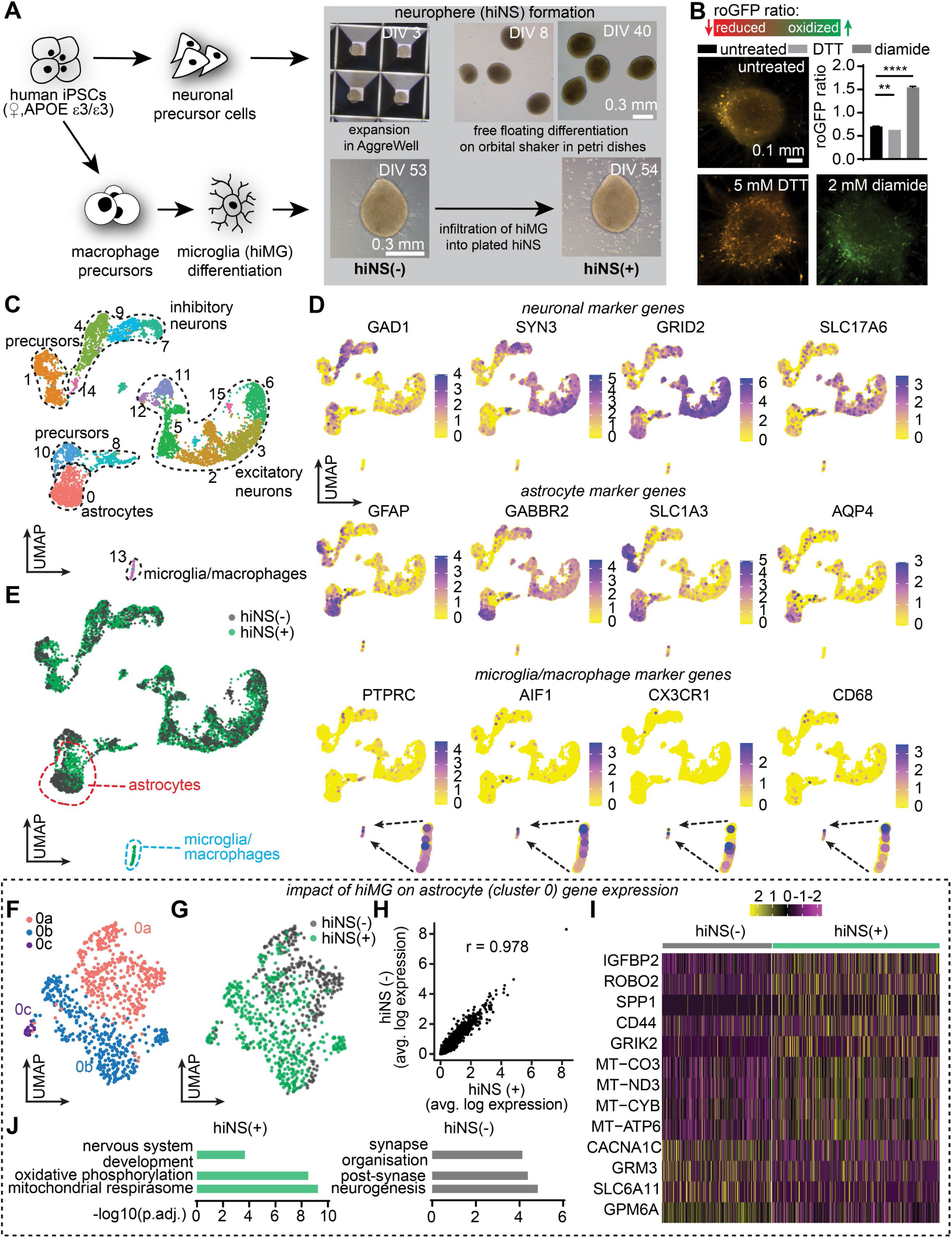
Human iPSC-derived mixed cell type neurospheres (hiNS) are comprised of neurons and glia. **A**: Schematic of hiNS generation and infiltration of human iPSC-derived microglia (hiMG) to form mature hiNS. hiNS after hiMG infiltration are defined as hiNS(+) while labelled hiNS(-) in the absence of hiMG. **B**: Redox imaging of hiNS(-) following transduction with AAV.php.eb.hSyn-roGFP1. A redox ratio is quantified by dividing green emission acquired at 405 nm by green emission acquired at 488 nm. As controls, DTT (5 mM) was used to fully reduce neurospheres and diamide (2 mM) to fully oxidize the tissue. Resting redox states of hiNS(-) (N=6) were only slightly higher than that of fully reduced DTT -treated hiNS (N=5) indicating healthy neurons throughout the 3D tissue. Full oxidation by diamide (N=4) indicates maximum roGFP ratios. Statistical testing performed using one-way ANOVA followed by Holm-Sidak post-hoc test. **C**: Single nuclei RNA sequencing (snRNA-seq) on hiNS(+/-). Overview of identified cell clusters and cell type identification in a merged data set of both groups **D**: Expression pattern of genes across cell populations for neuronal, astrocytic and microglia/macrophage displaying example marker genes. **E**: Datasets of origin confirming the presence of microglia/macrophage (marked in blue) is exclusive to hiNS(+) and that astrocyte gene expression (marked in red) appears shifted by the presence of hiMG. **F**: UMAP of isolated astrocyte populations from both data sets reveals 3 distinct sub-clusters with clusters 0b and 0c predominantly in hiNS(+) astrocytes and cluster 0a preferentially in hiNS(-) astrocytes. **G**: Analysis of the dataset of origin illustrating the shift in astrocyte gene expression in the presence of microglia. **H**: Pearson correlation (r = 0.978) confirms the close similarities between the two astrocyte populations. **I**: Differential gene expression (DEGs) pattern of a selection of DEGs shown in heatmaps. **J**: Gene ontology analysis of astrocyte clusters with and without hiMG.

To confirm the identity of cell types in hiNS and their patterns of gene expression, we performed single nuclei RNA sequencing (snRNA-seq). Specifically, we cultured hiNS for 50 days, and hiMG were added and allowed to infiltrate into the tissue for another 10 days. hiNS(+) or hiNS(-) were collected and flash frozen for subsequent snRNA-seq analysis. Single cell nuclei were isolated and sequenced using the 10X Genomics platform, and transcriptome analysis was performed using our previously described pipeline [20–24]. After filtering the datasets to remove low quality cells with few expressed genes or high mitochondrial proportions and cell doublets, we obtained 4750 and 1892 single cell transcriptomes for the hiNS(+) and hiNS(-), respectively. We then merged the datasets to allow a direct comparison of transcriptomes in the two conditions. Genes with high variance were used to compute principal components as inputs for projecting cells in two-dimensions using t-distributed stochastic neighbor embedding (t-SNE) and clustering, performed using a shared nearest neighbors-cliq (SNN-cliq)-inspired approach built into the Seurat R package at a range of resolutions.

Uniform Manifold Approximation and Projection for Dimension Reduction (UMAP) visualization and analysis of well-defined marker genes identified 15 clusters containing the predicted cell types (Figure 1C,D). Cell clusters with similar gene expressions were grouped together in a two-dimensional UMAP. These included astrocytes expressing *GFAP, AQP4, SLC1A3* and *GABBR2* (cluster 0), and two types of neurons in various stages of maturation: inhibitory interneurons (clusters 4, 7, 9 and 14) expressing *GAD1, GAD2* and *SYN3*, and excitatory neurons (clusters 2, 3, 5, 6, 11, 12 and 15) enriched for *GRID2, GRM7, SLC17A6* and *SYN3*. The dataset also included neural precursor cells (clusters 1, 8, 10 and 14), and microglia/macrophages expressing *PTPRC, AIF1, CX3CR1* and *CD68* (cluster 13). Note that *GFAP* expression was not exclusive to astrocytes but also highly expressed in some NPC populations.

Analysis of the dataset of origin (Figure 1E) confirmed that the microglial cluster included cells only from the hiNS(+) dataset. The other clusters included intermingled cells from both conditions, except for cluster 0 astrocytes, where many of the cells appeared transcriptionally distinct. To further explore this finding, we subsetted the astrocytes and reanalyzed them separately. This analysis (Figure 1F,G) defined two large clusters and one smaller cluster. Two of the clusters (0a and 0c) included astrocytes from both conditions, while the other large cluster (0b) was almost completely comprised of hiNS(+) astrocytes. Pearson correlation analysis of average gene expression confirmed that, as suggested by the clustering analysis, hiNS(+) astrocytes were similar but not identical to the hiNS(-) astrocytes, with r = 0.978 (Figure 1H).

To better understand these similarities and differences, we performed differential gene expression (DEG) analysis. There were 400 DEGs when comparing the two conditions, with 44 enriched in hiNS(+) astrocytes and 356 in hiNS(-) astrocytes (> 1.3-fold average change, p adj < 0.05; Suppl. Tables 2 and 3). Notably, DEG analysis coupled with gene ontology (Figure 1I,J; Suppl. Table 4) demonstrated that hiNS(+) astrocytes were highly enriched for genes and gene categories associated with oxidative phosphorylation such as *MT-CO3, MT-ND3*, *MT-ATP6,* and *MT-CYB*, suggesting they were more metabolically active. By contrast, hiNS(-) astrocytes gene ontology analysis identified significant gene categories including neurogenesis, post-synapse and synapse organisation (Figure 1J, Suppl. Table 5).

### hiNS are highly functional 3D human tissue cultures

The presence of the various differentiated major cell populations identified in our snRNA-seq analysis were confirmed using immunofluorescence staining for neurons (MAP2), astrocytes/NPCs (GFAP), and microglia/macrophages (IBA1) in fixed hiNS(+) at DIV 60. Neurons and astrocytes with extensive processes were observed throughout the neurospheres in addition to hiMG (Figure 2A). Imaging intracellular calcium concentration transients was used for tracking neuronal activity in hiNS monitored by imaging *hSyn*-GCaMP6f or *hSyn*-jRGECO1 expression for live imaging of neuronal activity versus imaging GCaMP8s in astrocytes induced by *gfaABC1D*-GCaMP8s expression. hiNS(-) displayed frequent synchronized and propagating calcium transients. Co-transduction of *hSyn*-jRGECO1 with *gfaABC1D*-GCaMP8s allowed us to image intracellular calcium of neuronal and astrocytic cell populations simultaneously and revealed that calcium elevations rise in tandem in both populations (Figure 2B). The synchronized large [Ca^2+^]_i_ transients traversed through hiNS in a wave-like pattern reminiscence of large calcium waves (Figure 2C) observed during neurodevelopment[25, 26].

**Figure 2:**
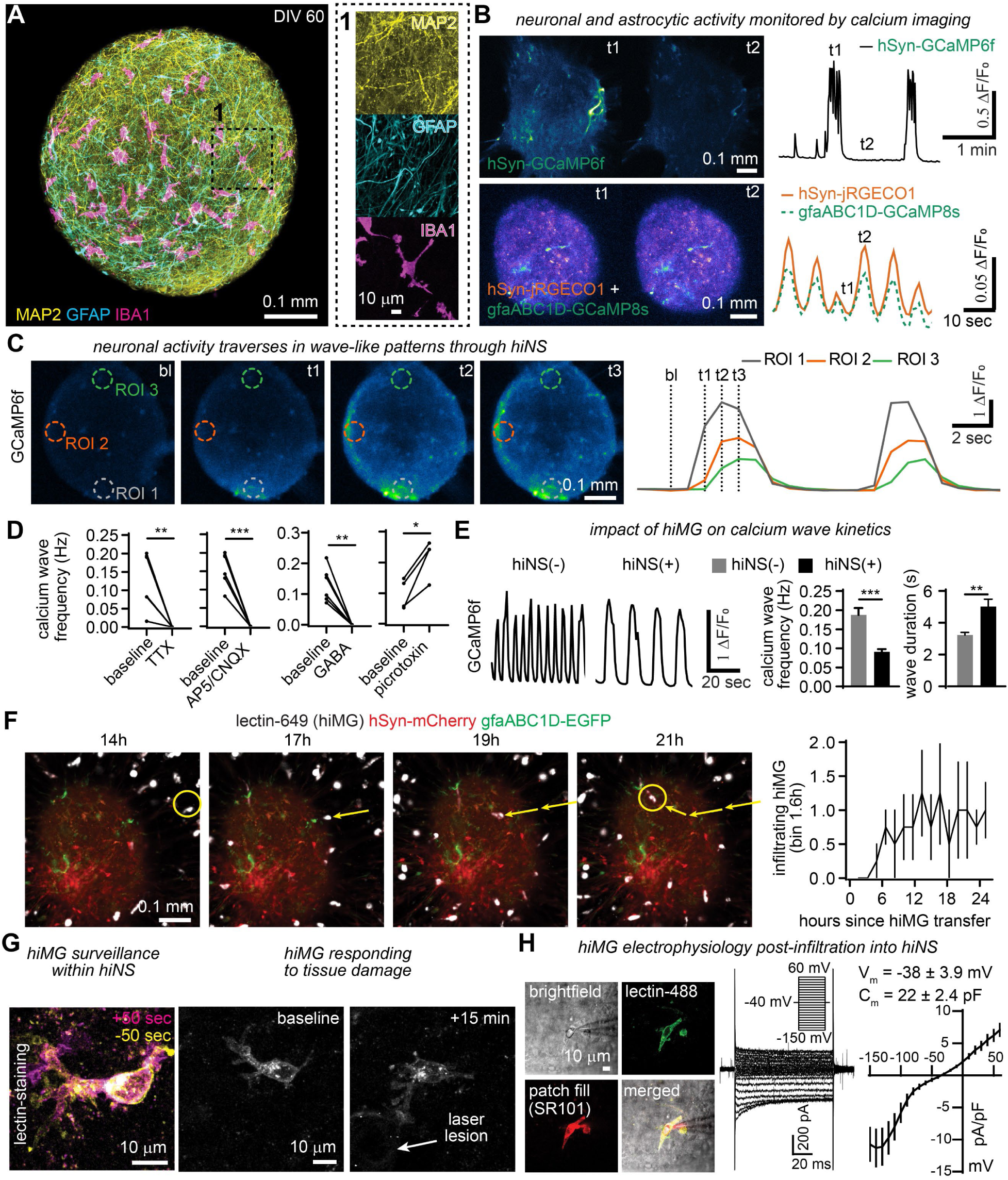
Neurons and glia are highly functional in hiNS mimicking early brain physiology. **A**: Immunofluorescence of MAP2, GFAP and IBA1 in a mature neurosphere at day in vitro (DIV) 60. **B**: Functional intracellular neuronal calcium (AAV.php.eb.-*hSyn*-GCaMP6f or AAV2/9.-*hSyn*-jRGECO1) and astrocytic calcium (AAV2/9-*gfaABC1D*-GCaMP8s) imaging shows highly synchronized and simultaneous calcium activity in both neurons and astrocytes in hiNS(-). **C**: Spontaneous neuronal calcium activity traverses through hiNS in a wave-like manner. **D**: Pharmacological block of calcium waves by TTX (1 μM) (N=6), AP5 (100 μM)/CNQX (20 μM) (N=6) and GABA (300 μM) (N=6). Application of picrotoxin (100 μM) (N=4) resulted in a significant increase in calcium wave frequencies. Statistical testing performed using paired student t-tests for each drug application vs its baseline. **E**: Infiltration of hiMG in hiNS(+) results in a significant reduction in calcium wave frequencies with longer wave durations (hiNS(-): N=10; hiNS(+): N=12). Statistical testing performed using unpaired student t-tests. **F**: Long-term confocal imaging of transduced hiNS(+) (N=4) with AAV.php.eb-hSyn-mCherry (red) and AAV.php.eb-gfaABC1D-EGFP (green) and tomato lectin DyLight 649 stained hiMG. hiMG infiltration into the tissue was quantified within the first 24h post-transfer (see yellow arrows and areas). **G**: hiMG within hiNS display surveillance behaviour with rapidly moving processes and a more stationary soma. Tissue damage induced by a laser lesion leads to rapid process outgrowth of microglia towards the lesion. **H**: Electrophysiological properties of hiMG 7-10 days post infiltration (N=9).

Astrocytic uptake of extracellular glutamate is known to regulate neuronal calcium activity[27, 28]. To further characterize the functional properties of hiNS we induced *hSyn*-iGluSnFr expression in neurons to monitor extracellular glutamate release during calcium wave propagations and found that the rise of extracellular glutamate was highly synchronized with large propagating [Ca^2+^]_i_ waves (Figure S1A). Our snRNA-seq data revealed widespread expression of the glutamate transporters EAAT1 (*SLC1A3*) and EAAT2 (*SLC1A2*), with high levels of expression in astrocytes (Figure S1B). Pharmacological block of EAAT1/2 with TFB-TBOA lead to a steady rise in extracellular glutamate levels which eventually inhibited wave formation (Figure S1C). We performed additional pharmacological experiments to characterize the nature of calcium waves in hiNS. [Ca^2+^]_i_ waves were suppressed by either the Na^+^ channel blocker TTX, or ionotropic glutamate receptor antagonists AP5/CNQX, or by applying GABA. Picrotoxin application, which antagonizes GABA receptors and chloride channels, significantly increased calcium wave frequencies (Figure 2D).

We then tested whether infiltrating hiMG alter calcium wave frequencies by comparing GCaMP6f recordings from hiNS(-) (N = 10) with hiNS(+) (N = 12). The presence of hiMG reduced the calcium wave frequency by half, from 0.19 to 0.09 Hz (*** p<0.001) and increased wave duration from 3.2 to 5.0 sec (** p<0.01), indicating that hiMG leads to altered neuronal activity or neurodevelopment (Figure 2E).

We next investigated the infiltration behaviour and physiological properties of hiMG during and after tissue infiltration into hiNS. hiMGs were identified after fixation by IBA1 immunofluorescence or were live-stained using tomato lectin conjugated dyes (Figure S2). We performed 24h-continuous confocal live imaging on hiNS expressing *hSyn*-mCherry and *gfaABC1D*-EGFP to monitor live-stained hiMG infiltrating into hiNS (N = 4). hiMG were pre-stained with tomato lectin DyLight 649. Tracking hiMG movements in hiNS revealed infiltration occurring 6-12h after the transfer (see yellow arrows, Figure 2F). To test if hiMG that infiltrated within hiNS(+) exhibit physiological microglia-like properties, we again stained hiMG with tomato lectin (DyLight 488) 7-9 days post-infiltration into hiNS. hiNS(+) were imaged by 2-photon microscopy to monitor microglial process movement, damage responses and for patch-clamp recordings. We observed tissue surveillance by hiMG processes as well as process outgrowth towards a focal laser lesion mimicking *in vivo*-like behaviour (N=3, Figure 2G). Patch-clamp recordings on hiMG within hiNS(+) confirmed microglial identity, revealing average reversal potentials of -38 ± 3.9 mV and cell capacitances of 22 ± 2.4 pF (N = 9). A series of de- and hyperpolarizing voltage steps revealed inward rectifying potassium currents typical for microglia [29–31] (Figure 2H). We conclude that infiltrating hiMG display a robust microglial-like phenotype exhibiting several properties that resemble *in vivo* microglia.

### Chronic treatment of hiNS with oligomeric Aβ results in plaque-like aggregates and a series of neurotoxic effects

To model how the human brain responds to Aβ aggregation, we first tested if supplementing hiNS(-) media with Aβ42 results in the formation of insoluble, plaque-like aggregates in the tissue. We followed a well characterized[32–36] protocol for the formation of oligomeric Aβ42 (oAβ) and tested three different concentrations added during regular media changes for a total of 7 days (0.012, 0.048 and 0.24 μg/mL). Only the highest Aβ concentration resulted in substantial accumulation of insoluble Aβ in the tissue (Figure S3A,B). As an alternative strategy, we tested whether exposing hiNS(-) to fibrillar Aβ42 (fAβ) for 7 days would have the same effect as that seen with oAβ. However, fAβ caused highly aggregated Aβ deposits that appeared as abnormally large sheets spanning the tissue, and were much different from the plaque-like aggregates found after oAβ stimulation (Figure S3C). We consequently focused on using oAβ treatment instead of fAβ for tissue culture treatments.

We next tested the effects of chronic oligomeric Aβ42 (0.24 μg/mL) for up to 35 days, first in hiNS(-), in the absence of microglia/macrophage-like cells (Figure 3A). Large quantities of aggregated Aβ were detected with different Aβ antibodies within hiNS(-) (Figure 3B). Immunostaining for neurons and astrocytes/NPCs revealed degenerated or swollen processes and cell bodies at the end of the treatment period indicating severe toxicity (Figure 3C, see arrows). To monitor neuronal redox levels, hiNS(-) expressing *hSyn-*roGFP1 were imaged after treatment, revealing severe oxidative stress evident by a significant increase in roGFP1 redox ratios (Figure 3D). Continuous (up to 20h) live roGFP1 imaging in Aβ treated hiNS(-) revealed a significant increase in the frequency of oxidizing cells compared to hiNS(-) not exposed to Aβ, further confirming Aβ-induced oxidative stress (Figure S4). Neuronal activity was repeatedly monitored by imaging GCaMP6f in hiNS(-) as described above. A significant reduction in calcium wave frequencies was observed at the end of the third week and in the fourth week of Aβ exposure (Figure 3E).

**Figure 3:**
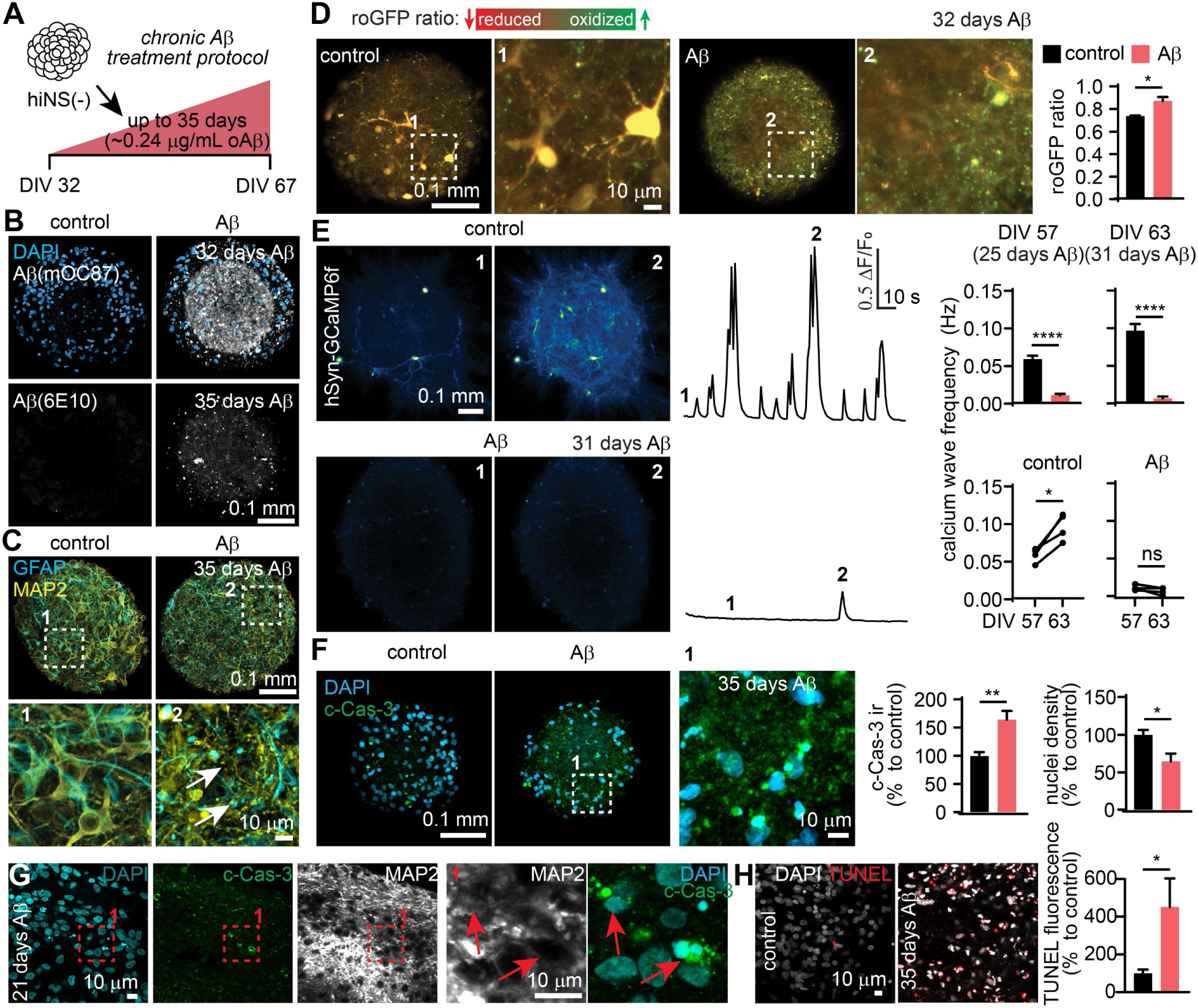
Chronic Aβ stimulation in hiNS(-) triggers a sequence of neurotoxic events and phenotypes in the absence of hiMG. **A**: hiNS(-) cell culture media was supplemented with ∼0.24 μg/mL oligomeric Aβ1−42 for up to 35 days. **B**: Treatment results in the formation of plaque-like Aβ aggregates in hiNS(-) which can be stained with different Aβ antibodies (here 6E10 and mOC87). **C**: Immunostaining for neurons (MAP2) and astrocytes/NPCs (GFAP) after 35 days of Aβ exposure reveals dystrophic appearing neuronal and astrocytic processes (see white arrows) **D**: roGFP1 based redox imaging in neurospheres reveals significant oxidation indicating oxidative stress in neurons following chronic Aβ treatment (N=4 each). **E**: Calcium wave frequencies (quantified by GCaMP6f live imaging) gradually decline with Aβ treatment resulting in dysfunctional neuronal activity (N=4 each). **F**: Immunostaining for cleaved Caspase-3 (c-Cas-3) indicates apoptosis in hiNS(-) after chronic Aβ exposure. Quantification of DAPI positive nuclei in hiNS(-) suggests neuronal death as the ultimate fate following 35 days of chronic Aβ treatment. **G**: Co-staining for c-Cas-3/MAP2 indicates neurons undergoing apoptosis and loss of nuclei density (N=4 each). **H**: Apoptosis after 35 days of Aβ exposure was further confirmed by a significant increase in TUNEL fluorescence (control: N=4; Aβ: N=5). All statistical testing was performed using unpaired student’s t-tests.

We then investigated whether chronic Aβ treatment ultimately results in neuronal loss. Immunostaining for cleaved Caspase-3 (c-Cas-3), a key mediator of neuronal apoptosis, indicated a ∼50% increase in apoptotic cells after Aβ treatment in hiNS(-) as well as a significant reduction of nuclei density within the tissue, suggesting cell death (Figure 3F). Neuronal apoptosis during Aβ exposure was observed by co-staining for c-Cas-3 and the neuronal marker MAP2 (Figure 3G). To confirm the increase in apoptotic cell death, we performed terminal deoxynucleotidyl transferase dUTP nick end labeling (TUNEL) to detect fragmented DNA. There was a significant increase in TUNEL fluorescence confirming apoptosis during chronic Aβ exposure (Figure 3H).

In conclusion, our chronic Aβ stimulation protocol resulted in a sequence of neurotoxic effects including the formation of plaque-like Aβ aggregates within days of exposure, oxidative stress and dysfunctional neuronal activity 3-4 weeks into Aβ treatment, and ultimately Caspase-3 dependent apoptosis and neuronal loss 4-5 weeks after Aβ exposure.

### hiMG dependent phagocytosis and removal of plaque-like Aβ aggregates

The role of microglia in AD pathogenesis has been extensively studied, with both neurotoxic- and neuroprotective roles described[37, 38]. We designed two different chronic Aβ treatment protocols to elucidate the impact of microglia on neurotoxicity when added to hiNS at different time points following Aβ exposure. This included six experimental conditions: control (ctrl), 3-week (3w Aβ hiNS) or 5-week (5w Aβ hiNS) exposure each with (+) or without (-) hiMG. hiMG were added to the tissue for the last 10 days of the treatment protocol (Figure 4A).

**Figure 4:**
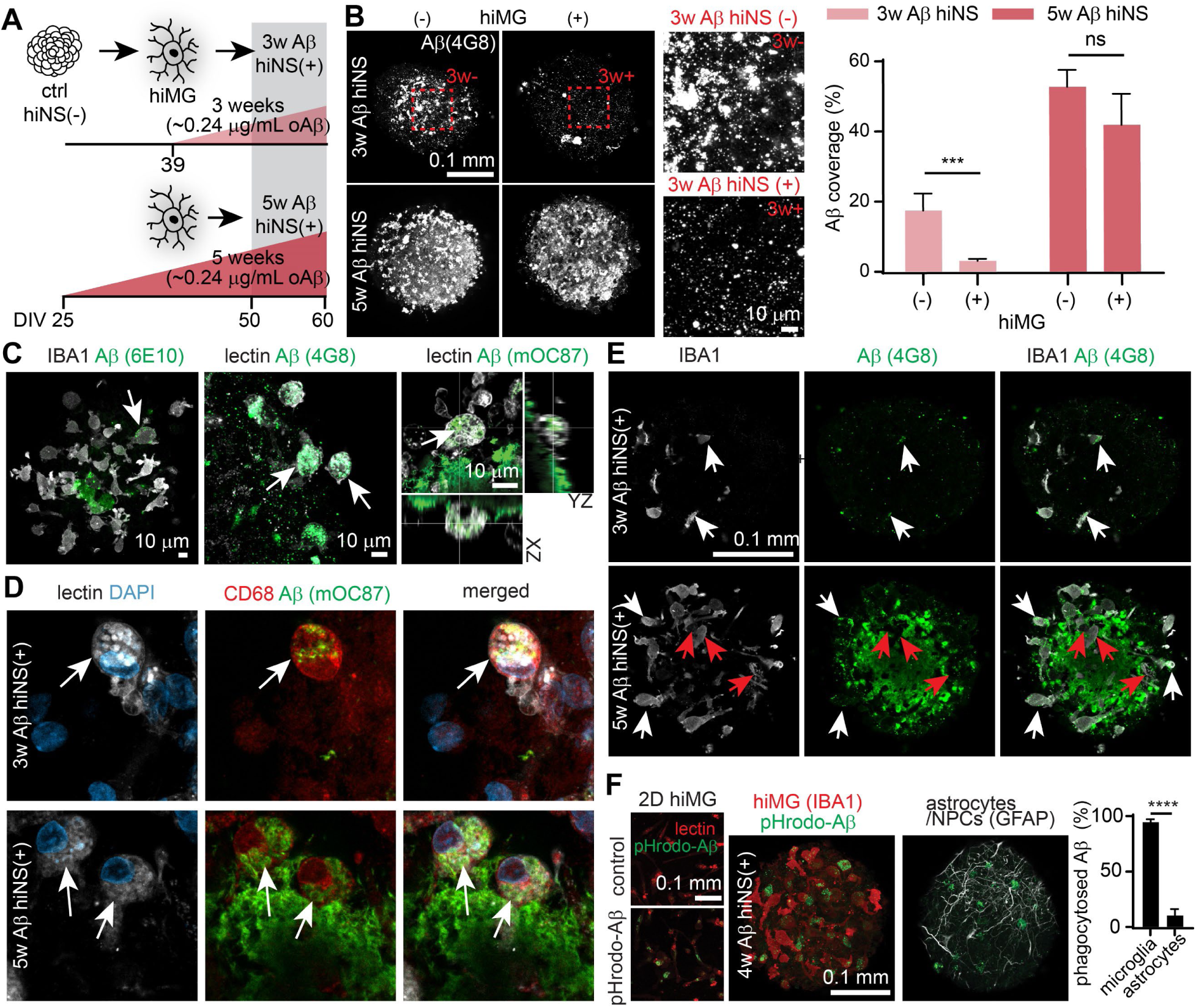
Timing-dependent neuroprotection by hiMG in hiNS(+) during two different chronic Aβ treatment protocols indicating hiMG are effective in phagocytosing Aβ. **A**: Schematic illustrating 3-weeks (3w Aβ hiNS(+)) and 5-weeks (5w Aβ hiNS(+)) long chronic Aβ protocols. In each, hiMG were transferred 10 days before the end of both protocols. **B**: Quantification of hiNS areas covered by Aβ (stained for 4G8) reveals less Aβ burden after 3w than 5w with a significant reduction in Aβ levels in 3w Aβ hiNS(+) compared to 3w Aβ hiNS(-) (N=6 each). Statistical testing performed using two-way ANOVA followed by Holm-Sidak post-hoc tests on logarithmically transformed data. **C**: Co-localization of Aβ in IBA1 or lectin-stained hiMG indicates phagocytosis of highly aggregated (mOC87 positive) Aβ. **D**: CD68 immunofluroescence confirms phagocytic microglia-like cells (white arrows) in both, 3w/5w Aβ treated hiNS(+). **E**: In both, 3w/5w Aβ treated hiNS(+), Aβ co-localizes with hiMG cell bodies (see white arrows). Only at 5w, hiMG appear to clear volumes of tissue from Aβ leaving dark shadows in the Aβ staining pattern (see red arrows). **F**: Conjugation of Aβ oligomers with pHrodo green (pHrodo-Aβ) confirms functional phagocytosis of Aβ in hiMG. Successful microglial uptake of pHrodo-Aβ was first tested in 2D hiMG (left). 3w Aβ hiNS(+) were treated for 1w with pHrodo-Aβ to confirm microglial Aβ phagocytosis in 3D. Additional co-staining for IBA1 and GFAP reveals that microglia are the main cell type phagocytosing Aβ in hiNS (right) (N=6 IBA1 co-stainings; N=3 GFAP co-stainings). Statistical testing was performed using an unpaired student’s t-test.

We first assessed the impact of microglia on Aβ aggregation. Aβ immunoreactivity was quantified using an anti-Aβ antibody (4G8) for all groups. The amount of Aβ was less in 3w Aβ hNS(-) compared to 5w Aβ (hNS(-) (Figure 4B). hiMG addition significantly reduced Aβ levels at 3w (∼17% Aβ coverage reduced to ∼3%, *** p < 0.001), but not at 5w, indicating that adding hiMG is more efficient at clearing Aβ aggregates during the shorter Aβ treatment protocol, with a shorter overall exposure to Aβ (Figure 4B). The more efficient removal of Aβ indicates that neuroprotective properties of hiMG could be more pronounced at 3w than at 5w Aβ exposure in hNS(+).

Next, we investigated whether hiMG phagocytosis of Aβ contributed to the observed reduction in Aβ deposits. Co-staining microglia (lectin or IBA1) and Aβ (6E10, 4G8 or mOC87) revealed that Aβ was readily found in the cytosol of infiltrated hiMG in Aβ-treated hiNS(+) (Figure 4C). The strongest intracellular Aβ signal was seen by staining specifically for fibrillar Aβ (mOC87[39]), while the Aβ antibody 6E10 revealed only a faint signal inside hiMG, indicating that Aβ inside hiMG appears mostly in a highly aggregated form. Immunostaining for CD68 confirmed that phagocytosing hiMG in both 3w and 5w Aβ treated hiNS(+) express CD68, a well known phagocytosis marker[40]. Intracellular Aβ was identified after both 3w and 5w Aβ treatment (Figure 4D, white arrows). While we did not find hiMG to significantly reduce overall extracellular Aβ burden after 5w, they nevertheless appear to have cleared significant amounts of Aβ, leaving Aβ−cleared tissue in their path (Figure 4E, red arrows). Successful phagocytosis requires the uptake of Aβ into acidic phagocytic compartments. To test that, we conjugated our oligomeric Aβ with pHrodo green which is fluorescent at acidic pH[41]. hiNS were treated with regular oligomeric Aβ for 3w followed by pHrodo-Aβ for an additional 7 days (4w Aβ total). Aβ treatment was then stopped prior to hiMG addition. hiNS were fixed 8 days later and imaged to quantify intracellular pHrodo-Aβ levels in the tissue. We found a strong uptake of pHrodo-Aβ inside IBA1 positive hiMG, whereas only small amounts overlapped with GFAP-positive astrocytes/NPCs, confirming that the main cell types in hiNS(+) facilitating Aβ phagocytosis are microglia/macrophage-like hiMG (Figure 4F).

### Time-dependent neuroprotective properties of hiMG

We next sought to elucidate neuroprotective properties of hiMG in response to chronic Aβ. hiNS were immunostained for neurons (MAP2), astrocytes/NPCs (GFAP), and microglia (IBA1) markers in all six experimental conditions. First, we quantified the number of hiMG inside hiNS(+) and found that the number of hiMG within 5w Aβ hiNS(+) increased by more than two-fold compared to ctrl hiNS(+), whereas the number of microglia in 3w Aβ hiNS(+) did not increase. Additionally, we found that GFAP immunofluorescence increased significantly in the presence of hiMG at 3w and 5w, indicating microglial-dependent astrocyte reactivity by Aβ (Figure 5A). We repeated roGFP1 redox imaging following viral transduction using our *hSyn*-roGFP1 construct. Live confocal microscopy, performed every few days, allowed us to follow neuronal redox states during the chronic Aβ treatment with or without hiMG present. Significant Aβ induced oxidative stress was detected at the end of the 3w Aβ treatment window at DIV59, evident by the increased roGFP ratio in 3w Aβ hiNS(-)(****, p<0.0001) (Figure 5B, white arrows). These changes appeared only at DIV 59, and not 5 days prior at DIV 54 (Figure 5B). Interestingly, oxidative stress was completely prevented by the presence of hiMG in 3w Aβ hiNS(+), suggesting robust neuroprotective functions of hiMG (Figure 5B). However, Aβ induced oxidative stress at 5w was severe and not significantly reduced after hiMG infiltration (Figure 5B, bottom part). These data suggest that hiMG are able to reduce Aβ-induced oxidative stress if added early enough during chronic Aβ treatment, but fail to recover or reverse severe oxidative stress late during Aβ treatment.

**Figure 5:**
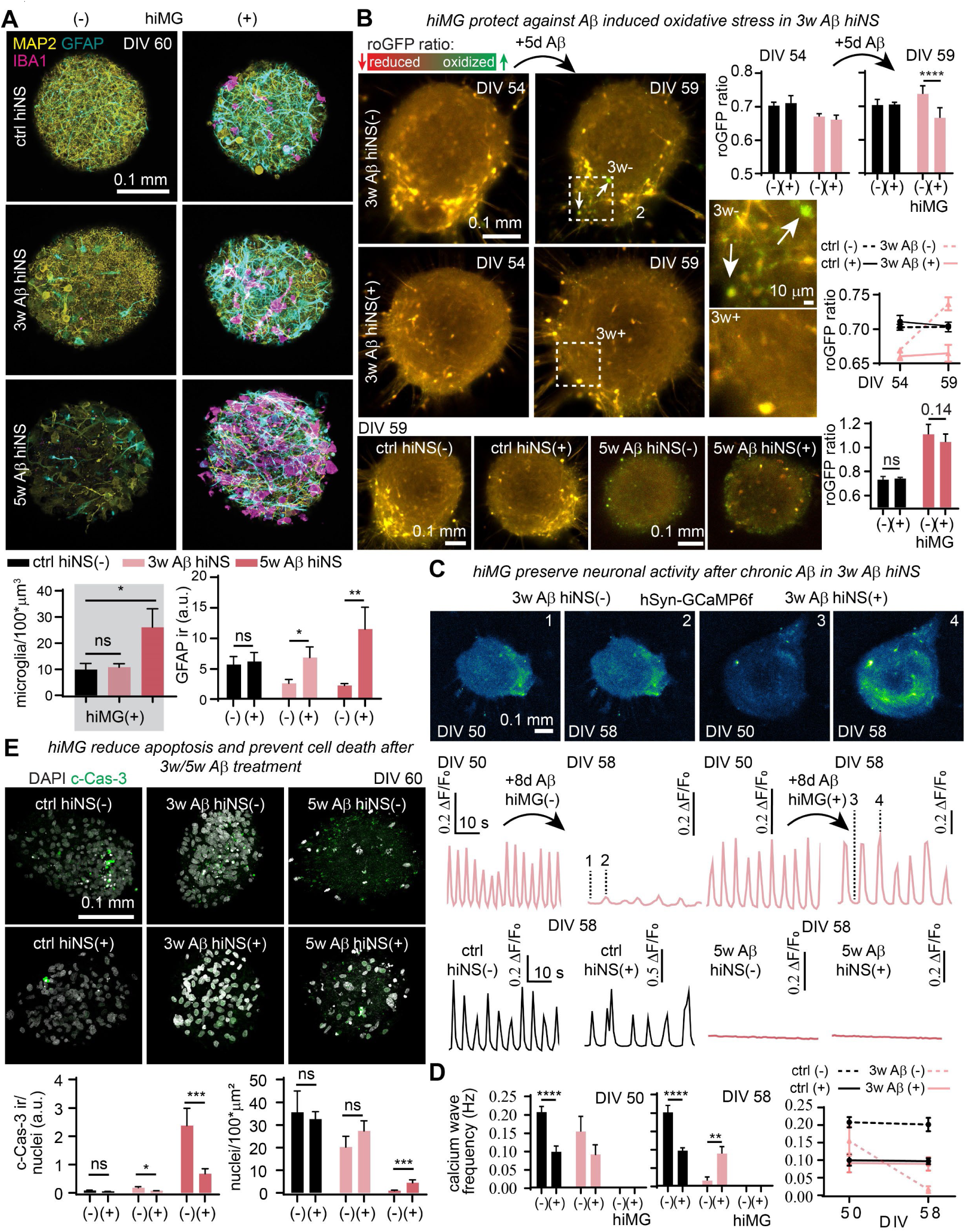
Functional neuroprotection by hiMG during chronic amyloidosis is limited to 3w Aβ hiNS(+). **A**: Immunofluorescence staining for MAP2, GFAP and IBA1 reveals a larger hiMG population in 5w Aβ hiNS(+) compared to 3w Aβ hiNS(+) or controls (ctrl hiNS(+)) as well as an increase in GFAP immunoreactivity in 3w/5w Aβ hiNS(+) (N=6 each). **B**: Redox imaging using roGFP1 indicates mild oxidative stress in 3w Aβ hiNS(-) with a full rescue in 3w Aβ hiNS(+). Severe oxidative stress in 5w Aβ hiNS(-) was not ameliorated by hiMG (N=6 each). **C**: Functional recovery in neuronal calcium wave activity by hiMG in 3w Aβ hiNS(+) quantified using GCaMP6f. Calcium waves in 5w Aβ hiNS were completely absent and hiMG did not result in a functional recovery (ctrl hiNS(-): N=10; ctrl hiNS(+): N=12; 3w/5w Aβ hiNS(-/+): N=6 each). **D**: Quantification of calcium wave frequencies. **E**: At both time points, hiMG resulted in a reduction in c-Cas-3 immunoreactivity and ameliorated cell death (ctrl hiNS(-): N=9; ctrl hiNS(+): N=11; 3w Aβ hiNS(-/+): N=10 each; 5w Aβ hiNS(-): N=8; 5w Aβ hiNS(+): N=11). Statistical testing performed using two-way ANOVA followed by Holm-Sidak post-hoc tests on logarithmically transformed data, except for a one-way ANOVA followed Holm-Sidak post-hoc test for microglia quantification.

Similar to roGFP1 live imaging, we monitored neuronal activity using GCaMP6f and found that calcium wave frequencies were reduced at the end of the 3w Aβ treatment window in 3w Aβ hiNS(-) (**, p<0.01) at DIV 58 while wave frequencies were unchanged by Aβ at DIV50 (Figure 5C,D). Matching the neuroprotective properties of hiMG, ameliorating oxidative stress in 3w Aβ hiNS(+), we found that hiMG fully rescued neuronal activity in 3w Aβ hiNS(+), confirming their neuroprotective ability preventing the decline of neural activity. At 5w Aβ, calcium waves were completely absent without ameliorating effects of hiMG, suggesting that functional recovery is only possible if hiMG are present early during chronic Aβ treatment (Figure 5D).

The functional recovery by hiMG in hiNS exposed to chronic Aβ correlates with their high capacity to remove plaque-like aggregates in 3w Aβ hiNS(+) (Figure 4B) suggesting a potential link between Aβ phagocytosis and neuroprotection by hiMG.

We then determined whether hiMG prevented neuronal death by staining for c-Cas-3 and quantifying nuclei in hiNS. While we found only a moderate increase in c-Cas-3 immunoreactivity at 3w, it was significantly reduced in 3w Aβ hiNS(+) (*, p<0.05) with an even larger difference at 5w Aβ hiNS(-)(***, p<0.001). Note that hiMG were identified by lectin staining, and their nuclei count was subtracted from whole-tissue nuclei quantification. Cell density in 5w Aβ hiNS(-) was dramatically reduced and partially, but highly significantly rescued in 5w Aβ hiNS(+) (***, p<0.001) (Figure 5E).

Aβ aggregation and accumulation in the AD brain are believed to precede and trigger subsequent Tau hyperphosphorylation[42]. A well characterized phosphorylation site is threonine 181 (pTau181) which we immunostained for in tissue from all six experimental groups (Figure S5A), together with a total Tau co-staining. The ratio of pTau181/total Tau was calculated to test whether Aβ or microglia impact Tau hyperphosphorylation in our model. While we detected small but significant changes in ratios due to the presence of hiMG, we did not detect any significant increase due to Aβ with or without hiMG present (Figure S5B). The lack of Aβ induced Tau hyperphosphorylation in our model was further confirmed by ELISA on supernatant of hiNS (Figure S5C) suggesting that the severe Aβ-induced pathology we observed does not result in Tau pathology in our embryonic model system.

Taken together, these data indicate that the neuroprotective potential of hiMG during chronic Aβ exposure is highly time-dependent: At 3w Aβ, added hiMG preserved neuronal function, preventing oxidative stress, and reducing Caspase-3 cleavage, while at 5w Aβ, hiMG did not rescue neuronal function or reduce oxidative stress but hiMG did significantly reduce apoptosis and loss of cell nuclei, demonstrating limited neuroprotection by hiMG at late stages of amyloidosis. Visual inspection revealed hiMG attached to and accumulated around hiNS prior to infiltration, and 5w Aβ treated hiNS displayed morphological changes indicating tissue degeneration (Figure S6) matching our immunofluorescence staining which revealed substantial tissue deterioration in 5w Aβ hiNS (Figure 5A).

We further investigated the electrophysiological properties of hiMG in Aβ treated hiNS. Membrane capacitances were significantly increased at 5w Aβ (Figure S7A-C), but no other changes were significant.

### hiMG alter astrocyte gene expression profiles in hiNS in response to Aβ

To elucidate potential mechanisms underlying the protective effects of microglia in hiNS chronically exposed to Aβ, we performed snRNA-seq on all six hiNS culture conditions (ctrl, 3w Aβ, 5w Aβ with or without hiMG). We isolated and sequenced single cell nuclei using the 10X Genomics platform and put the resultant transcriptomes through our pipeline[20–24]. After filtering the datasets for low quality cells, we merged these transcriptomes with those of the previously analyzed control hiNS (as in Figure. 1C,E).

UMAP visualization and analysis of the datasets of origin showed that the datasets merged well (Figure 6A,B). Marker gene analysis defined all of the same cell types that were seen in the ctrl hiNS cultures including neural and glial precursor cells, astrocytes, inhibitory and excitatory neuron lineages, and microglia/macrophages. However, analysis of the datasets of origin identified differences in cellular composition that were most obvious in the 5w Aβ hiNS. Specifically, in the 5w Aβ hiNS(-), the only surviving cells were astrocytes and inhibitory neurons, while excitatory neurons were almost entirely absent which is consistent with the dramatic reduction of nuclei numbers in this group (Figure 5E). This observation corroborates the known finding that excitatory neurons are more vulnerable in AD than inhibitory neurons[43, 44]. The 5w Aβ hiNS(+) contained more excitatory and inhibitory neurons, but the overall cell density was reduced relative to the 3w Aβ cultures and ctrl groups (Figure 6B). In addition, relative to the ctrl and 3w Aβ hiNS (+), there were proportionately more hiMG in 5w Aβ hiNS(+), likely reflecting increased proliferation or infiltration of hiMG matching our quantification of IBA1 positive microglia/macrophage-like cells (Figure 5A). By contrast, the distribution of cell types in the 3w Aβ hiNS cultures were similar to that seen in the ctrl groups.

**Figure 6:**
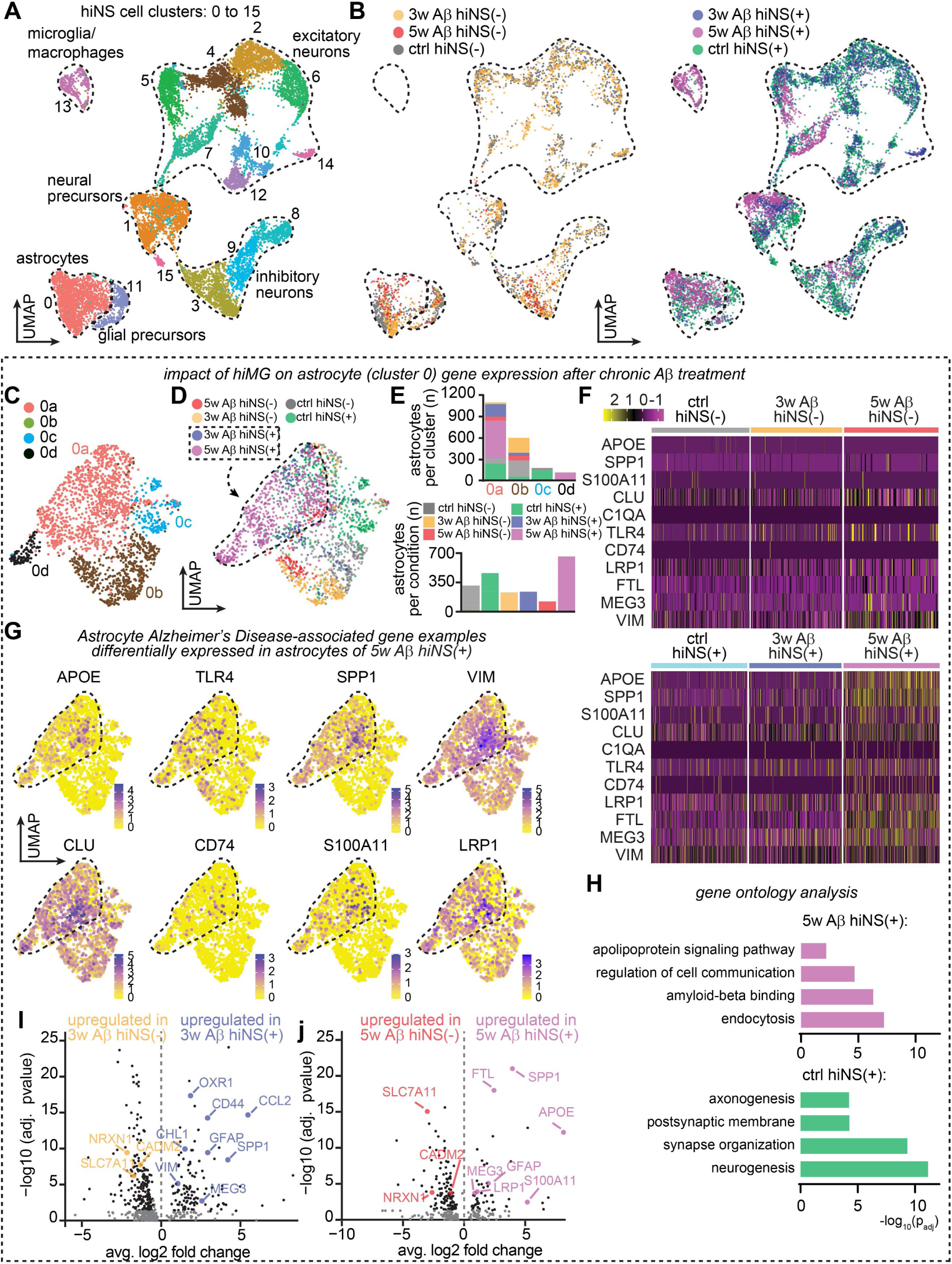
snRNA-seq on hiNS(-/+) in ctrl, 3w and 5w Aβ conditions reveals a hiMG-dependent astrocyte AD-like transcriptional profile in 3w/5w Aβ hiNS(+). **A**: Merged UMAP of all 6 experimental groups displaying cell clusters and cell type identification. **B**: Split merged UMAPs without hiMG (left) and with hiMG (right). Note the lack of a microglial population in hiNS(-), reduced neuronal populations in the absence of microglia indicating severe cell loss after 5w Aβ treatment. **C**: Isolated astrocyte population in a merged UMAP displaying four distinct clusters. **D**: Data set of origin for all 6 conditions illustrating significant shifts of astrocyte gene expression depending on Aβ treatment and hiMG presence. Dashed lines mark the rough outline of the predominantly 3w/5w Aβ hiNS(+) astrocyte populations. **E**: Number of astrocytes sorted per sub-cluster (top panel) and sorted by condition (bottom panel). Note that astrocyte numbers varied between conditions with 5w Aβ hiNS(+) marking the largest astrocyte population. **F**: Heatmap of differentially expressed genes selected from direct comparison between 5w Aβ hiNS(+) and ctrl hiNS(+) astrocytes. Note that upregulation of genes with Aβ treatment required the presence of hiMG, including *APOE*. **G**: Overlay examples of AD-associated gene expression in astrocytes in 5w Aβ hiNS(+) astrocytes. **H**: Gene ontology analysis of ctrl hiNS(+) and 5w Aβ hiNS(+) astrocyte groups highlight distinct hiMG-dependent phenotypical differences in Aβ treated astrocytes. **I**: Direct comparison of DEGs depending on hiMG in 3w (left) and 5w (right) Aβ treated hiNS shown in volcano plots (compared in pairs). Genes of interest are labelled. Note that many AD-associated genes, such as *GFAP*, *VIM* and *APOE*, are only upregulated if hiMG were present during Aβ treatment.

We first compared astrocytes in these different culture conditions by subsetting and reanalyzing transcriptomes from the astrocyte cluster (cluster 0). UMAP cluster and dataset of origin visualizations (Figure 6C,D) identified four clusters containing variable proportions of astrocytes from the different culture conditions. In particular, most astrocytes that came from cultures without added microglia were localized to cluster 0b regardless of whether they were exposed to control peptide or Aβ. Conversely, most astrocytes from cultures with hiMG were localized to cluster 0a. The exceptions to this intermingling were cluster 0d, which was almost completely comprised of transcriptomes from the 5w Aβ hiNS(+) cultures, and cluster 0c, which was mostly comprised of astrocytes from the control hiNS(+) cultures (Figure 6E).

These data suggest that astrocytes in the different cultures were transcriptionally distinct. We therefore assessed how Aβ altered astrocytes in the presence of hiMG by performing DEG analysis comparing the 5w Aβ hiNS(+) cells with ctrl hiNS(+). This analysis identified 288 DEGs when comparing the two conditions, with 209 enriched in 5w Aβ hiNS+ cells and 159 in ctrl hiNS(+) without Aβ (> 1.3-fold average change, p adj < 0.05; Suppl. Tables 6 and 7). Single-cell gene expression heatmaps (Figure 6F) showed that only a few of these mRNAs were upregulated in the 3w Aβ hiNS(+) astrocytes. Notably, gene expression overlays showed that some of the most highly upregulated genes following 5w Aβ exposure have previously been associated with the astrocyte response in AD, including *APOE, CLU, SPP1, TLR4, VIM, S100A11* and *LRP1* (Figure 6G). Gene ontology comparing mRNAs upregulated in 5w Aβ hiNS(+) relative to ctrl hiNS(+) astrocytes (Suppl. Table 8) showed that the top-ranked term in the molecular function category was Aβ binding (Figure 6H). By contrast, genes that were significantly increased in ctrl hiNS(+) astrocytes were highly associated with gene ontology terms like neurogenesis and postsynaptic membrane (Suppl. Table 9), suggesting that the presence of Aβ interferes with normal astrocyte biology and cellular interactions.

Next, we compared astrocytes expression profiles, with or without hiMG addition, at 3w as well as 5w, following chronic Aβ exposure, and plotted the results in volcano plots (Figure 6I,J, Suppl Tables 10 and 11). The expression of AD-associated astrocyte genes depends on the presence of hiMG during Aβ exposure, including genes such as *GFAP*, *APOE*, and *VIM.* Interestingly, we found DEGs linked to oxidative stress handling (*OXR1*, *CCL2* and *CHL1*) to be upregulated only in 3w Aβ hiNS(+), the experimental group in which we observed a significant reduction in Aβ induced oxidative stress (Figure 5B). Thus, in hiNS(+) exposed to Aβ for 3 weeks, hiMG increase the expression of subsets of astrocyte marker genes associated with AD as well as the expression of genes handling oxidative stress.

### hiMG drive unique expression profiles in neurons, including APOE

Similar to the subsetted astrocyte snRNA-seq analysis (Figure 6), we subsetted the neuronal populations (both excitatory and inhibitory) for all six experimental groups (Figure S8A). Total neuronal nuclei count reflected the severe cell death caused by 5w Aβ exposure, particularly in the absence of hiMG (Figure S8B). Interestingly, we found that the presence of hiMG during 5w Aβ exposure significantly increased neuronal *APOE, SPP1 and FTL* expression (Figure S8C,D, Suppl. Table 12), similar to the hiMG effect on astrocytes (Figure 6). Since we only observed functional neuroprotection by hiMG during 3w Aβ exposure (Figure 5), we isolated neuronal DEGs in 3w Aβ hiNS(+) (Suppl. Table 13) and 5w Aβ hiNS(+) (Suppl. Table 12) to potentially identify pathways linked to neuroprotection and found 228 and 711 upregulated genes respectively. Gene ontology analysis for 3w Aβ hiNS(+) (Suppl. Table 14) suggests biological processes linked to maintenance of cell-cell and synaptic signaling matching the preserved neuronal activity we observed in the presence of hiMG while 5w Aβ hiNS(+) revealed significant upregulation of processes linked to extracellular exosomes, response to stress, regulation of programmed cell death and amyloid-beta binding (Figure S8E, Suppl. Table 15). Despite the fewer DEGs in 3w Aβ hiNS(+), we performed a number of direct comparisons between treatment groups, similar to our astrocyte gene expression analysis (Figure 6I,J), to identify more subtle genes of interests and graphed them in volcano plots (Figure S8F, Suppl. Tables 16-18). Genes were only plotted if minimum expression percentage exceeded 10% and the upper 10 top genes judging by fold change and/or adjusted pvalue were labeled by name. At 3w Aβ exposure, the timepoint at which the addition of hiMG reverses Aβ-induced oxidative stress and neuronal network dysfunction, is associated with unique gene expression profiles in the combined inhibitory and excitatory neurons. We observed larger differences comparing ctrl hiNS with 3w Aβ hiNS in the absence of hiMG (Figure S8F, left volcano plot) compared to when hiMG were present (Figure S8F, center volcano plot), suggesting that the presence of hiMG preserves the functional gene expression profile of neurons in ctrl hiNS(+). When directly comparing 3w Aβ hiNS(-) with 3w Aβ hiNS(+) (Figure S8F, right volcano plot) we found that a number of AD-linked genes appeared upregulated in 3w Aβ hiNS(+). Several of the highest expressed genes have previously been associated with AD, including *CCL2, VIM* and *STAT3*.

Our data indicate that hiMG drive several genes of interest following different exposure times to Aβ. These genes, including *APOE*, are of high interest for their possible direct role in Aβ-mediated neuronal dysfunction on our hiNS model, particularly as they might relate to differential vulnerability of excitatory and inhibitory neuronal populations in AD.

Considering our astrocytic and neuronal gene expression analysis together, we found larger changes in gene expression in 5w Aβ hiNS(+) astrocytes and neurons while we observed functional neuroprotection only at 3w Aβ hiNS(+) indicating that AD-related reactive astrocyte and neuronal gene expression changes in the surviving population correlate with a more severe pathology at 5w Aβ hiNS(+). This implicates that more subtle gene expression changes at 3w Aβ hiNS(+), both in neurons and astrocytes, should be explored in the future to identify signaling pathways of neuroprotection.

### Heterogenous gene expression profiles in hiMG

After characterizing the shift in astrocyte and neuronal gene expression by hiMG during chronic Aβ exposure, we subsetted the microglia/macrophages (cluster 13, Figure 6A) of the 5w Aβ hiNS(+) experimental group to investigate gene expression profiles in hiMG to better understand their reactive state. We identified 3 distinct clusters within the microglia/macrophage group which we identified further by DEG analysis (Figure 7A, Suppl. Table 19). Cluster 13a showed similarities to a more macrophage-like population characterized by upregulation of *CD36*, *TLR2*, *SYK*, *SLC11A1* and *PTPRC.* Cluster 13c mimics gene expression of homeostatic microglia-like cells including upregulation of *ITGAM*, *CX3CR1* and *P2RY12*. Cluster 13b gene expression was similar to a disease associated microglia (DAM)[45]-like state, sharing a large number of DEGs including *APOE*, *TREM2*, *CD9*, *LPL*, *CD9* and *B2M,* indicating that chronic Aβ exposure induces a similar activation state in a subset of hiMG in 5w Aβ hiNS(+) (Figure 7B). The microglia quantity in the 3w hiNS groups were not sufficient to perform snRNA-seq at this timepoint. We conclude that the microglia/macrophage population in Aβ treated hiNS(+) is heterogeneous and that each subpopulation could play distinct pathophysiological roles. We performed gene ontology analysis on all three clusters highlighting distinct biological processes of each individual group (Figure 7C, Suppl. Table 20). Volcano plots highlighting DEGs genes (Suppl Table 21) of interest for each sub-cluster displayed in Figure 7D (see heatmap with top 10 most significant DEGs in Figure S9).

**Figure 7:**
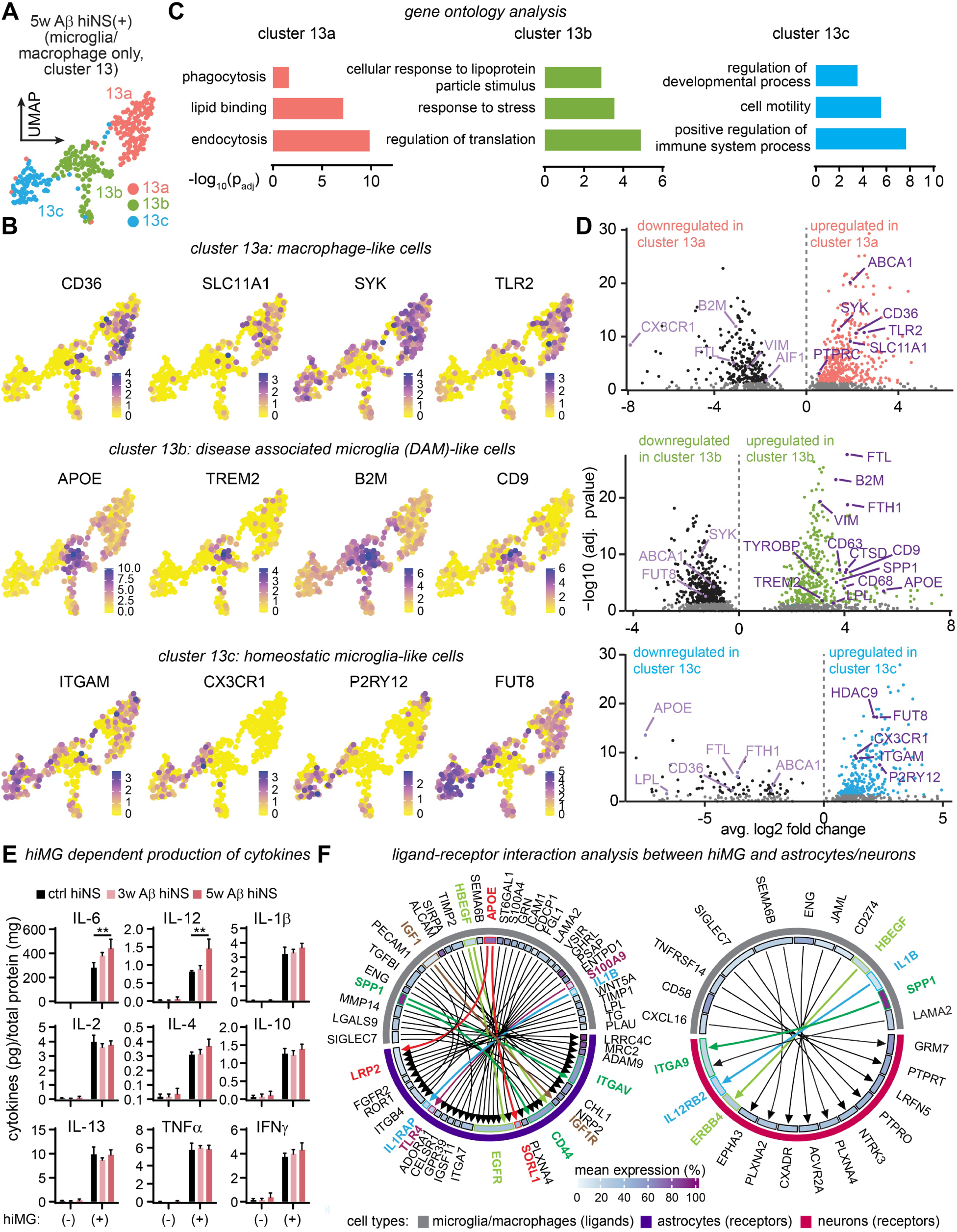
hiMG after chronic Aβ treatment are a heterogeneous cell population of microglia and macrophage-like cells mimicking AD. **A**: The subset of hiMG generated from cell cluster 13 (Figure 5A) from the 5w Aβ hiNS(+) data set reveals three distinct sub-clusters (13a-c). **B**: Overlay examples are shown for each cluster of hiMG. **C**: Gene ontology analysis suggests different biological properties of each cluster. **D**: Volcano plots of DEGs with highlighted genes of interest, including *APOE* in cluster 1 of DAM-like microglia. **E**: Cytokine levels in hiNS concentrated supernatant of all experimental conditions measured via multiplex ELISA (N=3; pooled supernatant from six individual hiNS each). In all hiNS(-) samples, cytokine levels were mostly undetectable or low. Cytokine production increases in hiNS(+) in ctrl and Aβ treated hiNS with IL-6 and IL-12 levels significantly increased from ctrl hiNS(+) to 5w Aβ hiNS(+). Statistical testing performed using two-way ANOVA followed by Holm-Sidak post-hoc tests on logarithmically transformed data. **F**: Ligand-receptor interaction analysis between hiMG ligands paired with receptors on astrocytes and neurons in 5w Aβ hiNS(+). Differentially expressed ligands of microglia/macrophages (cluster 13) and differentially expressed receptors on astrocytes (cluster 0) or neuronal cell populations (all neuronal clusters pooled) were identified and ligand-receptor (mean expression > 10%) pairs were matched and plotted against each other. Genes of interests are highlighted in color.

### hiMG cytokine and ligand expression indicating signaling to neurons and astrocytes

To better understand the phenotypical changes of hiMG during chronic Aβ treatment, we analyzed cell culture supernatant of the above hiNS conditions for proinflammatory cytokine levels using a multiplex ELISA. Supernatant was collected on the last day of the treatment protocol. Overall, cytokine levels were enriched in media of all hiNS(+) samples but not in hiNS(-), indicating that hiMG produce the listed cytokines independent of Aβ exposure. Note that IL-8 data was excluded from the panel due to saturated readings for hiNS (+) groups, preventing accurate quantification. Interestingly, only IL-6 and IL-12 levels increased significantly (** p<0.01) with 5w Aβ treatment, indicating an Aβ induced shift in hiMG cytokine production (Figure 7E).

Our data indicate a significant neuroprotective impact of hiMG in our chronic amyloidosis model by reducing amyloid deposits, oxidative stress and preserving neuronal activity inside hiNS (Figure 4 and 5). hiMG induce significant transcriptomic changes, particularly in astrocytes, mimicking human AD-astrocyte gene expression (Figure 6), as well as neurons (Figure S8). To identify potential pathways by which hiMG exert neuroprotection we performed ligand-receptor interaction analysis based on our 5w Aβ hiNS(+) snRNA-seq data set. DEG lists were generated to identify microglia/macrophage specific ligands (cluster 13), astrocyte specific receptors (cluster 0) and neuronal specific receptors (all neuronal clusters pooled Suppl. Table 22). Ligand-receptor DEG pairs were identified with each having at least 10% mean expression in the respective cell population (Figure 7F, Suppl. Table 23). We identified 44 pairings between hiMG and astrocytes but only 11 between hiMG and neurons, matching our observations that hiMG dramatically change the transcriptional landscape of astrocytes (Figure 6), with fewer changes observed on neuronal populations (Figure S8), with specific pathways of interest highlighted.

### APOE expression in hiNS is hiMG-dependent

*APOE* is well known for being the most important genetic risk factor for sporadic AD and its role in amyloid clearance[46]. The iPSC-line used in this study carries the common homozygous ε allele thought to be ‘neutral’ without increasing or decreasing AD risk. We found astrocytic and neuronal *APOE* expression to be upregulated in 5w Aβ hiNS(+) (Figure 6F,G and Figure S8C,D) as well as in a subset of hiMG (Figure 7B,D) from the same treatment group, suggesting widespread *APOE* expression across multiple cell types in 5w Aβ hiNS(+).

We next immunostained for APOE in fixed slices (80 μm thickness) of hiNS of all six experimental groups and found significant increases in immunoreactivity exclusively in 3w/5w Aβ hiNS(+), indicating that high expression levels of APOE appear only in Aβ-treated hiNS that also contain hiMG (Figure 8A,B). Next, we investigated if APOE is present intracellularly in hiMG and/or astrocytes by co-staining for IBA1 and GFAP. We observed APOE-positive hiMG whereas APOE-positive astrocytes were less frequent, with the majority of APOE apparently extracellular (Figure 8B). We then tested if soluble APOE was detectable by western blot in supernatant from hiNS media of all six groups, and found APOE in all groups containing hiMG, while APOE levels in 5w Aβ hiNS(+) appeared significantly lower compared to ctrl and 3w Aβ hiNS(+). Analysis of soluble Aβ levels using ELISA, showed a reduction in 5w Aβ hiNS(+) compared to 5w Aβ hiNS(-), suggesting potential aggregation of Aβ-APOE complexes in this specific condition resulting in the large quantities of extracellular APOE in 5w Aβ hiNS(+) (Figure 8C), and consequently reduced soluble levels. Additional co-staining for APOE and Aβ (4G8) of 5w Aβ hiNS(±) confirmed that extracellular APOE aggregates co-localized with extracellular Aβ at the outer tissue volumes of hiNS in a hiMG-dependent manner (Figure 8D).

**Figure 8:**
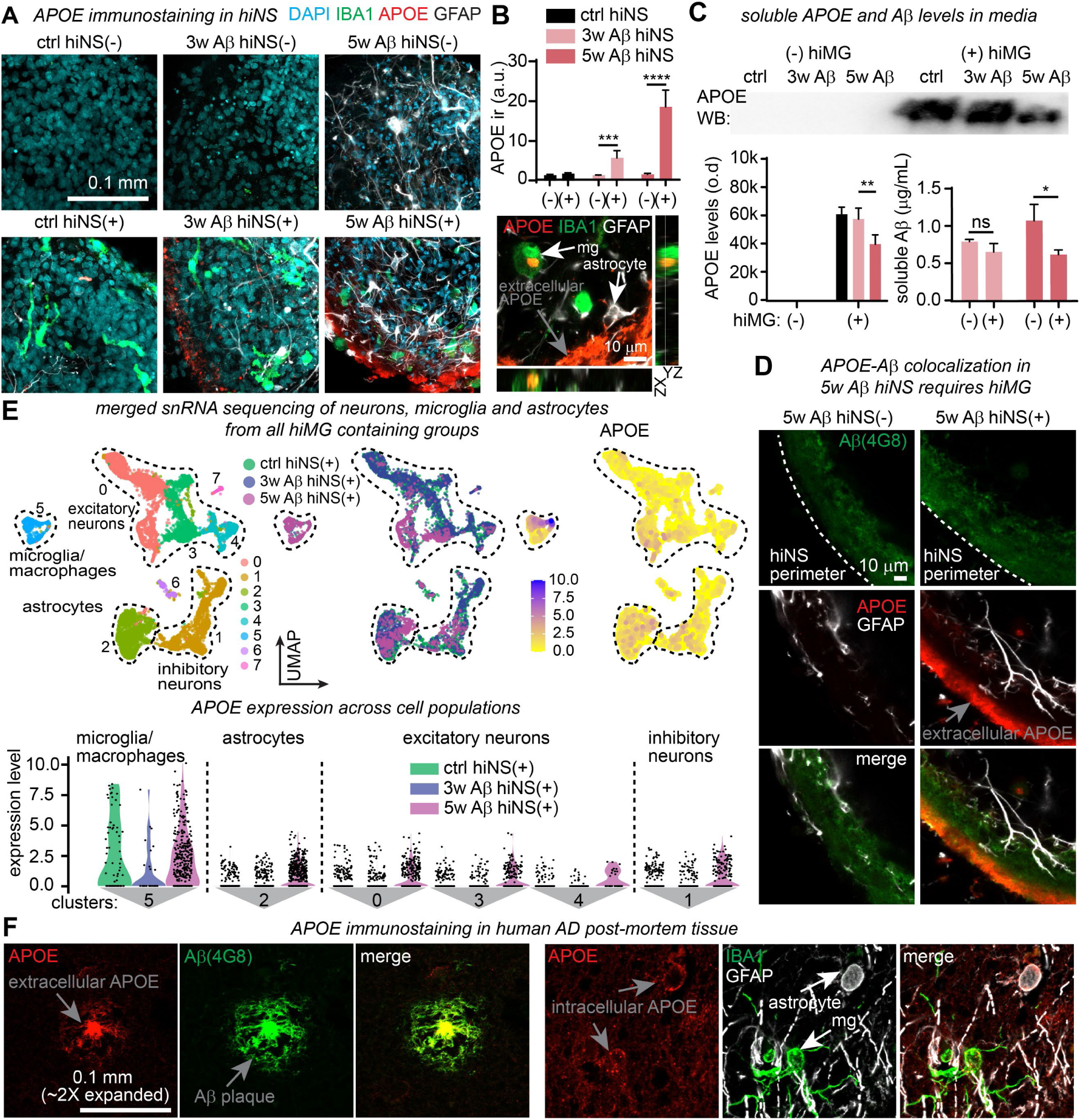
APOE expression in hiNS is driven by hiMG. **A**: APOE immunofluorescence in hiNS slices reveals large quantities of extracellular APOE in 5w Aβ hiNS(+). **B**: (Upper part) Quantification of APOE immunofluorescence (ctrl hiNS(-/+): N=4 each; 3w Aβ hiNS(-/+): N=5 each; 5w Aβ hiNS(-/+): N=5 each). (Lower part) Intracellular APOE identified in hiMG (stained for IBA1, cell labelled ‘mg’) and astrocytes in 5w Aβ hiNS(+). **C**: Soluble APOE and Aβ levels in hiNS supernatant at DIV 60 measured by APOE western blot and Aβ ELISA (4G8) (N=3; pooled supernatant from six individual hiNS each). APOE was only detectable in supernatant of hiNS containing hiMG. At 5w Aβ hiNS(+) soluble Aβ (compared to 5w Aβ hiNS(-)) and APOE levels (compared to 3w Aβ hiNS(+)) were significantly reduced. **D**: Co-staining of APOE with 4G8 indicates that extracellular APOE is in close proximity to Aβ aggregates close to the surface of hiNS (N=3 each). **E**: snRNA-seq clusters of neurons, microglia and astrocytes from ctrl, 3w Aβ and 5w Aβ hiNS(+). hiMG express *APOE* independent of Aβ exposure, while astrocytic and neuronal *APOE* expression depends on the presence of hiMG after 5w Aβ exposure. Note that hiMG expression of *APOE* remains higher than that of neurons/astrocytes. **F**: Immunofluorescence staining in cleared and expanded human AD post-mortem brain sections confirms the close proximity of APOE with Aβ deposits and intracellular APOE in astrocytes and microglia in AD (N=3, 1 sporadic AD, 2 familial AD). All statistical testing performed using two-way ANOVA followed by Holm-Sidak post-hoc tests on logarithmically transformed data

Our snRNA-seq analysis revealed *APOE* expression in microglia, astrocytes/NPCs as well as neurons in 5w Aβ hiNS(+). To compare expression levels between cell types we subsetted all neuronal, astrocytic and microglia/macrophage clusters from all three experimental groups containing hiMG (ctrl, 3w Aβ, 5w Aβ hiNS(+)). Plotting all expression levels side by side confirmed the specific increase in astrocytes and neurons in 5w Aβ hiNS(+) but also indicate that microglia/macrophages express most *APOE*, despite driving *APOE* expression in the other cell types (Figure 8E).

To validate our *APOE* expression data in our AD neurosphere model, we examined APOE immunoreactivity patterns in human post-mortem AD brain sections to confirm known AD APOE expression[47]. To remove background and autofluorescence of paraffin-embedded human cryosections, we cleared and expanded the tissue, based on a recently developed protocol[48] prior to immunofluorescence, and stained sections from two familial AD as well as one sporadic AD patient. In all three patients, we found APOE co-localized with dense-core Aβ plaques, while co-staining for GFAP or IBA1 revealed APOE-positive astrocytes and microglia respectively in AD patients (Figure 8F).

In summary, our 5w Aβ hiNS(+) model mimics *APOE* expression patterns in AD and revealed that microglia, while expressing the highest levels of *APOE*, also drive significant *APOE* expression in astrocytes and neurons. This tool offers unique opportunities to study cell type and APOE isoform specific functions in the future.

## Discussion

### Functional characteristics of hiNS demonstrate healthy brain physiology

Our protocol for the generation of hiNS relies on the differentiation of human iPSC-derived NPCs into neuronal and astrocyte-like cell populations into a spherical 3D neural tissue culture model (Figure 1,2). We extensively characterized the physiological properties of hiNS and showed that neurons in hiNS maintain a healthy reduced redox status with high levels of synchronized wave-like neuronal and calcium activity, matching previous reports of calcium waves in both brain development[49, 50] and other neurosphere cultures[51, 52]. In addition, we found that extracellular glutamate increases at the peak of wave activity and that glutamate transporters EAAT1/2 are required to maintain glutamate balance implicating a direct role of astrocytes. Our pharmacological data suggest that both excitatory and inhibitory neurons contribute to calcium wave formation and dynamics matching other studies[52]. During brain development, not only do neurons and astrocytes play key roles, but infiltrating microglial cells into the brain are vital in shaping the neuronal network[53]. Few other studies[14, 54] have incorporated microglia into neurospheres using different protocols suggesting robust tissue infiltrating properties of iPSC-derived microglia/macrophages. hiMG readily infiltrate hiNS attached to Matrigel-coated surfaces, and we demonstrate that hiMG display microglia-like electrophysiological properties similar to mature *in situ* ramified mouse or human microglia[29, 31, 55]. Moreover, we show that the presence of microglia shapes neuronal calcium wave activity kinetics and also alters the transcriptional landscape of other cell types in hiNS (Figure 1). Our gene expression analysis of astrocytes in hiNS with vs. without hiMG revealed an increase of gene expression linked to mitochondrial function and oxidative phosphorylation, indicating increased metabolic function in the presence of microglia. We believe that our hiNS offer unique opportunities to investigate neuron-glia interaction in human brain development.

### Chronic Aβ exposure models plaque-like aggregation and neurotoxic effects

Instead of utilizing familial AD mutations in *APP* or *PS1* to drive Aβ pathology, we modeled chronic Aβ exposure by supplementing hiNS media with high concentrations of oligomeric Aβ (∼0.24 μg/mL) for 3 to 5 weeks to induce progressive neurotoxicity (Figure 3). We found that oligomeric rather than fibrillar Aβ was more suitable for inducing plaque-like aggregates inside neurosphere tissue within 7 days of media supplementation. Accumulation of Aβ in the brain of AD patients is believed to be one of the earliest pathological events, occurring decades before clinical symptoms[56]. We did not observe any detectable pathological impact within the first 2 weeks of Aβ exposure in hiNS, suggesting that chronic exposure is required to trigger neurotoxicity, mimicking Aβ-driven neurodegeneration in the human brain.

At the end of 3-week long exposure to Aβ we found that calcium wave frequency decreased and that oxidative stress became evident (Figure 3). Oxidative stress is a known early hallmark of AD believed to precede the loss of neurons[57, 58]. It can be observed by immunofluorescence staining of post-mortem AD brain tissue[59]. Here we demonstrate how chronic Aβ exposure over time leads to the build-up of oxidative stress in neurons, which precedes the severe cell death observed after 4-5 weeks of Aβ treatment. Similarly, network dysfunction and synaptic dysregulation[60] are early known features of AD pathogenesis and are believed to precede neurodegeneration and coincide with the formation of Aβ induced oxidative stress in hiNS. Our immunofluorescence staining of hiNS after long Aβ exposure indicates an increase in cleaved Casp-3 immunoreactivity and TUNEL fluorescence, implicating apoptosis as a key cell death pathway. Although caspase-dependent cell death has been proposed as a neuronal death pathway in AD[61, 62], multiple other pathways have more recently been proposed to play a role in later stages of disease progression[63]. More research is needed to decipher the exact cell death pathways in our Aβ treated hiNS, but our model offers a slow-progressing pathology to study key AD disease mechanisms in a human 3D tissue environment.

### Impact of hiMG on Aβ clearance and neuroprotection

While microglia have gained substantial attention in the field over the past 10 years, their precise role in AD pathogenesis remains elusive[64–66]. During the initial stages of AD, microglia are generally believed to release anti-inflammatory cytokines and neurotrophic factors, and are efficient in phagocytosing Aβ; later in AD pathology microglia switch to producing pro-inflammatory cytokines, actively contribute to neurodegeneration and have a reduced capacity to clear Aβ from the brain[67–69]. In our chronic amyloidosis model system, we found that hiMG markedly reduced Aβ levels in the tissue of 3w Aβ hiNS(+) and were found to effectively phagocytose Aβ (Figure 4), further highlighted by positive CD68 immunoreactivity, a marker for phagocytic microglia[40]. Other groups have reported that human iPSC-derived microglia are capable of phagocytosing Aβ[70, 71] matching our findings.

We have demonstrated that hiMG, within a 10-day time window, prevent oxidative stress and sustain neuronal calcium wave activity early during chronic Aβ treatment. However, when hiMG were introduced at a later stage, loss of neuronal activity or severe oxidative stress did not recover, although overall cell death in hiNS was markedly reduced, highlighting that robust neuroprotection could persist (Figure 5B-E). Quantifying levels of a series of pro-inflammatory cytokines from the supernatant of all treatment groups revealed that all measured cytokines were produced in all hiNS(+) groups. However, typical pro-inflammatory cytokines, like IL1β and TNFα, were not elevated in 3w/5w Aβ hiNS(+) when compared to ctrl hiNS(+). Indeed, we saw a robust release of these cytokines (Figure S2) when hiMG were stimulated with LPS, which is known to induce pro-inflammatory cytokine production[72]. We conclude that the production of cytokines in ctrl hiNS(+) marks the baseline levels without indicating a pro-inflammatory phenotype. The only two cytokines significantly increased with Aβ treatment were IL-6 and IL-12 (Figure 7E), which have previously been reported to play either pro- or anti-inflammatory roles[73–77]. Defining the definite roles of these cytokines should be addressed in future studies to shed light on which cytokines have a clear neuroprotective function in our human AD model system.

We conclude that hiMG acquired an overall neuroprotective phenotype in our chronic Aβ model system. In future studies, actively priming hiMG to a pro-inflammatory state, for instance by LPS, prior to adding them to hiNS would be an interesting avenue to pursue.

Recently, Takata et al.[78] have demonstrated in a similar 3D cell culture model that the addition of iPSC-derived primitive macrophages during Aβ treatment resulted in less Aβ aggregation and better survival of neurons, indicating that this activation state of microglia-like cells is consistent between similar model systems. Modelling interaction with additional infiltrating peripheral immune cells is a possible requirement to trigger robust pro-inflammatory phenotypes. Blood-brain-barrier (BBB) breakdown leads to infiltration of soluble factors and cells from the periphery and is a known hallmark of AD and linked to neuroinflammation[79], this process is not modelled in the current study. Most recently, in an amyloid-overproducing 3D tissue culture model comprised of neurons, astrocytes and microglia, the addition of peripheral immune cells, particularly CD8^+^ T cells, greatly facilitated Aβ induced neurodegeneration; although the addition of microglia alone appeared to have neurotoxic effects[14], matching previous observations using the same model system[54]. It is therefore reasonable to suggest that our hiMG could be polarized to a more neurotoxic function by further modelling aspects of BBB breakdown or interaction with peripheral immune cells.

### AD-related astrogliosis and astrocyte activation states depend on hiMG

Cytokines produced by microglia are known to shape astrocyte activation states[80, 81]. We found that the addition of hiMG to hiNS increases the soluble level of a range of different cytokines in the cell culture media (Figure 7E) and we found hiMG to be able to change the astrocytic transcriptomic state under control conditions (Figure 1). We hypothesize that microglial cytokines are the key driver in shaping astrocyte function in our neurosphere model system but further experiments are needed to identify the mechanism by which hiMG alter the transcriptional profile during chronic amyloidosis.

Our immunofluorescence staining revealed a dramatic increase in reactive microglia and GFAP immunoreactivity in 5w Aβ hiNS(+), suggesting gliosis triggered by Aβ exposure (Figure 5A). Reactive astrocytes are known to play both beneficial and detrimental roles in AD pathology[82]. To understand the reactive state of astrocytes in our AD model system we performed snRNA-seq analysis on Aβ treated hiNS with or without hiMG, comparing gene expression levels to control hiNS.

We found that the presence of hiMG shifted the transcriptional profile of the astrocyte cluster in Aβ treated groups. Interestingly, 3w and 5w Aβ hiNS(+) astrocytes appeared to have unique transcriptional profiles (Figure 6C,D). DEG analysis comparing astrocytes of 5w Aβ vs. ctrl hiNS(+), to examine Aβ-induced transcriptional changes exclusively in the presence of hiMG, highlighted significant upregulation of a number of genes associated with reactive astrocytes in human AD: 1) *LRP1* is a receptor for APOE and has been associated with neuroprotection in animal models of AD[83, 84], 2) *CLU* is an extracellular chaperone linked to neuroprotection via Aβ aggregation and clearance by astrocytes[85, 86], and 3) *VIM* was reported to be upregulated in AD although its role in pathology remains elusive[87–89]. Significant gene ontology processes in 5w Aβ hiNS(+) astrocytes included Aβ binding, phagocytosis and endocytosis, indicating that, in the presence of microglia, astrocytes become reactive and could participate in Aβ clearance.

Next, we analyzed astrocytes comparing DEGs of hiNS(-) vs hiNS(+) after 3w/5w Aβ treatment separately to identify more microglia-dependent astrocyte genes. Many DEGs were re-identified to the previous 5w Aβ hiNS(+) vs ctrl hiNS(+) comparison such as *VIM*, *SPP1*, *LRP1* and *S100A11*, highlighting the importance of hiMG in driving astrocyte gene expression (Figure 6I,J). Interestingly, *GFAP* was significantly upregulated in hiNS(+) astrocytes compared to their hiNS(-) counterpart in both 3w and 5w Aβ comparisons. Note that *GFAP* was not differentially expressed when comparing 5w Aβ hiNS(+) to ctrl hiNS(+) astrocytes earlier, although our immunofluorescence analysis strongly suggests an increase in GFAP immunoreactivity indicating astrocyte activation (Figure 5A). Our snRNA-seq quantification of astrocyte numbers (Figure 6E) indicates that 5w Aβ hiNS(+) harbors the largest population of astrocytes compared to all other groups. Therefore, the increased GFAP immunoreactivity could be a result of either or both an increase in astrocyte numbers or increase in *GFAP* expression. This highlights possible limitations of snRNA-seq, as it only captures mRNA still inside the nuclei at the time of tissue collection, not the full transcriptome[90]. Upregulation of *GFAP* in AD-associated astrocytes is a well-known biomarker for AD[88, 91, 92] and is believed to be associated with a neuroprotective reactive phenotype[93, 94]. A recent study described a loss of homeostatic gene expression in AD astrocytes, including downregulation of *NRXN1* and *CADM2*[95]. Both genes were significantly downregulated in astrocytes of 3w/5w Aβ hiNS(+) compared to hiNS(-), indicating hiMG dependence. Another gene of interest that was downregulated is *SLC7A11*. Since its upregulation was recently associated with neurotoxic functions of astrocytes[96], this further supports a neuroprotective astrocyte phenotype in the presence of hiMG.

The presence of hiMG in 3w Aβ hiNS dramatically ameliorated Aβ induced oxidative stress and prevented decline of neural activity (Figure 5B,C). Interestingly, genes linked to oxidative stress response in astrocytes (*OXR1*, *CCL2*, *CHL1)* were significantly upregulated with hiMG, indicating astrocytes could play a role in reducing neuronal oxidative stress independent of Aβ clearance. *OXR1* increases oxidative stress resistance[96]. A recent study found upregulation of *OXR1* following omaveloxolone treatment in a mouse model of AD linked to markedly reduced AD pathology[97]. *OXR1* therefore could be an interesting target for future studies.

### Gene expression changes in surviving neuronal populations after Aβ treatment

Transcriptional changes in the presence of hiMG during Aβ exposure was not limited to astrocytes but extended to neuronal populations (Figure S8). Most strikingly, we found *APOE, SPP1* and *FTL* to be highly upregulated in the surviving 5w Aβ hiNS(+) neuronal populations (Figure S8C,D). *APOE* will be discussed separately below. *SPP1* expression is more commonly observed in immune cells, and elevated FTL levels have been observed in the AD brain[98]. Due to the almost complete death of all excitatory neurons in 5w Aβ hiNS(-) we restrained from attempting any direct 5w Aβ hiNS(+) vs hiNS(-) comparisons in neuronal gene expression patterns. Since we observed functional neuroprotection at 3w Aβ hiNS(+) we investigated DEGs in this experimental group and GO analysis revealed significant hits for terms associated with cell-cell and synaptic signaling, suggesting that the presence of hiMG preserves functional synaptic activity despite the presence of Aβ. When performing GO analysis on the surviving neurons of 5w Aβ hiNS(+) instead, we identified significant terms associated with Aβ binding or the response to stress and programmed cell death (Figure S8E). While we did identify more DEGs in neurons of 5w Aβ hiNS(+) than in 3w Aβ hiNS(+), we performed direct group comparisons of ctrl/3w Aβ hiNS with or without hiMG present to identify potential genes of interests and neuroprotective pathways during amyloidosis. We found stronger differences in gene expression comparing ctrl and 3w Aβ hiNS in the absence of hiMG, while differences with hiMG present yielded much smaller fold changes (Figure S8F). This could indicate, that hiMG are capable of preserving neuronal gene expression profiles of ctrl hiNS(+) maintaining their functional properties. When directly comparing 3w Aβ hiNS(-) with 3w Aβ hiNS(+) we identified genes of interest in AD to be upregulated in the presence of hiMG: *CCL2*, as already mentioned above, is implicated in AD but its precise role in terms of neuroprotection or neurotoxicity remains elusive[99, 100]. *STAT3* is a transcription factor which was shown to be activated by Aβ and is implicated in inflammation driving apoptosis in the AD brain[101, 102]. We speculate that the presence of hiMG, or a subpopulation of them, drives the STAT3 pathway in neurons mimicking aspects of neuronal AD gene expression. *VIM*, as discussed above, is known to be expressed in astrocytes in AD, but has been reported to be expressed in neurons early in the AD brain as part of a damage response mechanism[103].

In conclusion, we believe that further research into astrocytic and neuronal gene expression changes in 3w Aβ hiNS(+) could uncover novel underlying mechanisms of neuroprotection in AD.

### hiMG drive astrocytic and neuronal APOE expression during chronic amyloidosis

One of the most interesting DEGs we identified, and discussed here separately from the others mentioned above, is *APOE*. Our data indicate that microglia could be required to induce astrocyte AD gene expression profiles and speculate that hiMG cytokine production could be the trigger for *APOE* expression in astrocytes since a crosstalk of cytokines and *APOE* expression has been reported in various tissues and model systems[104]

One of the most significantly upregulated genes in 5w Aβ hiNS(+) astrocytes was *APOE*, which is known to be the largest genetic risk factor for sporadic AD and is expressed predominantly by astrocytes, microglia as well as neurons under stress in the human AD brain[46, 105]. Surprisingly, our results show that chronic Aβ (exclusively after 5w Aβ exposure, and not 3w) induced upregulation of *APOE* depends on the presence of hiMG. Much in the same way, hiMG also increased *APOE* expression in neuronal populations, indicating that specifically during long chronic Aβ exposure microglia are key drivers in upregulating *APOE* expression in other cell types in the tissue. *APOE* is closely involved in amyloid clearance, and its alleles ε2 and 4ε are extensively studied for their protective and deleterious roles, respectively[106]. The iPSC-line ε3 alleles not associated with any changes in AD risk. When analyzing the transcriptional profile of hiMG, we found that hiMG in 5w Aβ hiMG(+) sub-cluster into three distinct populations (Figure 7) including one group resembling a unique transcriptional profile of disease associated microglia[45] which includes the expression of *APOE* and *TREM2*, two genes extensively studied in AD research.

Next, we compared *APOE* expression levels between the three cell types in all hiNS(+) groups and found that the population of microglia/macrophage-like cells showed higher *APOE* expression levels compared to astrocytes or neurons, which were similar to each other in expression levels (Figure 8E). Interestingly, we did identify obvious extracellular APOE aggregates in the tissue of 3w Aβ hiNS(+) (Figure 8A) although *APOE* gene expression in this group was not elevated in astrocytes or neurons suggesting that the soluble levels of APOE were produced by hiMG and required the presence of Aβ to manifest as aggregates in the tissue. It is generally believed that astrocytes express most *APOE* in the healthy brain, while gradually microglial *APOE* expression becomes more dominant in AD matching our expression pattern[107].

### hiMG are required for APOE-Aβ co-localization

*APOE* remains the most important risk gene for sporadic AD [46, 108, 109]. Despite this fact, the mechanisms by which each *APOE* expressing cell in the brain impacts AD pathogenesis is poorly understood. APOE is a secreted protein which directly binds Aβ and has been implicated in both its aggregation and clearance, depending on the genetic isoform of *APOE* and stage of AD pathology[110–112]. Besides its role in clearance and aggregation of Aβ, it was established over 20 years ago that amyloid plaques co-localize with extracellular APOE in the human AD brain marking a well-known disease hallmark[47] and was later confirmed in AD mouse models[113] as well. We have confirmed APOE-Aβ co-localization in plaques of AD human postmortem tissue (Figure 8). Interestingly, in our here presented 3D human amyloidosis model we only detected APOE-Aβ interaction if hiMG were present during the Aβ treatment, particularly 5w and to a lesser extend after 3w long exposure, mimicking the colocalization found in the AD brain. Note that soluble APOE and Aβ levels were equal or higher in 3w Aβ hiNS(+) compared to 5w Aβ hiNS(+) (Figure 8C) but the number of hiMG was substantially higher in 5w compared to 3w Aβ hiNS(+) (Figure 5A) further supporting the importance of microglial/macrophage presence for APOE-Aβ to aggregate in the tissue. The functional significance of this co-localization is not well understood.

### Amyloidosis triggered AD-like pathology independent of Tau hyperphosphorylation

During AD pathogenesis, it is generally believed that Aβ accumulation and aggregation is one of the early hallmarks of AD and upstream later hyperphosphorylation and aggregation of Tau[114–116]. We did not find evidence of an Aβ driven increase in Tau hyperphosphorylation, neither with or without the addition of hiMG (Figure S5), by either measuring pTau levels in the tissue or the supernatant. Moreover, we did not find any significant changes in expression of kinases linked to Tau hyperphosphorylation, such as *GSK-3b*, *CDK5*, *PKA* or *MAPK*, in our snRNA-seq data sets, supporting this finding. Our immunofluorescence staining for pTau181 generally indicated bright immunoreactivity in all hiNS, including ctrl hiNS(±). The fact that iPSC-derived neurons are young neurons in early developmental stages and that Tau phosphorylation is a known neurodevelopmental phenotype[117], it is possible that any Aβ induced hyperphosphorylation of Tau is difficult or impossible to detect in our model system. We therefore conclude that our tissue culture model reflects heavy amyloidosis driven features of AD pathology, independent of Tau hyperphosphorylation. Other human 3D tissue culture models *in vitro* have reported a Tau pathology but utilized cells derived from fAD or sAD patients to drive AD pathologies and not a chronic synthetic Aβ stimulation paradigm we use here[118–121]. We conclude that synthetic Aβ supplementation is not sufficient to drive additional Tau pathology in our model.

### Pathway analysis of hiMG ligands matching astrocytic and neuronal receptors

To identify potential pathways of how hiMG affected astrocytic and neuronal function in our 5w Aβ hiNS(+) group, we performed ligand-receptor analysis on our snRNA-seq data (Figure 7F). To highlight the most specific ligands and receptors, we exclusively analysed ligand/receptor DEG pairs with a minimum mean expression of >10%. While determining the exact signaling mechanisms with pharmacological experiments or genetical knockdowns of specific candidates exceed the scope of this manuscript, we here highlight and discuss the most promising pathways in respect to their reported relevance to AD pathogenesis.

We first focused on *APOE* (pathway highlighted in red, Figure 7F). With microglia/macrophages displaying the highest *APOE* expression in 5w Aβ hiNS(+) (Figure 8E), we were not surprised that it was identified as a DEG, despite its expression in astrocytes and neurons. Two receptors, *LRP2* (with a relatively low expression rate) and *SORL1* (with a moderate expression rate) were identified on astrocytes but none on neurons, suggesting that soluble APOE directly acts on astrocytes. *SORL1* is a known AD risk gene and its signalling pathways are more characterized in neurons, rather than astrocytes[122, 123]. In a most recent preprint article, APOE-SORL1 was also shown in LR analysis using spatial transcriptomics on human AD autopsy material[124]. It was further suggested that SORL1-TREM2-APOE signalling is strongly linked to Aβ clearance indicating a functional role of astrocytes in Aβ clearance pathways in our model system[125]. *LRP2* expression in astrocytes has also been linked to Aβ clearance[126] and supports this assumption.

Next, we focused on *SPP1* (pathway highlighted in dark green, Figure 7F), expressed by microglia/macrophages, with the matching receptor genes *CD44* and *ITGAV* on astrocytes and *ITGA9* on neurons. *SPP1* encodes for the soluble protein osteopontin (OPN) which we found to be upregulated in astrocytes in the presence of hiMG, presumably independent of Aβ in ctrl (Figure 1), 3w and 5w Aβ hiNS(+) (Figure 6). Note that *SPP1* was also differentially expressed in the DAM-like microglia/macrophage population (Figure 7D, cluster 13b volcano plot), the same cluster expressing *APOE* and *TREM2*. OPN has been has been recently suggested as an early AD biomarker[127] and its deletion or reduction through anti-OPN antibodies was reported to reduce plaque burden in AD mice suggesting an overall neurotoxic function during AD pathogenesis[128]. One of its well described receptors is CD44 and highly expressed in astrocytes of AD patients[129], which we found to be upregulated by the presence of hiMG with and without Aβ exposure. A second OPN receptor we found to be expressed on astrocytes is Integrin-α5 (encoded by *ITGAV*), which was shown to exert neuroprotective functions in a mouse model of AD[130]. Another OPN receptor Integrin-α5 (encoded by *ITGA9*), part of the integrin family, was found to be expressed on neurons suggesting that OPN signaling is not limited to microglia/macrophage to astrocyte crosstalk and involves multiple members of the integrin receptor family.

Another microglia/macrophage ligand we identified, with different receptor genes on astrocytes (*EGFR*) and neurons (*ERBB4*), is *HBEGF* (pathways highlighted in light green, Figure 7F). *HBEGF* encodes for heparin-binding epidermal growth factor (HB-EGF) and was suggested to be an endogenous neuroprotective molecule in the brain[131] and inhibition of HB-EGF was reported to facilitate Aβ expression in rats[132]. The role of its receptors we identified in our data set is more documented to be involved in AD. Inhibiting EGFR was shown to exert neuroprotection in AD[133] highlighting it as a novel potential target for AD[134] overall suggesting that EGFR plays a neurotoxic role in AD pathogenesis[135]. On the neuronal side, we identified *ERBB4* (encoding for ErbB) which was shown to be expressed in neurons of AD mice[136] and was also associated with neurotoxicity[137]. The conflicting literature on HB-EGF, as a neuroprotective factor, and the neurotoxicity associated with the receptors EGFR/ErbB further stresses the need of novel model systems to investigate these pathways in more depths and their therapeutic potential for AD treatment.

We further identified *IL1B* (encoding for IL1β, (pathway highlighted in blue, Figure 7F)) as a differentially expressed microglia/macrophage ligand matching the genes *IL1RAP* (on astrocytes) and *IL12RB2* (on neurons). The role of IL1β signaling in AD is well documented as a pathological factor [138] and reported to be upregulated in AD, driving astrocyte activation and neuroinflammation overall suggesting a neurotoxic contribution in AD[139, 140], potentially acting through astrocytes and neurons in our experimental model.

The role of Insulin-Like Growth Factor -1 (IGF-1) (encoded by *IGF1,* identified ligand expressed in our microglia/macrophage population in low levels, (pathway highlighted in brown, Figure 6F)) signaling in neuroinflammation and AD is controversial and context dependent. While an anti-inflammatory role promoting neuronal survival through its receptor, IGF-1R (encoded by *IGF1R*, identified ligand expressed in our astrocyte population in moderate levels) on neurons and astrocytes is well supported [141] another study reports that blocking IGF-1R results in lowered Aβ levels and reduced microglia activation in a mouse model of AD[142].

Another potential crosstalk implicated in AD we identified is the ligand gene *S100A9* and *TLR4* (pathway highlighted in purple, Figure 7F) expressed on astrocytes. While *TLR4* expression is well documented for microglia, its role in astrocytes, particularly in AD has been suggested[143]. The agonist S100A9 was reported to be expressed by activated microglia surrounding Aβ plaques implicated in phagocytosis, Aβ aggregation and neurotoxicity[143, 144]. To what extend and where S100A9 and TLR4 are expressed in our 5w Aβ hiNS(+) model system would be worth pursuing in the future.

In summary, we found more ligand-receptor matches between microglia/macrophages with receptors on astrocytes rather than neurons, suggesting a higher level of glia-glia rather than microglia/macrophage-neuron interaction. We identified several pathways of interests, both linked to neuroprotection or neurotoxicity in AD pathology indicating that our model partially mimics both aspects of AD allowing the investigations of these pathways and their relevance to Aβ pathology in future studies.

## Conclusions

This study introduces a human iPSC-derived neurosphere model, including neural progenitor cells, neurons, astrocytes and microglia residing in a healthy brain-like 3D environment. We have performed extensive functional characterizations of our neurospheres demonstrating synchronized neural activity, the absence of necrotic tissue in the core, and characterized functional properties of microglia inside the tissue, demonstrating surveillance behaviour and damage response properties.

Using this tool, we developed a novel chronic amyloidosis model by supplementing media with Aβ to gradually induce an AD-like pathology that effectively recapitulates key hallmarks of AD, providing a robust platform for studying the interplay of neurons, astrocytes, and microglia in chronic amyloidosis. Through this system, we demonstrated that microglia exhibit critical neuroprotective properties by mitigating Aβ induced oxidative stress, preserving neuronal activity, and reducing apoptosis. The timing of microglial infiltration during chronic amyloidosis determined the extend of their neuroprotective properties. The flexibility of this model to introduce microglia at a defined time during chronic amyloidosis driven neurodegeneration offers a unique platform to study their impact on AD pathology.

Importantly, we identified microglia as pivotal drivers of unique expression profiles in neurons and astrocytes, with several transcriptional changes resembling that found in human AD. This microglial-dependent modulation of astrocytic and neuronal responses underscores the intricate cell-cell signaling that governs disease progression. Although our model successfully mimics Aβ-driven neurotoxicity and transcriptional shifts, it does not recapitulate Tau hyphosphorylation, highlighting its specificity for investigating amyloid centered mechanisms.

By bridging cellular, molecular, and transcriptional dynamics in a human context, this system provides a scalable and translationally relevant platform for preclinical research. Future studies leveraging this model could elucidate the pathogenic effects of identified gene expression profiles, including APOE isoforms, to explore the earliest, cell-specific pathophysiologic changes in the human brain in response to Aβ

## Acknowledgments

For identifying the *APOE* genotype of the used iPSC line we thank Dr. Mari DeMarco for assistance.

## Declarations

### Data availability

Our snRNA-seq has been uploaded to Gene Expression Omnibus (GEO) and can be found under the accession number GSE272186. Other data are available from the corresponding authors upon reasonable request.

### Ethics approval

Our work on hiPSCs was approved by the University of British Columbia Clinical Research Ethics Board (H21-02261).

### Author contributions

Conceptualization, H.B.N., S.W.; Methodology, S.W., S.N.E., F.M., K.K., J.F.; Experimental and Analysis, S.W., A.J.L., S.N.E., D.J.B., W.C., Y.B., D.Y.K., H.B.C., M.D.; S.S., C.J.G., C.L.; Resources, I.R.M.; Writing, S.W., H.B.N., A.J.L., B.A.M., F.M.; Visualization, S.W., A.J.L, K.K.; Supervision, H.B.N., B.A.M., F.M., D.R.K.; Funding Acquisition, H.B.N., B.A.M., F.M., S.W.

## Methods

### Generation of human iPSCs and induction of NPCs

The presented work on hiPSCs was approved by the University of British Columbia’s Clinical Research Ethics board. A single cell line was used, obtained from a healthy female individual carrying the *APOE* alleles ε3/ε3. Peripheral blood was drawn and hiPSCs were generated from erythroid progenitor cells. The cell line has been used and characterized by our laboratory in the past[145, 146].

hiPSCs were grown on Matrigel (BD Biosciences, Franklin Lakes, USA) coated 6-well plates in mTeSR^TM^ Plus (STEMCELL Technologies, Vancouver, Canada). Media was supplemented with 10 μM Y-27632 (EMD Millipore, Burlington, USA) for 24h after plating. Cells were fed every 2-3 days and used for both the induction of neural progenitor cells (NPCs) and microglial differentiation. For storage, cells were frozen in media, supplemented with Y-27632 and 10% DMSO (EMD Millipore) in liquid nitrogen (LN).

hiPSCs to NPC differentiation was achieved by a dual SMAD inhibition protocol as described before[147]. In short, hiPSCs were plated in 6-well plates in Basal Neural Maintenance media (BNMM) (Suppl. Table 1) supplemented with 10 μM SB-431542 (Tocris, Bristol, United Kingdom) and 0.5 μM LDN-193189 (Tocris) for 7 days, followed by a 3-week differentiation in Neural Stem Cell Media (see Suppl. Table 1) with media changes every 2-3 days. NPCs were then expanded and passaged twice before freezing them down in LN (BNMM supplemented with Y-27632 and 10% DMSO) at P2.

### Neurosphere generation

To generate human iPSC-derived neurospheres (hiNS), P2 NPCs were thawed and plated in Matrigel-coated 6-well plates using Neurosphere Expansion Media (NEX) (Suppl. Table 1) supplemented with 10 μM Y-27632 for 24h. Cells were fed every 2-3 days and allowed to recover for at least 5 days until the cell density was sufficient to start hiNS generation. To start hiNS formation, NPCs were lifted using Accutase (STEMCELL Technologies) and plated in single wells of AggreWell^TM^800 plates at a density of 1.2 * 10^6^ NPCs per well in NEX supplemented with 10 μM Y-27632 for 24h and centrifuged for 2 min at 100g. hiNS were fed daily by careful partial media changes until day 7 (days in vitro, DIV 7). At DIV7 hiNS were dislodged using wide bore tips and 37 μm reversible strainers (STEMCELL Technologies) and transferred to standard 9 cm cell culture petri dishes (VWR International, #10861-592, Edmonton, Canada) in Neurosphere Differentiation Media (NDM) (Suppl. Table 1) and immediately placed on an orbital shaker (Celltron, Infors HT, Bottmingen, Switzerland) in the cell culture incubator at 65 rpm. hiNS were fed every 2-3 days by full media change and switched to Neurosphere Maintenance Media (NMM) (Suppl. Table 1) on DIV 28. hiNS were kept in the petri dishes until DIV 39-41 when they were transferred to Matrigel coated (1% overnight) single wells of 48-well plates for future experiments either on regular cell culture plates, 8 mm coverslips or glass bottom live imaging plates (MatTek, Ashland, USA). For precise positioning on glass surfaces a tissue dissection microscope (Zeiss Stereo Discovery V8) was used inside a biosafety cabinet. Plated hiNS readily attach to surfaces allowing for easy confocal live imaging. To induce the expression of genetically encoded fluorescent proteins, hiNS were transduced with 0.5-0.75 μL of custom AAVs (AAV2/PHP.eB) on the day of plating. Promoters used were *gfaABC1D* (for EGFP) and *hSyn* (for mCherry, roGFP1, GCaMP6f and iGluSnFr) and all AAVs titers ranged from 1-2 * 10^13^ gene copies per mL. After plating and transduction, hiNS media was partially changed every 2-3 days.

### Differentiation of human iPSC-derived microglia

Human iPSC-derived microglia (hiMG) were differentiated based on a protocol by the Cowley lab with minor adjustments[145] as follows. hiPSCs were collected from 6-well plates using Accutase and 1.5 * 10^6^ cells were added to a single well of an AggreWell^TM^800 plate in Embryoid Body Media (Suppl. Table 1) and centrifuged at 100g for 2 min. Media was changed daily for 4 days. On day 5, EBs were collected using a wide bore tip and a 37 μm reversible strainer and resuspended in 10 mL Hematopoietic Differentiation Media (HDM) (Suppl. Table 1) in a standard T75 Cell Culture flask. The flask was kept in the incubator for 7 days without feeding or moving for EBs to attach to the surface. Afterwards, EBs were fed with 5 mL of HDM twice a week. As early as day 35, accumulating macrophage precursors were harvested for subsequent microglia differentiation. Collected HDM enriched with macrophage precursors was centrifuged at 400g for 5 min and cells were resuspended in Microglial Differentiation Media (MDM) (Suppl. Table 1) and plated in 6-well glass bottom plates at densities of 0.5-1 * 10^6^ cells/well. Partial media changes were conducted every 2-3 days with MDM and hiMG were allowed to differentiate for 10-14 days.

### hiMG transfer into neurospheres

Mature hiMG after 10-14 days of differentiation were collected using TrypLE Express (Thermo Scientific, Waltham, USA) and resuspended in NMM supplemented with 100 ng/mL IL-34 (BioLegend, San Diego, USA)) and 25 ng/mL TGFβ1(PeproTech, Cranbury, USA) and added to well-attached hiNS in 48-well plates at a density of 35,000 cells/well between DIV 49-60. hiNS(+) were fed with partial media changes every 2-3 days with NMM supplemented with IL-34/TGFβ1. Note that IL-34/ TGFβ1 supplementation was performed for all treatment groups, including hiNS(-) throughout the experiment.

### ELISA

For multiplex ELISA assays of hiNS media, the supernatant of 6 individual hiNS (across all treatment groups) was pooled together and frozen at -80°C. Before analysis, samples were thawed on ice and concentrated using Amicon Ultra 3kDa centrifugal filters (EMD Millipore) by centrifuging 500 μL at 14,000g for 15 min. This step was repeated twice until all media was concentrated. Concentrated media was then collected after flipping the filter and centrifuging again at 1000 g for 2 minutes. The resulting total volume for each pooled sample of concentrated media was ∼80 μL. For soluble Aβ and pTau181 quantification the V-PLEX Aβ Peptide Panel 1 (4G8) and S-PLEX Human Tau (pT181) kit (Meso Scale Diagnostics, Rockville, USA) was used respectively for electro chemiluminescent detection of Aβ38, Aβ40, Aβ42 and pTau181 in concentrated supernatant. Samples were diluted 1/5,000 in Diluent 35 (provided in the kit). All samples and calibrators were run in duplicate following manufacturers’ instructions. To quantify cytokine expression in hiNS media, the same samples were used for the V-PLEX Cytokine Panel 1 Human Kit (Meso Scale Diagnostics) for the quantification of ten cytokines: GM-CSF, IL-1α IL-5, IL-7, IL-12/IL-23p40, IL-15, IL-16, IL-17A, TNF-β, and VEGF-A. All concentrated samples were diluted 15x and all samples/calibrators were run in duplicate following the manufacturer’s instructions. MESO QuickPlex SQ 120MM was used for plate reading and protein concentrations were determined using the Discovery WorkBench Software provided by Meso Scale. For cytokine quantification, concentrated supernatant protein concentration was detected via BCA for normalization of cytokines between samples.

### Oligomeric Aβ preparation

Oligomeric Aβ preparation was performed based following well-established and characterized protocol[32–36]: Synthetic lyophilised Aβ1-42 (ERI Amyloid Laboratory, New Haven, USA) was mixed thoroughly and dissolved in 40 μL of DMSO for 20 minutes at room temperature (RT) at a concentration of ∼2 mM. The mixture was then separated into two 20 μL aliquots and 1 mL of F12 (Thermo Scientific) media was added to each aliquot. The dissolved Aβ was incubated at RT for 24 hours to induce oligomerization. Next, 500 μL of the oligomeric Aβ1-42 (Aβ) was centrifuged at 14,000 g for 15 minutes using Amicon Ultra 3kDa centrifugal filters. This was repeated 3 times to concentrate the A. The A was then washed twice with 500 μL of PBS and centrifuged at 14,000 g for 15 minutes. The filters were then flipped onto collection tubes and centrifuged at 1000 g for 2 minutes to collect the Aβ. The final Aβ was stored at –80°C. To determine the final Aβ concentration we used a multiplex Aβ (6E10) ELISA panel (see above) on four different batches of Aβ and averaged the results. The final stock concentration of Aβ was 0.24 ± 3.4 mg/mL and was used throughout the experiment in a 1:1000 dilution, unless stated otherwise.

To conjugate Aβ with pHrodo red/green (Thermo Scientific), 2 μL were added into each 1 mL of the F12/Aβ mix prior to centrifugation and incubated at 4°C in the dark for 3 hours. Centrifugation steps were continued afterwards as described above with one additional PBS washing step and the conjugated pHrodo-Aβ was stored in the dark at -80°C.

### Fibrillar Aβ preparation

For fibrillar Aβ-1-42 (fAβ) generation(adapted from[148]) (see Figure S3), the same lyophilised Aβ1-42 powder was used as described above. After dissolving the powder in 40 μL of DMSO for 20 minutes at room temperature (RT), and splitting the solution in two, 980 uL of 10 mM HCl was added each. Tubes were then stored at 37°C overnight. Afterwards, the same filtration steps were taken to concentrate the fAβ.

### Chronic Aβ treatment paradigm

To model chronic amyloidosis in our 3D neurosphere cultures, we supplemented NMD/NMM with Aβ in a 1:1000 dilution resulting in a final Aβ concentration in the media to be 0.24 μg/mL. Aβ was added during partial media changes (150 μL from a total of 400 μL per well) calculated for the total volume of the wells resulting in a gradual increase of aggregated Aβ over time. The duration of Aβ treatment was either 3 or 5 weeks. For the control groups (ctrl hiNS(-/+)), media was supplemented with 3.6 μg/mL scrambled Aβ 42 for a total of 3 weeks.

### Confocal microscopy of neurospheres

For live neurosphere imaging, hiNS were plated and transduced with AAVs in 48-well glass bottom plates as described above. hiNS were imaged 7-10 days post transfection using a confocal microscope equipped with a live imaging incubator system (LSM 880, Zeiss, Oberkochen, Germany). For *hSyn*-GCaMP6f, *hSyn*-iGluSnFr and *gfaABC1D*-GCaMP8s 3-5 min time series were recorded of green emission excited using a 488 nm argon laser. hiNS display synchronized wave-like neuronal and calcium activity leading to rises in intracellular calcium levels (quantifying GCaMP6f or GCaMP8s fluorescence) or a rise in extracellular glutamate concentrations (quantifying iGluSnFr) (Figure 2). To simultaneously image *hSyn*-iGluSnFr and *gfaABC1D*-GCaMP8s with neuronal calcium levels, we co-transduced to express *hSyn*-jRGECO1, a red fluorescent calcium indicator[149]. AQuA[150] was used for calcium wave quantification as follows. For each hiNS, calcium activity was recorded at one z-plane for 300 frames at a frame rate of 0.931 seconds per frame, and the resolution was 0.692 μm per pixel. Prior to analysis, the resolution was downsized to 1.384 μm/pixel in FIJI to allow for smoother processing. In AQuA, an ROI was drawn around each neurosphere. For event detection, AQuA uses the parameters intensity threshold, smoothing, and minimum pixel size, which were adjusted so that signals that spanned the entire neurosphere were counted as a single event, and other background single cell activity was classified as noise and ignored.

To quantify neuronal redox states in hiNS we employed roGFP1[18], a genetically encoded redox indicator we have used in the past[19]. roGFP is a ratiometric indicator in which upon oxidation, the excitation spectrum of roGFP1 shifts to lower wavelengths (∼400 nm) while fully reduced roGFP1 has a maximum excitation peak at ∼490 nm. The green fluorescence of roGFP1 on the other hand is not affected by its redox states. Therefore, roGFP1 was imaged on our live imaging confocal microscope alternately with either a 405 nm laser (maximum excitation for oxidized roGFP1) and a 488 nm laser (maximum excitation for reduced roGFP1) to quantify neuronal redox states across experimental conditions. Laser settings were kept constant and green emission for both excitation wavelengths were captured and averaged across a 30 μm z-stack. The ‘*roGFP1 ratio’* (Figure 1) was defined as green emission acquired exciting at 405 nm and divided by green emission acquired exciting at 488 nm. For one data set (Figure 3D) hiNS expressing roGFP1 were fixed in 4% PFA substituted with 20 mM N-Ethylmaleimide to preserve the redox state of roGFP1 allowing post-fixation redox imaging[19].

To quantify hiMG infiltration into hiNS and simultaneously image neurons and astrocytes, we pre-stained differentiating hiMG in their respective 6-well glass bottom plate for 30 min with tomato-lectin-649 (Thermo Scientific) in a 1:1000 dilution and subsequently added to hiNS co-transduced with AAV.php.eb.-*hSyn*-mCherry and AAV.php.eb.-*hSyn*-EGFP. hiNS and hiMG were then live imaged for 24h (see Figure 2F).

### Human formalin-fixed paraffin-embedded (FFPE) Magnified Analysis of Proteome (MAP) tissue clearing and expansion

The FFPE MAP procedure was adapted from the original MAP procedure[151] with slight modifications[48]. Three FFPE brain sections with 4 μm thickness were used, two familial AD patients and one sporadic AD patient.

#### Deparaffinization and rehydration

High precision #1.5 coverslips were cut and glued directly on the original slides to which the sections were attached on each side of the sections. These served as spacers for the gel embedding (see below). Then sections were deparaffinized and rehydrated by washing them with xylene three times for 5 min each and then two times for 5 min each in the following solutions: 100% ethanol 95% ethanol, 70% ethanol, and 50% ethanol. Finally, the samples were rinsed two times with deionized water for 5 min each. From this step onward, samples were protected from light.

#### Post-fixation

Sections were kept in fixative solution (4% PFA in PBS) for 2 h at 37°C then washed three times in washing solution (PBS with 0.02% of NaN3) from 30 min each at 37°C with gentle shaking.

#### AA-integration

Sections were incubated in low-AA solution (4% acrylamide (Sigma Aldrich, A3553), 4% PFA in PBS) at 4°C with agitation overnight, followed by a 2 h incubation at 37°C with gentle shaking.

#### Inactivation

Brain sections were washed with washing solution three times at 37°C for 30 min each. Sections were then incubated in Inactivation solution (1% acetamide (Sigma Aldrich, A0500), 1% glycine (Sigma Aldrich, G7126), 0.02% NaN3 (Sigma Aldrich, S2002) with the pH titrated to

9.0 with NaOH) at 37°C for 3 h with gentle shaking.

#### Monomer incubation

Brain sections were washed with washing solution three times at 37°C for 30 min each. MAP solution was made by freshly adding V-50 initiator (Sigma Aldrich; 440914) to a solution of 30% acrylamide, 0.1% bisacrylamide (Sigma Aldrich, M7279), and 10% sodium acrylate (Sigma Aldrich, 408220) in PBS; the brain sections were incubated in MAP solution overnight at 4°C with gentle shaking.

#### Mounting and gel embedding

The gelation was done directly on the original slides to which the sections were attached. 100 μL of MAP solution with freshly added V-50 initiator was placed directly on each section and topped with a coverslip to form gelation chambers. Gelation chambers were placed in a humid gelation box which was sealed, and the air purged with nitrogen gas. The gelation box was put in the oven at 45°C for 2 h 30 min, following which the hybrid gel–tissue was removed and excess gel around the tissue cut away.

#### Denaturation, clearing, and expansion

The tissue–gel hybrid sections still attached to the slides were immersed in denaturation solution (in mM, 200 SDS, 200 NaCl, 50 Tris, pH 9) and incubated at 65°C for 3 days with gentle shaking using an EasyClear device (LifeCanvas Technologies). The FFPE-MAP tissue–gel hybrids generally floated free of the slides after denaturation. Sections were washed 4 times for 1 h each with 0.001× PBS and gentle shaking and then incubated in 0.001× PBS at RT overnight with gentle shaking. To prepare the sections for immunostaining, the 0.001× PBS was replaced with PBST (0.1% Triton X-100 in PBS) for 1 h before staining. The side that had been facing the original slide, thus which had the tissue at the surface, was mounted next to the coverslip for imaging.

### Immunofluorescence

For immunofluorescence staining, hiNS were plated on 8 mm glass coverslips and fixed in 4% paraformaldehyde for 1h. Fixed hiNS were washed and kept in PBS at 4°C before staining. Immunofluorescence stainings were conducted as described before[19]. hiNS were either stained in full or sliced with a vibratome into 80 μm thin sections (VT1200, Leica, Nussloch, Germany) before immunostainings. In brief, hiNS (full or slices) were blocked with 10% normal donkey serum (NDS) (Jackson ImmunoResearch, West Grove, USA) in 0.01M PBS containing 2% Triton X-100 and 20% DMSO for 1h. Afterwards, samples were washed once with PBS before incubating in primary antibody solution containing 0.1M PBS, 2.5% NDS, 2%Triton X-100 and 20% DMSO. Primary antibodies used were Aβ (6E10), 1:500 (BioLegend, San Diego, USA); Aβ (4G8), 1:500 (BioLegend); Aβ (mOC87), 1:500 (abcam, Cambridge, United Kingdom); APOE (D7I9N), 1:1000 (New England Biolabs, Ipswich, USA); GFAP, 1:1500 (Thermo Scientific); MAP2, 1:1500 (abcam); IBA1, 1:500 (FUJIFILM Wako Chemicals, Richmond, USA); Cleaved Caspase-3, 1:500 (Cell Signaling Technologies, Danvers, USA); CD68, 1:200 (Novus Biologicals, Toronto, Canada); Tau (HT7), 1:500 (Thermo Scientific); pTau181 (D9F4G), 1:500 (Cell Signaling Technologies). Afterwards, samples were thoroughly washed with PBS 3-4 times before incubating in secondary antibody solution containing 0.1M PBS, 2.5% NDS, 2%Triton X-100 and 20% DMSO and secondary antibodies targeted against the host species of primary antibodies conjugated with either Alexa Fluor 405, 488, 568 or 647 (Thermo Scientific) and incubated for 1h. In some stainings, DAPI (Thermo Scientific) was added in a 1:5000 dilution for 15 min, after the secondary antibody staining. For quantification of apoptosis, TUNEL (Click-iT^TM^ Plus, Invitrogen) was used following the manufacturer guidelines after immunostainings were performed. After 3-4 washing steps in PBS, stained hiNS were imaged on the LSM 980 confocal microscope (Zeiss). Quantification of fluorescence was performed using FIJI (ImageJ 1.53t). The immunostaining protocol for FFPE MAP human sections was performed the same way except for using a 0.1% Triton-X-100 PBS solution for blocking and staining buffers instead of the 2% Triton-X-100/20% DMSO mix.

### Western Blot

Western blot analysis was performed on differentiating hiMG harvested at different time points. Cell lysates were prepared using RIPA lysis buffer supplemented with phosphatase and protease inhibitor cocktail (Roche, Basel, Switzerland), followed by centrifugation at 13,000g for 10 minutes at 4°C. The resulting supernatant was diluted with 4x Laemmli sample buffer and boiled for 5 minutes before loading onto Mini-Protean TGX gels (Bio-Rad, Hercules, USA). Subsequently, SDS-PAGE was conducted, and the separated proteins were transferred onto PVDF membranes using a Trans-Blot Turbo semi-dry transfer system (Bio-Rad). Membranes were blocked using 3% BSA in TBS with 0.1% Tween20 (TBST), followed by overnight incubation at 4°C in TBST with primary antibodies against IBA1 (GTX635400, 1:500, GeneTex, Irvine, USA,), APOE (D7I9N), 1:1000 (New England Biolabs) or GAPDH (PA1988, Thermo Scientific, 1:1000). After washing, membranes were probed with HRP-conjugated secondary antibodies (Jackson ImmunoResearch) for 2 hours at room temperature. Protein bands were visualized using Bio-Rad Clarity Western ECL substrate and imaged with a ChemiDoc MP imaging system. Protein band size and intensity were quantified using ImageJ software.

### Electrophysiology and 2-photon imaging

Patch-clamp recordings on microglial cells were performed on hiNS(+) either ctrl or Aβ (3w/5w) treated after plating them on coverslips, which allowed moving them to a 2-photon microscope for patch-clamp and image recordings (Zeiss 510Meta microscope with a Ti:Sapphire Chameleon laser). hiMG were stained with 2 μL tomato-lectin-488 (for patch-clamp) or 594 (for process motility quantification) (Thermo Scientific) per well for 30 min to allow for the identification of microglia in neurospheres using the 2-photon microscope. hiNS were supplied with a continuous perfusion of carbogen (95% O_2_/ 5% CO_2_) infused BrainPhys Imaging Optimized media (STEMCELL Technologies) and the following intracellular solution was used for microglial patch-clamp recordings, as described before (in mM)[29, 30]: KCl,130; MgCl_2_,2;CaCl_2_, 0.5; Na-ATP, 2; EGTA, 5; HEPES, 10 and sulforhodamine 101, 0.01 (Sigma Aldrich) at an osmolarity of ∼370 mOsm and a pH of 7.3. Borosilicate patch pipettes were pulled with a final pipette resistance of 3-5 MΩ. hiMG were targeted using the lectin fluorescence and clamped at -40 mV after reaching whole-cell configuration. A series of de- and hyperpolarizing voltage steps were applied ranging from -150 to + 60 mV with 10 mV increments and patched hiMG with abnormally small capacitances (<10 pF) or a series resistance of >65 MΩ were discarded. To test if hiMG perform surveillance functions and respond to tissue damage, ctrl hiNS(+) (N=3) were stained with lectin-594 and z-stack (27 μm thick) time series were acquired using our 2-photon microscope at 850 nm. After recording a baseline to monitor process movement, a laser lesion was burned into the centre of the neurosphere by tuning the laser to 800 nm at 100% power for 15 seconds. hiMG process movement towards the lesion was then imaged again using 850 nm excitation wavelengths in a z-stack time series recording.

### Single nuclei RNA sequencing

Single nucleus capture was performed at the Princess Margaret Genomics Centre (Toronto, ON) using the relevant Genomics Chromium system protocol. Libraries were sequenced on the Illumina NovaseqX Plus instrument. FASTQ reads were aligned to the wild-type human genome (hg38) using the relevant 10x Genomics Cell Ranger software. Output data was analyzed using a previously described computational pipeline[20–24]. In brief, cells with low UMI counts, red blood cells, cells with high mitochondrial DNA content, and putative doublets were removed. Genes detected in fewer than three cells were removed and cell transcriptomes were normalized using the scran R package to account for read depth and library size[21]. The Seurat package (v4.2.0)[20] was then used to process the normalized expression matrices. Following this, principal component analysis (PCA) was performed using the top 2000 highly variable genes. Next, the top principal components were used to project the dataset into two-dimensional space as Uniform Manifold Approximation and Projection (UMAP) embeddings using the RunUMAP function in Seurat. Afterwards, the top principal components were also used to iteratively carry out SNN-Cliq-inspired clustering using the FindClusters function in Seurat at resolutions ranging from 0.2 to 2.4. Each dataset was analyzed by choosing the most conservative resolution; resolution was only increased to interrogate cell heterogeneity or if two cell populations of known identity, based on the expression of canonical markers, were not separating into distinct clusters as expected. To merge or subset datasets, barcodes corresponding to the cells or cell types of interest were used to extract relevant gene expression information from filtered gene expression matrices. For the merging of datasets, the gene expression information from each respective dataset was then combined. The cells were then once again run through the pipeline as described above.

For the hiNS(-/+) merged dataset (as in Figure 1C), clusters were assigned at a resolution of 0.4 (16 clusters identified with at least 107 DEGs between most similar clusters, FDR < 0.01). To generate the subsetted astrocyte dataset (as in Figure 1D), cells from cluster 0 (res. 0.4) were extracted and run through the pipeline. Clusters were assigned at a resolution of 0.2 (4 clusters identified with at least 112 DEGs between most similar clusters, FDR <0.01). For the hiNS(-/+) in control, 3w and 5w Aβ merged dataset (as in Figure 6A), clusters were assigned at a resolution of 0.4 (16 clusters identified with at least 204 DEGs between most similar clusters, FDR < 0.01). To generate the subsetted astrocyte dataset (as in Figure 6C), cells from cluster 0 (res 0.4) were extracted and run through the pipeline. Clusters were assigned at a resolution of 0.2 (4 clusters identified with at least 112 DEGs between most similar clusters, FDR <0.01). For subsequent subsetted data sets we used resolutions of 0.4 for microglia/macrophages (as in Figure 7A), for neurons 0.2 (as in Figure S8A) and 0.1 for the hiNS(+) only merge of all six experimental groups (as in Figure 8E).

### DEG statistical analysis

DEG analysis of snRNA-seq data was performed using the FindMarkers function in Seurat. Expression levels were tested using a Wilcoxon Rank Sum Test. A Bonferroni-adjusted p-value smaller than 0.05 (padj < 0.05), coupled with an average fold-change of greater than 1.3 (FC > 1.3), was considered statistically significant.

### Gene ontology analysis

g:Profiler (version e109_eg56_p17_1d3191d, https://biit.cs.ut.ee/gprofiler/)[152] was used to determine which gene ontology (GO) terms were significantly differentially overrepresented in cell populations following gene expression (fold-change > 1.3, adjusted p-value < 0.05) comparisons. If multiple Ensembl IDs were detected for a particular gene, the ID with the most GO annotations was used. In cases with multiple IDs with the same number of GO annotations, the first ID was used. Some genes were excluded from analysis if Ensembl IDs could not be found. The g:SCS multiple testing correction algorithm was used for p-value correction with a significance threshold of 0.05.

### Ligand-receptor interaction analysis

For ligand-receptor interaction analysis between differentially expressed ligands with differentially expressed receptors on astrocytes and neurons we used our 5w Aβ hiNS(+) snRNA-seq group and subsetted from the merged Seurat including all treatment groups, and clustered with 0.4 resolution. The clusters that represented microglia/macrophages (13), astrocytes (0), and excitatory and inhibitory neurons (5, 4, 2, 7, 12, 10, 14, 3, 9, 8) were isolated for ligand-receptor analysis. Genes for microglia ligands were matched with their astrocyte and neuron receptor genes using the LR network database from Zenodo (https://zenodo.org/record/7074291/files/lr_network_human_21122021.rds). For each ligand or receptor, to determine if the gene was specific to its cluster, differential gene analysis was done with the Seurat function FindAllMarkers. Any genes with a Bonferroni-adjusted p-value smaller than 0.05 (padj < 0.05) and an average logfold-change of greater than 1.3 (log2FC > 1.3) were considered cluster specific. Only ligand-receptor pairs where both genes were cluster specific and had a mean percent expression greater than 10% were plotted with cc_circos in the CCPlotR package.

### Pearson and single-cell correlation analyses

Pearson correlation analysis (Figure 1H) was performed by averaging the expression of a given transcript within each cluster/cell type of interest and correlating the expression values between clusters/cell types using the cor.test function in R.

### Statistical analysis

For the comparison of calcium wave frequencies (quantified by GCaMP6f imaging), paired student-t-tests were used for pharmacological experiments and comparing two different time points. Unpaired student t-tests were used when comparing baseline calcium wave frequencies, Aβ phagocytosis, immunoreactivity and nuclei density of control hiNS vs. Aβ treated hiNS. For comparing microglial densities and GFAP immunoreactivity between control, 3w Aβ and 5w Aβ hiNS(+) ordinary one-way ANOVA followed by Holm-Sidak post-hoc tests were used. Logarithmic transformation was performed on all data sets comparing control, 3w Aβ and 5w Aβ hiNS with or without hiMG prior to using two-way ANOVAs, testing for the effect of hiMG, followed by Holm-Sidak post-hoc tests. Statistical significance was indicated as follows: * p < 0.05; ** p < 0.01; *** p < 0.001; **** p < 0.0001.

## Figures legends

**Figure S1:**
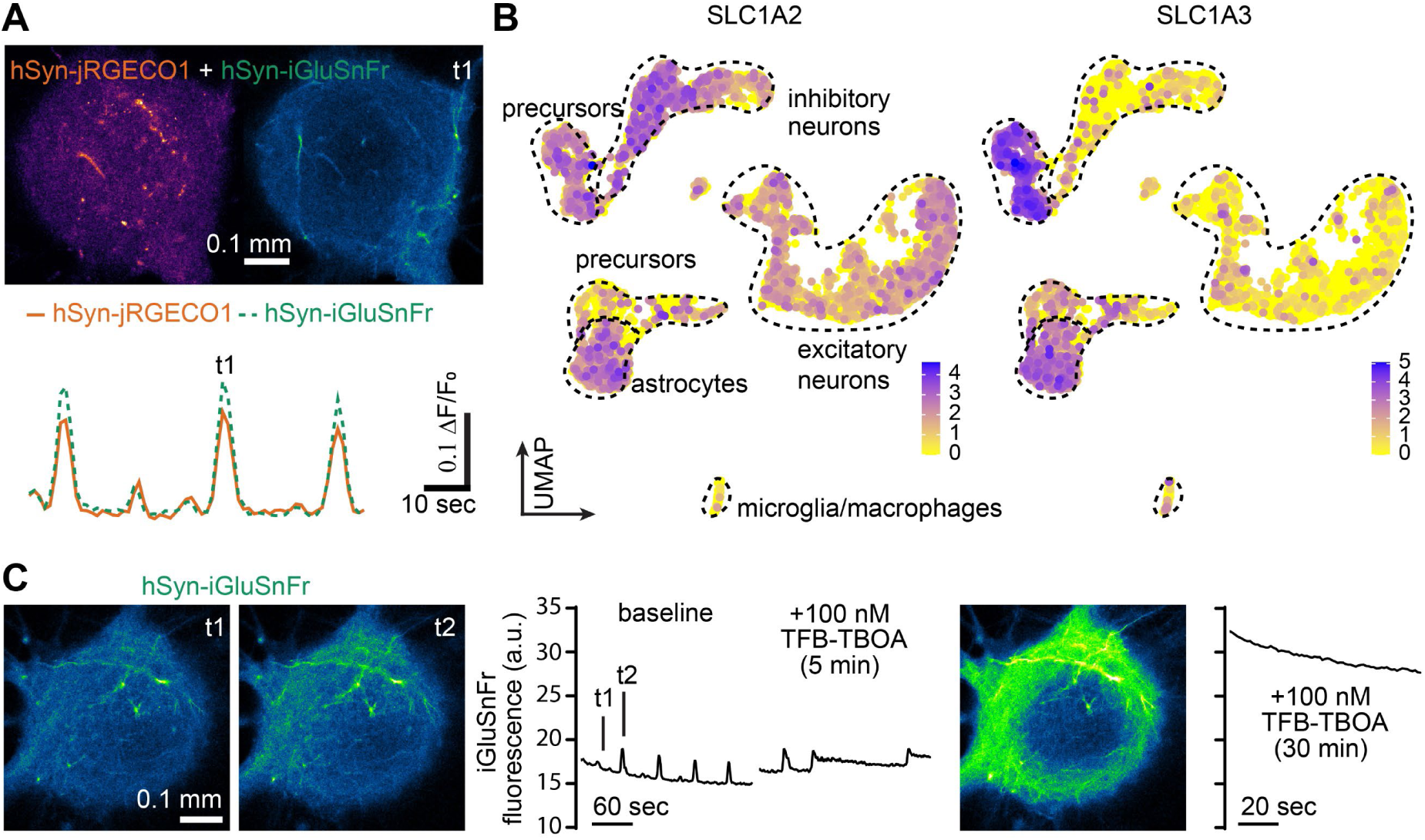
Extracellular glutamate elevations coincide with calcium waves and maintaining calcium wave activity requires glutamate transporter activity in hiNS. **A**: Elevated extracellular glutamate (detected by *hSyn*-iGluSnFr expression) during calcium wave activity in hiNS (detected by *hSyn*-jRECO1 expression). **B**: Glutamate transporter EAAT2 (SLC1A2) and EAAT1 (SLC1A3) expression in hiNS. While SLC1A2 expression is widespread in neurons, progenitors and astrocytes, SLC1A3 expression appears more specific for progenitor and astrocyte cell populations. **C**: iGluSnFr fluorescence, indicating extracellular glutamate levels, gradually increase by blocking EAAT1/2 with 100 nM TFB-TBOA (N=3) which prevents wave formation within minutes in hiNS indicating a functional role of glutamate transporters in maintaining calcium/glutamate wave activity.

**Figure S2:**
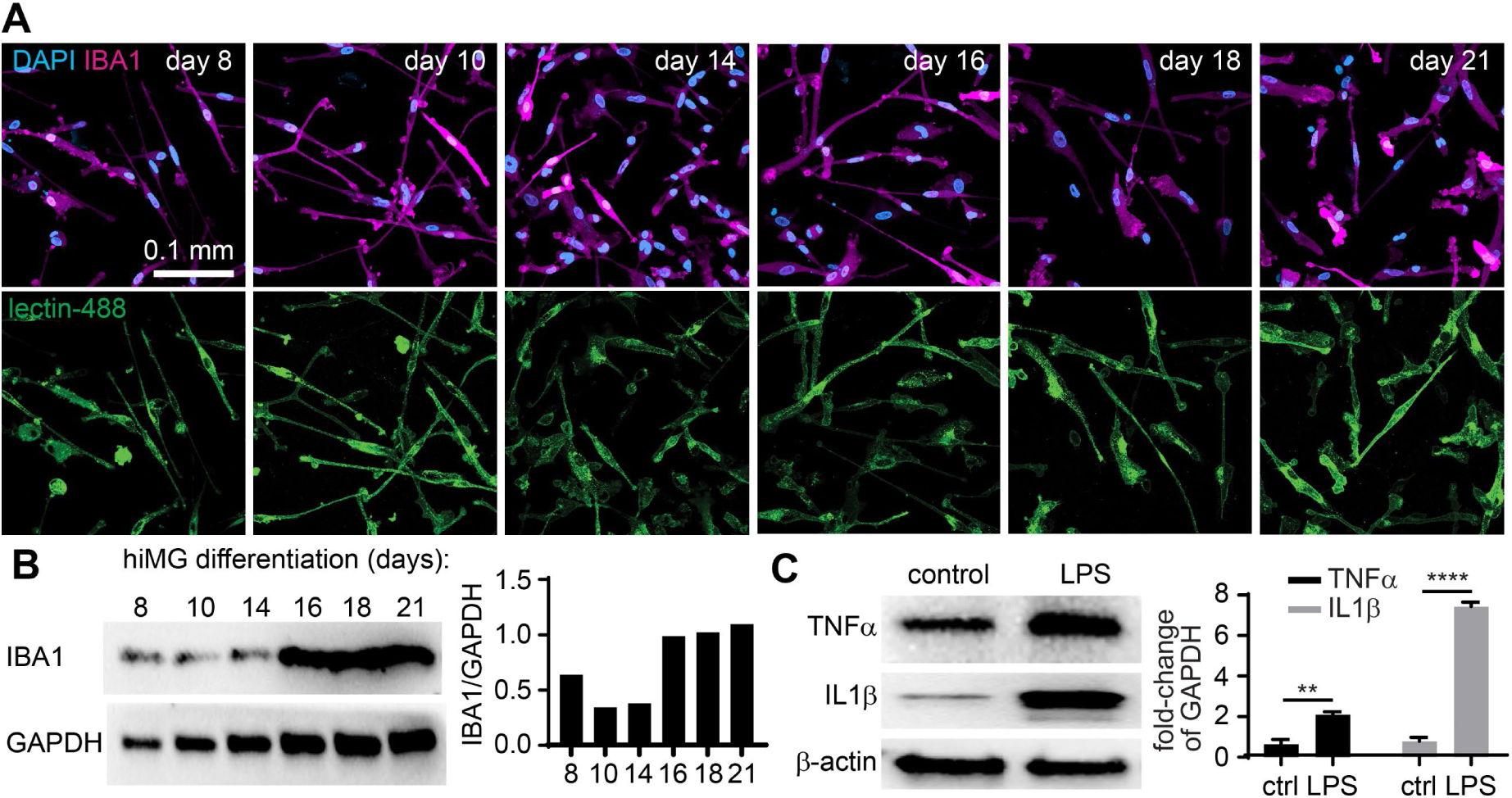
hiMG differentiation timeline shows a gradual increase in IBA1 expression. **A**: hiMG were plated on coverslips and fixed at different time points during differentiation (8 to 21 days) (N=3 coverslips each time point). Immunofluorescence for IBA1 (magenta) is detectable at all time points. Co-staining with tomato lectin-488 confirms that lectin can be used to label hiMG *in vitro*. **B**: Western blot for IBA1 confirms stable expression in hiMG up to at least day 21 (N=3 each). **C**: Stimulation of hiMG with LPS for 24h results in significant TNFα and IL1β release confirming their capacity to react to proinflammatory stimuli (N=3 each).

**Figure S3:**
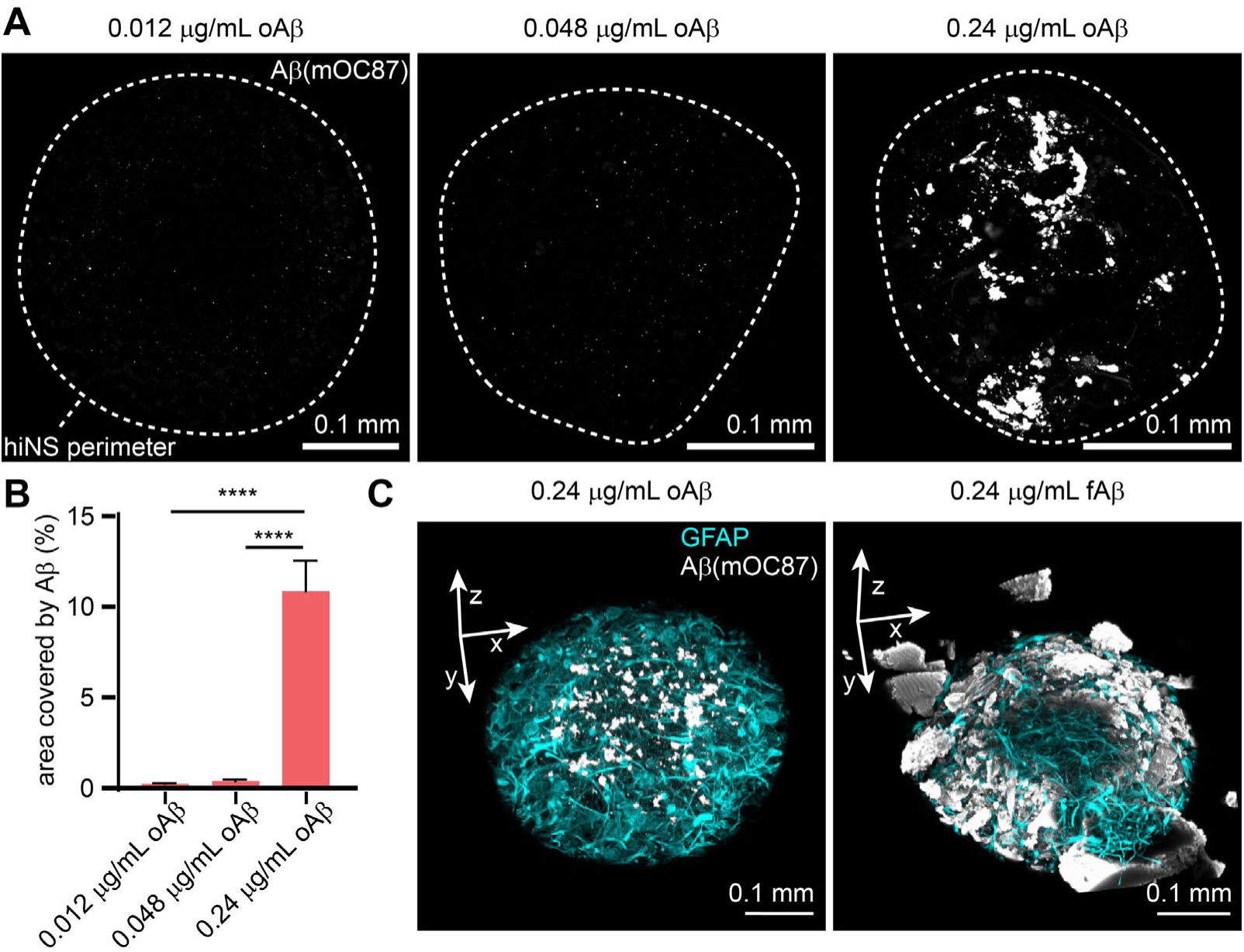
Aβ aggregates form within 7 days of oligomeric Ab (oAβ) treatment in hiNS. **A**: hiNS cell culture medium was supplemented with three differend doses of oAβ. Only the highest (0.24 μg/mL) concentration resulted in robust formation of aggregated Aβ (inside the tissue **B**: Quantification of mOC87 (fibril specific anti-Aβ antibody) positive area in hiNS. One-way ANOVA followed by Holm-Sidak’s post-hoc test **C**: Three-dimensional reconstruction of Aβ aggregates in hiNS. While oAβ treatment results in plaque-like aggregates inside the tissue (left image), supplementing cell culture media with pre-aggregated fibrillar Aβ (fAβ) results in large Aβ sheets covering the hiNS tissue which do not mimic plaque-like structures (right image).

**Figure S4:**
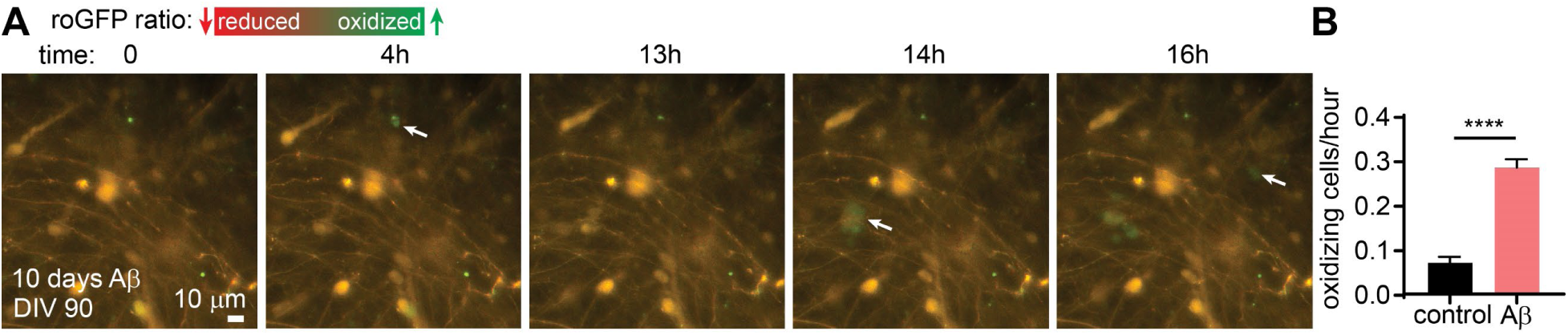
Over-night live confocal imaging of *hSyn* -roGFP1 expressing hiNS(-) during chronic Aβ treatment. Recordings of 18-22h duration were conducted between DIV 87-104 with chronic Aβ treatment ranging from 7 to 23 days (N=5 over-night recordings each). **A**: Example section of an Aβ treated hiNS displaying multiple neurons oxidizing during the recording time period (see arrows). **B**: The total number of oxidizing cells per hour was quantified for both Aβ treated as well as control hiNS. In Aβ treated spheres 0.29 ± 0.02 cells oxidized per hour compared to 0.07 ± 0.01 under control conditions (student’s t-test, **** p<0.0001).

**Figure S5:**
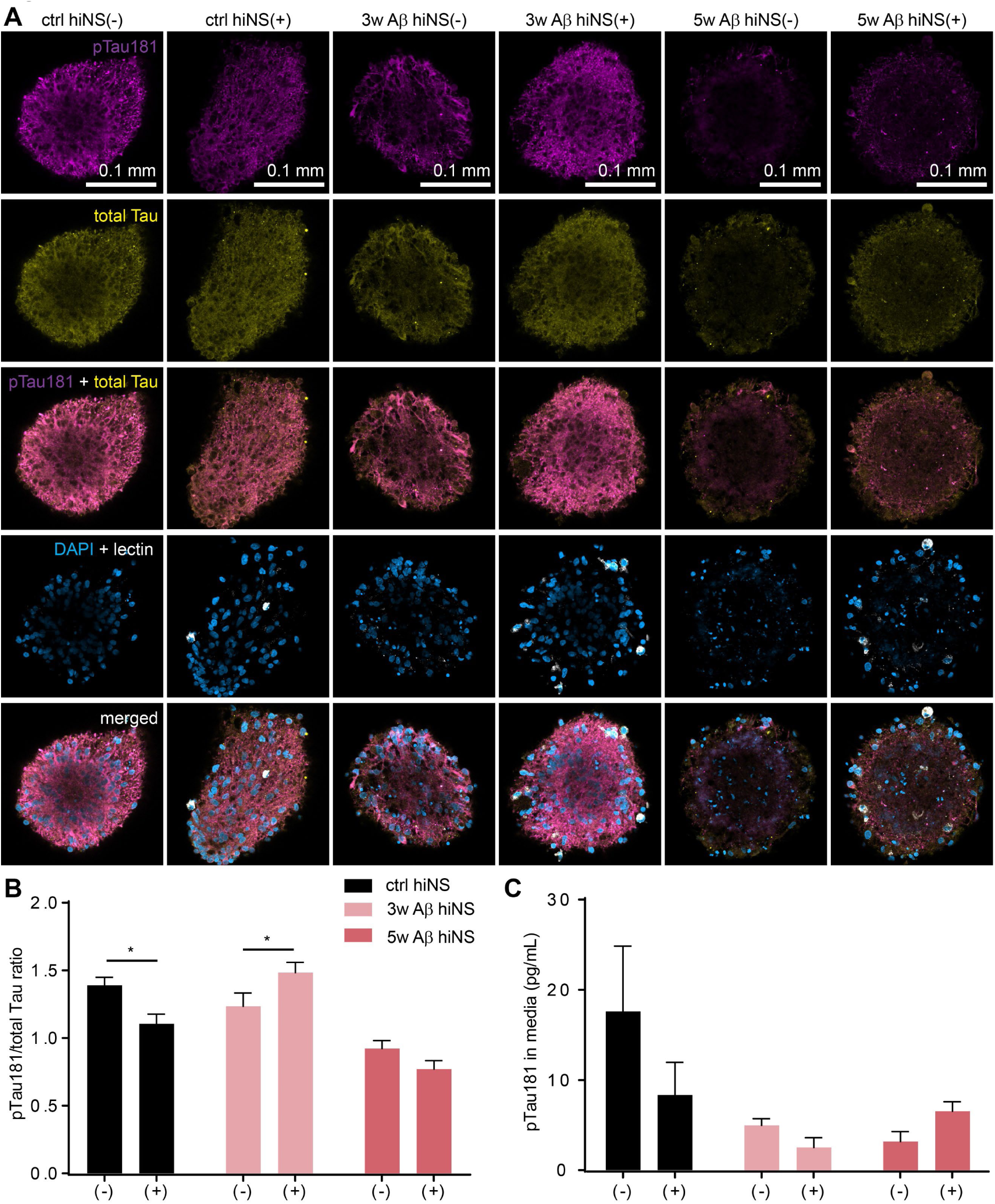
Chronic amyloidosis does not result in Aβ induced Tau hyperphosphorylation. **A**: Immunofluorescence staining for phosphorylated Tau T181 (pTau181) and total Tau in hiNS of all 6 experimental conditions. **B**: Quantification of pTau181/total Tau ratio in hiNS indicates pTau modulation by hiMG independent of Aβ exposure. **C**: Soluble pTau181 was quantified using ELISA on supernatants from hiNS (N=3; pooled supernatant from six individual hiNS each) confirms the lack of Aβ induced Tau hyperphosphorylation.

**Figure S6:**
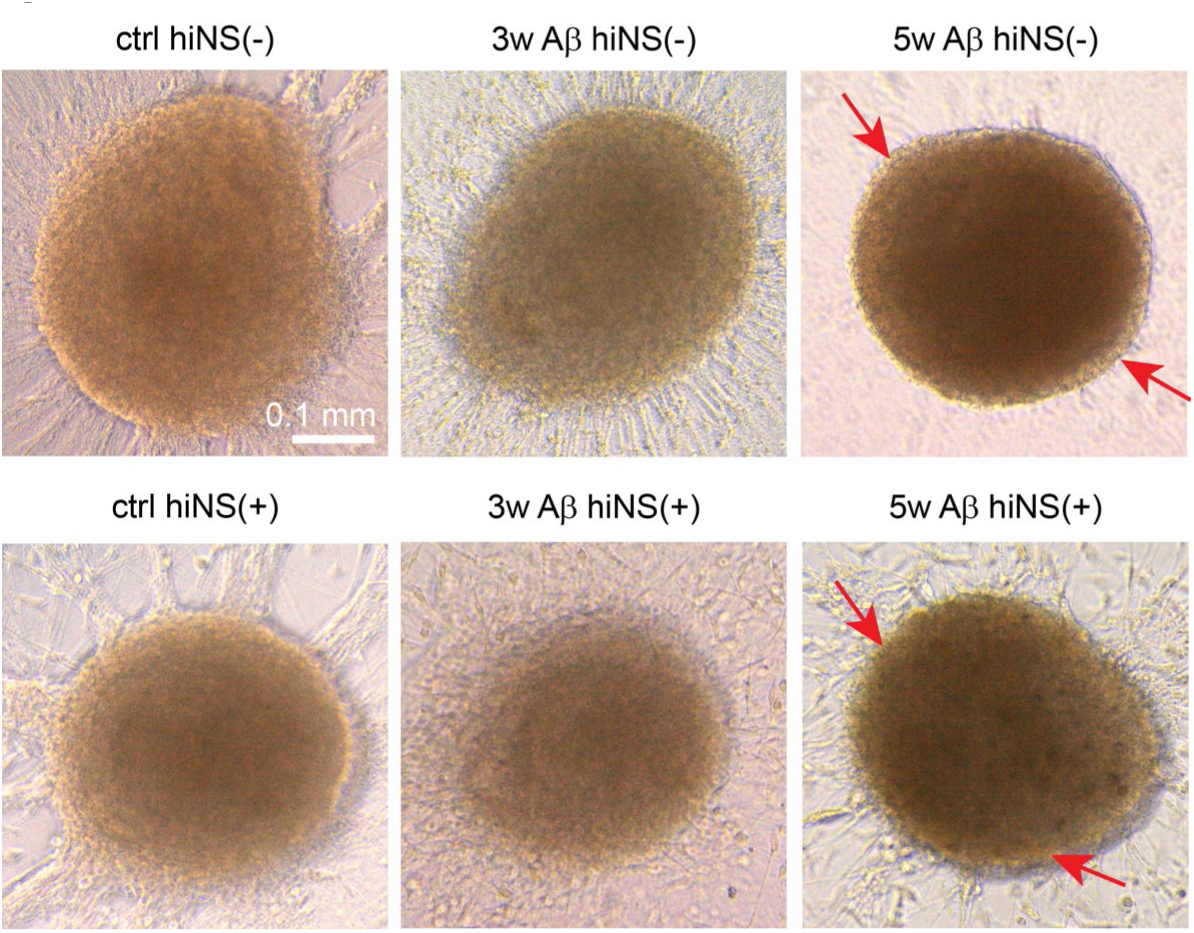
Example transmitted light images of control and Aβ treated hiNS with and without hiMG. hiMG attach to the surface surrounding hiNS Note that 5w Aβ hiNS (+/-) display a darker contrast and appear to detach from the surface (see red arrows).

**Figure S7:**
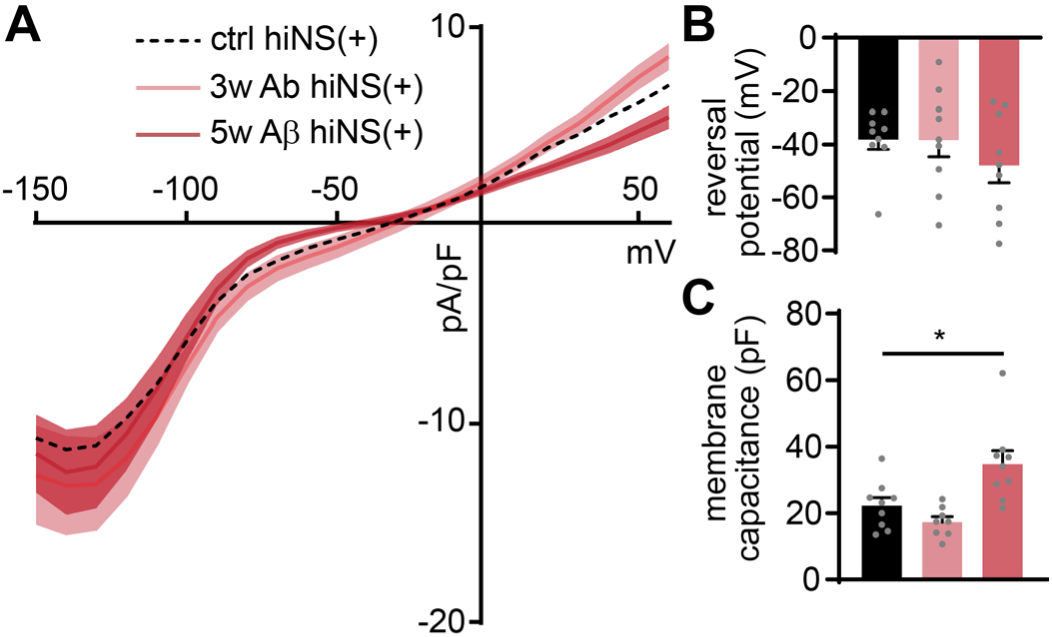
Electrophysiological properties of hiMG in 3w and 5w Aβ hiNS(+). **A**: Current-voltage relationship of hiMG ranging from -150 to +50 mV after chronic Aβ treatment appears similar to hiMG from ctrl hiNS(+) with no significant increase in outward or inward currents. **B**: Reversal potentials of hiMG do not differ between treatment groups indicating membrane potentials of ∼-40 mV. **c**: Membrane capacitance of hiMG was significantly increased in hiMG of 5w Aβ hiNS(+) indicating potentially larger cell sizes with prolonged Aβ treatment (ctrl, 3w Aβ, 5w Aβ hiNS(+): N=9 each).

**Figure S8:**
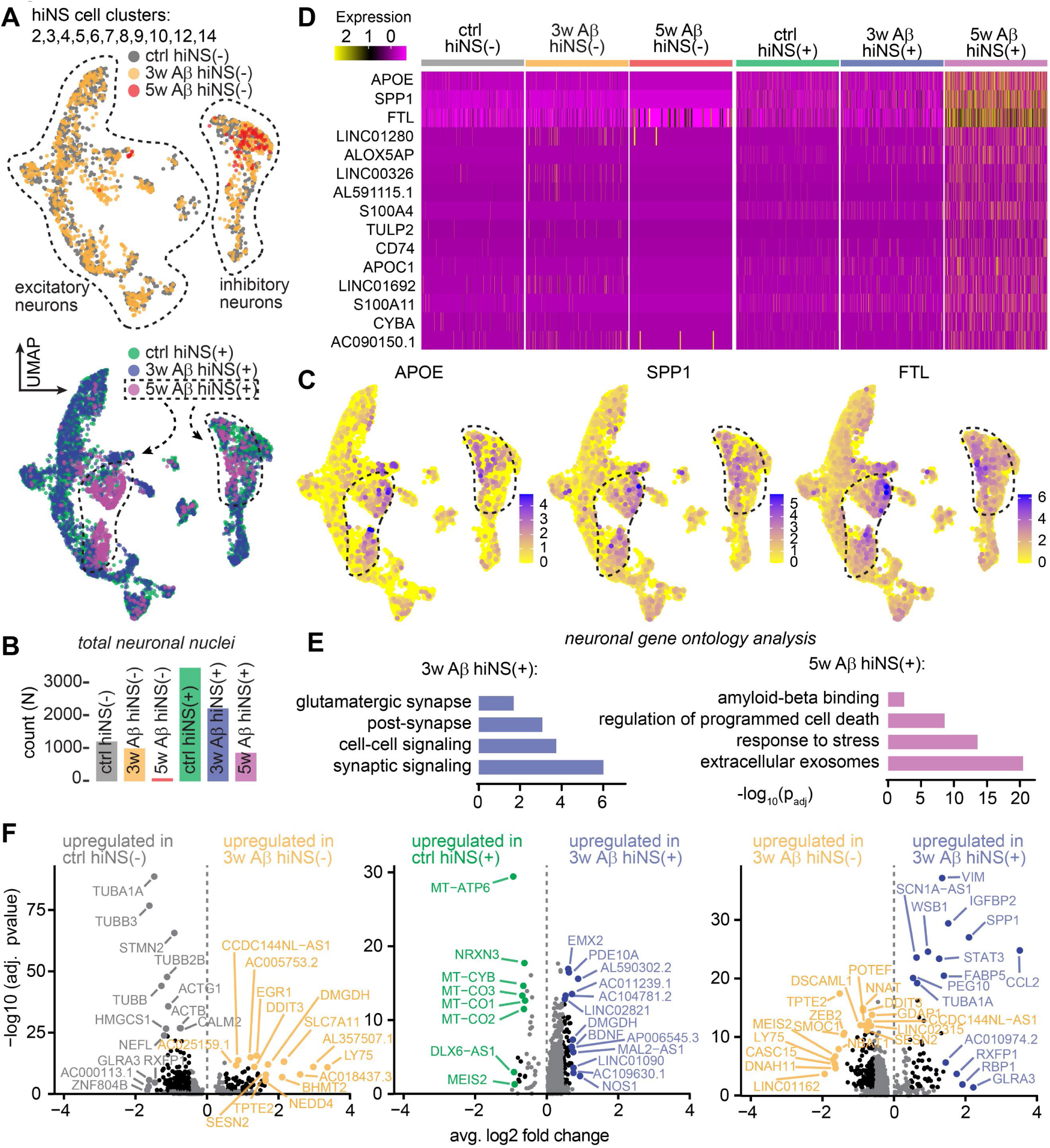
Neuronal gene expression changes during chronic amyloidosis in the presence of hiMG include *APOE*. **A**: Isolated neuronal cell populations from snRNA-seq on hiNS(-/+) in ctrl, 3w and 5w Aβ conditions. Merged UMAPs of all 6 experimental groups displaying excitatory neurons (left) and inhibitory neurons (right). Note that the majority of 5w Aβ hiNS(+) neurons cluster together in the individual neuronal populations. **B**: Quantification of total neuronal nuclei count in all 6 experimental groups. Note that only few neurons remain after chronic amyloidosis in the absence of hiMG. **C**: UMAP overlay of the three most significant DEGs *APOE, SPP1* and *FTL*. Cell populations enriched with 5w Aβ hiNS(+) neurons are encircled in dashed black lines. **D**: Heatmap depicting the top 15 most significant DEGs in 5w Aβ hiNS(+) neurons, with *APOE* being the most significant upregulated gene. **E**: Neuronal gene ontology analysis in 3w/5w Aβ hiNS(+) neurons. **F**: Direct comparison of neuronal DEGs depending on Ab or hiMG for ctrl and 3w Aβ hiNS shown in volcano plots (compared in pairs). Top 10 DEGs in respect to fold change and/or adjusted pvalue are labeled. Note that AD-associated genes, such as *CCL2*, *STAT3* and *VIM*, are only upregulated if hiMG were present during Aβ treatment.

**Figure S9:**
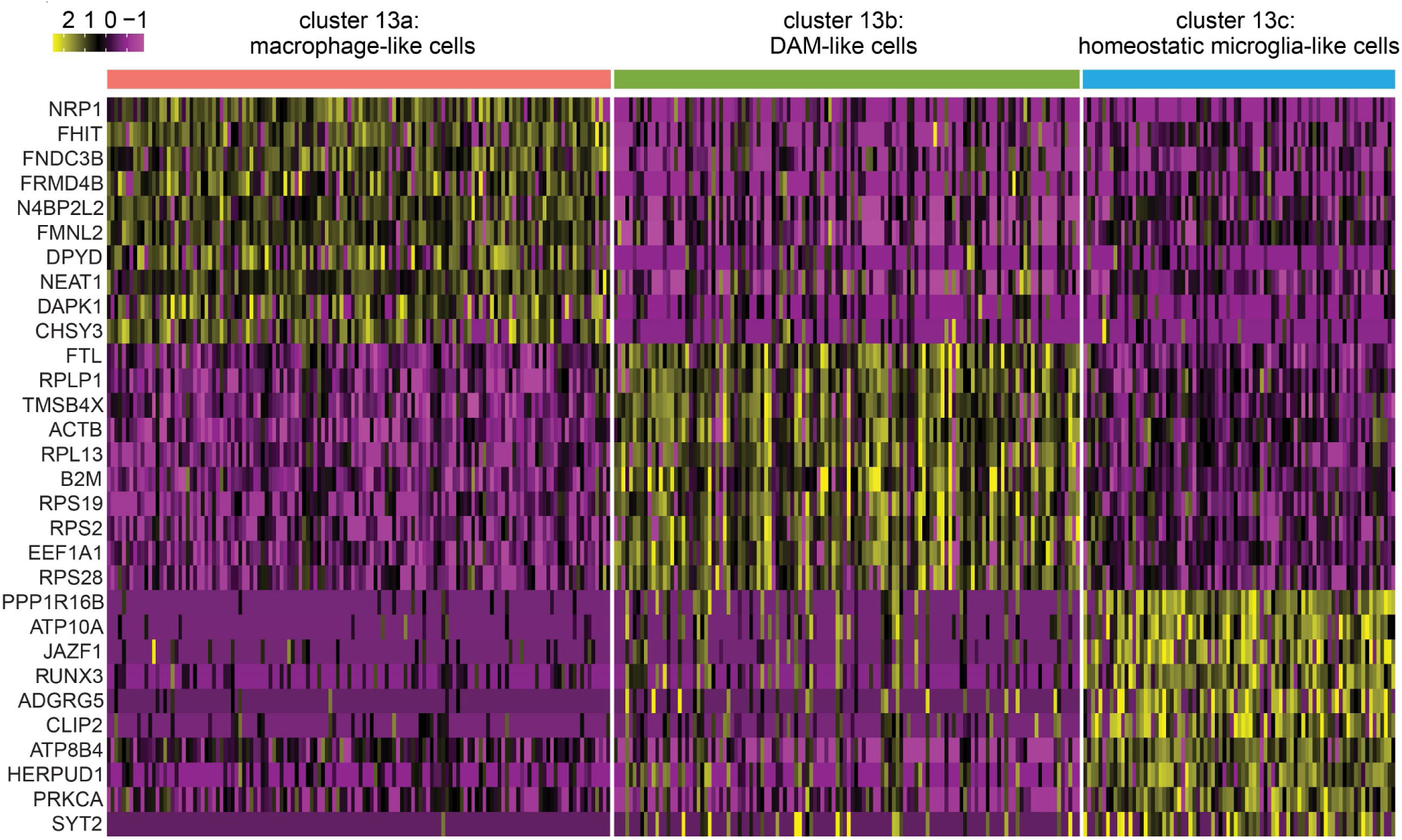
Top 10 most significant DEGs per subcluster of hiMG from 5w Aβ hiNS(+) displayed in a heatmap.

## References

1. Guerreiro R, Wojtas A, Bras J, Carrasquillo M, Rogaeva E, Majounie E, Cruchaga C, Sassi C, Kauwe JS, Younkin S, et al: TREM2 variants in Alzheimer’s disease. N Engl J Med 2013, 368:117–127.

2. Zhang Y, Chen H, Li R, Sterling K, Song W: Amyloid beta-based therapy for Alzheimer’s disease: challenges, successes and future. Signal Transduct Target Ther 2023, 8:248.

3. Leisher S, Bohorquez A, Gay M, Garcia V, Jones R, Baldaranov D, Rafii MS: Amyloid-Lowering Monoclonal Antibodies for the Treatment of Early Alzheimer’s Disease. CNS Drugs 2023, 37:671–677.

4. Piccioni G, Mango D, Saidi A, Corbo M, Nistico R: Targeting Microglia-Synapse Interactions in Alzheimer’s Disease. Int J Mol Sci 2021, 22.

5. Grubman A, Chew G, Ouyang JF, Sun G, Choo XY, McLean C, Simmons RK, Buckberry S, Vargas-Landin DB, Poppe D, et al: A single-cell atlas of entorhinal cortex from individuals with Alzheimer’s disease reveals cell-type-specific gene expression regulation. Nat Neurosci 2019, 22:2087–2097.

6. Sadick JS, O’Dea MR, Hasel P, Dykstra T, Faustin A, Liddelow SA: Astrocytes and oligodendrocytes undergo subtype-specific transcriptional changes in Alzheimer’s disease. Neuron 2022, 110:1788–1805 e1710.

7. Gao C, Shen X, Tan Y, Chen S: Pathogenesis, therapeutic strategies and biomarker development based on "omics" analysis related to microglia in Alzheimer’s disease. J Neuroinflammation 2022, 19:215.

8. Drummond E, Wisniewski T: Alzheimer’s disease: experimental models and reality. Acta Neuropathol 2017, 133:155–175.

9. Barak M, Fedorova V, Pospisilova V, Raska J, Vochyanova S, Sedmik J, Hribkova H, Klimova H, Vanova T, Bohaciakova D: Human iPSC-Derived Neural Models for Studying Alzheimer’s Disease: from Neural Stem Cells to Cerebral Organoids. Stem Cell Rev Rep 2022, 18:792–820.

10. Papaspyropoulos A, Tsolaki M, Foroglou N, Pantazaki AA: Modeling and Targeting Alzheimer’s Disease With Organoids. Front Pharmacol 2020, 11:396.

11. Sreenivasamurthy S, Laul M, Zhao N, Kim T, Zhu D: Current progress of cerebral organoids for modeling Alzheimer’s disease origins and mechanisms. Bioeng Transl Med 2023, 8:e10378.

12. Choi SH, Kim YH, Quinti L, Tanzi RE, Kim DY: 3D culture models of Alzheimer’s disease: a road map to a "cure-in-a-dish". Mol Neurodegener 2016, 11:75.

13. Hernandez-Sapiens MA, Reza-Zaldivar EE, Cevallos RR, Marquez-Aguirre AL, Gazarian K, Canales-Aguirre AA: A Three-Dimensional Alzheimer’s Disease Cell Culture Model Using iPSC-Derived Neurons Carrying A246E Mutation in PSEN1. Front Cell Neurosci 2020, 14:151.

14. Jorfi M, Park J, Hall CK, Lin CJ, Chen M, von Maydell D, Kruskop JM, Kang B, Choi Y, Prokopenko D, et al: Infiltrating CD8(+) T cells exacerbate Alzheimer’s disease pathology in a 3D human neuroimmune axis model. Nat Neurosci 2023, 26:1489–1504.

15. Kim YH, Choi SH, D’Avanzo C, Hebisch M, Sliwinski C, Bylykbashi E, Washicosky KJ, Klee JB, Brustle O, Tanzi RE, Kim DY: A 3D human neural cell culture system for modeling Alzheimer’s disease. Nat Protoc 2015, 10:985–1006.

16. Kwak SS, Washicosky KJ, Brand E, von Maydell D, Aronson J, Kim S, Capen DE, Cetinbas M, Sadreyev R, Ning S, et al: Amyloid-beta42/40 ratio drives tau pathology in 3D human neural cell culture models of Alzheimer’s disease. Nat Commun 2020, 11:1377.

17. Poli D, Magliaro C, Ahluwalia A: Experimental and Computational Methods for the Study of Cerebral Organoids: A Review. Front Neurosci 2019, 13:162.

18. Hanson GT, Aggeler R, Oglesbee D, Cannon M, Capaldi RA, Tsien RY, Remington SJ: Investigating mitochondrial redox potential with redox-sensitive green fluorescent protein indicators. J Biol Chem 2004, 279:13044–13053.

19. Wendt S, Johnson S, Weilinger NL, Groten C, Sorrentino S, Frew J, Yang L, Choi HB, Nygaard HB, MacVicar BA: Simultaneous imaging of redox states in dystrophic neurites and microglia at Abeta plaques indicate lysosome accumulation not microglia correlate with increased oxidative stress. Redox Biol 2022, 56:102448.

20. Hao Y, Hao S, Andersen-Nissen E, Mauck WM, 3rd, Zheng S, Butler A, Lee MJ, Wilk AJ, Darby C, Zager M, et al: Integrated analysis of multimodal single-cell data. Cell 2021, 184:3573–3587 e3529.

21. Lun AT, McCarthy DJ, Marioni JC: A step-by-step workflow for low-level analysis of single-cell RNA-seq data with Bioconductor. F1000Res 2016, 5:2122.

22. Yuzwa SA, Borrett MJ, Innes BT, Voronova A, Ketela T, Kaplan DR, Bader GD, Miller FD: Developmental Emergence of Adult Neural Stem Cells as Revealed by Single-Cell Transcriptional Profiling. Cell Rep 2017, 21:3970–3986.

23. Borrett MJ, Innes BT, Tahmasian N, Bader GD, Kaplan DR, Miller FD: A Shared Transcriptional Identity for Forebrain and Dentate Gyrus Neural Stem Cells from Embryogenesis to Adulthood. eNeuro 2022, 9.

24. Borrett MJ, Innes BT, Jeong D, Tahmasian N, Storer MA, Bader GD, Kaplan DR, Miller FD: Single-Cell Profiling Shows Murine Forebrain Neural Stem Cells Reacquire a Developmental State when Activated for Adult Neurogenesis. Cell Rep 2020, 32:108022.

25. Arjun McKinney A, Petrova R, Panagiotakos G: Calcium and activity-dependent signaling in the developing cerebral cortex. Development 2022, 149.

26. Namiki S, Norimoto H, Kobayashi C, Nakatani K, Matsuki N, Ikegaya Y: Layer III neurons control synchronized waves in the immature cerebral cortex. J Neurosci 2013, 33:987–1001.

27. Uwechue NM, Marx MC, Chevy Q, Billups B: Activation of glutamate transport evokes rapid glutamine release from perisynaptic astrocytes. J Physiol 2012, 590:2317–2331.

28. Fellin T, Pascual O, Gobbo S, Pozzan T, Haydon PG, Carmignoto G: Neuronal synchrony mediated by astrocytic glutamate through activation of extrasynaptic NMDA receptors. Neuron 2004, 43:729–743.

29. Boucsein C, Zacharias R, Farber K, Pavlovic S, Hanisch UK, Kettenmann H: Purinergic receptors on microglial cells: functional expression in acute brain slices and modulation of microglial activation in vitro. Eur J Neurosci 2003, 17:2267–2276.

30. Wendt S, Maricos M, Vana N, Meyer N, Guneykaya D, Semtner M, Kettenmann H: Changes in phagocytosis and potassium channel activity in microglia of 5xFAD mice indicate alterations in purinergic signaling in a mouse model of Alzheimer’s disease. Neurobiol Aging 2017, 58:41–53.

31. Wendt S, Wogram E, Korvers L, Kettenmann H: Experimental Cortical Spreading Depression Induces NMDA Receptor Dependent Potassium Currents in Microglia. J Neurosci 2016, 36:6165–6174.

32. Haas LT, Kostylev MA, Strittmatter SM: Therapeutic molecules and endogenous ligands regulate the interaction between brain cellular prion protein (PrPC) and metabotropic glutamate receptor 5 (mGluR5). J Biol Chem 2014, 289:28460–28477.

33. Um JW, Kaufman AC, Kostylev M, Heiss JK, Stagi M, Takahashi H, Kerrisk ME, Vortmeyer A, Wisniewski T, Koleske AJ, et al: Metabotropic glutamate receptor 5 is a coreceptor for Alzheimer abeta oligomer bound to cellular prion protein. Neuron 2013, 79:887–902.

34. Um JW, Nygaard HB, Heiss JK, Kostylev MA, Stagi M, Vortmeyer A, Wisniewski T, Gunther EC, Strittmatter SM: Alzheimer amyloid-beta oligomer bound to postsynaptic prion protein activates Fyn to impair neurons. Nat Neurosci 2012, 15:1227–1235.

35. Lauren J, Gimbel DA, Nygaard HB, Gilbert JW, Strittmatter SM: Cellular prion protein mediates impairment of synaptic plasticity by amyloid-beta oligomers. Nature 2009, 457:1128–1132.

36. Kaufman AC, Salazar SV, Haas LT, Yang J, Kostylev MA, Jeng AT, Robinson SA, Gunther EC, van Dyck CH, Nygaard HB, Strittmatter SM: Fyn inhibition rescues established memory and synapse loss in Alzheimer mice. Ann Neurol 2015, 77:953–971.

37. Zhang G, Wang Z, Hu H, Zhao M, Sun L: Microglia in Alzheimer’s Disease: A Target for Therapeutic Intervention. Front Cell Neurosci 2021, 15:749587.

38. Gao C, Jiang J, Tan Y, Chen S: Microglia in neurodegenerative diseases: mechanism and potential therapeutic targets. Signal Transduct Target Ther 2023, 8:359.

39. Hatami A, Albay R, 3rd, Monjazeb S, Milton S, Glabe C: Monoclonal antibodies against Abeta42 fibrils distinguish multiple aggregation state polymorphisms in vitro and in Alzheimer disease brain. J Biol Chem 2014, 289:32131–32143.

40. Jurga AM, Paleczna M, Kuter KZ: Overview of General and Discriminating Markers of Differential Microglia Phenotypes. Front Cell Neurosci 2020, 14:198.

41. Jung H, Lee SY, Lim S, Choi HR, Choi Y, Kim M, Kim S, Lee Y, Han KH, Chung WS, Kim CH: Anti-inflammatory clearance of amyloid-beta by a chimeric Gas6 fusion protein. Nat Med 2022, 28:1802–1812.

42. Busche MA, Hyman BT: Synergy between amyloid-beta and tau in Alzheimer’s disease. Nat Neurosci 2020, 23:1183–1193.

43. Mathys H, Boix CA, Akay LA, Xia Z, Davila-Velderrain J, Ng AP, Jiang X, Abdelhady G, Galani K, Mantero J, et al: Single-cell multiregion dissection of Alzheimer’s disease. Nature 2024, 632:858–868.

44. DeChellis-Marks MR, Wei Y, Ding Y, Wolfe CM, Krivinko JM, MacDonald ML, Lopez OL, Sweet RA, Kofler J: Psychosis in Alzheimer’s Disease Is Associated With Increased Excitatory Neuron Vulnerability and Post-transcriptional Mechanisms Altering Synaptic Protein Levels. Front Neurol 2022, 13:778419.

45. Keren-Shaul H, Spinrad A, Weiner A, Matcovitch-Natan O, Dvir-Szternfeld R, Ulland TK, David E, Baruch K, Lara-Astaiso D, Toth B, et al: A Unique Microglia Type Associated with Restricting Development of Alzheimer’s Disease. Cell 2017, 169:1276–1290 e1217.

46. Raulin AC, Doss SV, Trottier ZA, Ikezu TC, Bu G, Liu CC: ApoE in Alzheimer’s disease: pathophysiology and therapeutic strategies. Mol Neurodegener 2022, 17:72.

47. Strittmatter WJ, Saunders AM, Schmechel D, Pericak-Vance M, Enghild J, Salvesen GS, Roses AD: Apolipoprotein E: high-avidity binding to beta-amyloid and increased frequency of type 4 allele in late-onset familial Alzheimer disease. Proc Natl Acad Sci U S A 1993, 90:1977–1981.

48. Delhaye M, LeDue J, Robinson K, Xu Q, Zhang Q, Oku S, Zhang P, Craig AM: Adaptation of Magnified Analysis of the Proteome for Excitatory Synaptic Proteins in Varied Samples and Evaluation of Cell Type-Specific Distributions. J Neurosci 2024, 44.

49. Kuga N, Sasaki T, Takahara Y, Matsuki N, Ikegaya Y: Large-scale calcium waves traveling through astrocytic networks in vivo. J Neurosci 2011, 31:2607–2614.

50. Yuryev M, Pellegrino C, Jokinen V, Andriichuk L, Khirug S, Khiroug L, Rivera C: In vivo Calcium Imaging of Evoked Calcium Waves in the Embryonic Cortex. Front Cell Neurosci 2015, 9:500.

51. Xiang Y, Tanaka Y, Patterson B, Kang YJ, Govindaiah G, Roselaar N, Cakir B, Kim KY, Lombroso AP, Hwang SM, et al: Fusion of Regionally Specified hPSC-Derived Organoids Models Human Brain Development and Interneuron Migration. Cell Stem Cell 2017, 21:383–398 e387.

52. Woodruff G, Phillips N, Carromeu C, Guicherit O, White A, Johnson M, Zanella F, Anson B, Lovenberg T, Bonaventure P, Harrington AW: Screening for modulators of neural network activity in 3D human iPSC-derived cortical spheroids. PLoS One 2020, 15:e0240991.

53. Michell-Robinson MA, Touil H, Healy LM, Owen DR, Durafourt BA, Bar-Or A, Antel JP, Moore CS: Roles of microglia in brain development, tissue maintenance and repair. Brain 2015, 138:1138–1159.

54. Park J, Wetzel I, Marriott I, Dreau D, D’Avanzo C, Kim DY, Tanzi RE, Cho H: A 3D human triculture system modeling neurodegeneration and neuroinflammation in Alzheimer’s disease. Nat Neurosci 2018, 21:941–951.

55. Rifat A, Ossola B, Burli RW, Dawson LA, Brice NL, Rowland A, Lizio M, Xu X, Page K, Fidzinski P, et al: Differential contribution of THIK-1 K(+) channels and P2X7 receptors to ATP-mediated neuroinflammation by human microglia. J Neuroinflammation 2024, 21:58.

56. Golde TE: Alzheimer’s disease - the journey of a healthy brain into organ failure. Mol Neurodegener 2022, 17:18.

57. Misrani A, Tabassum S, Yang L: Mitochondrial Dysfunction and Oxidative Stress in Alzheimer’s Disease. Front Aging Neurosci 2021, 13:617588.

58. Huang WJ, Zhang X, Chen WW: Role of oxidative stress in Alzheimer’s disease. Biomed Rep 2016, 4:519–522.

59. Fracassi A, Marcatti M, Zolochevska O, Tabor N, Woltjer R, Moreno S, Taglialatela G: Oxidative Damage and Antioxidant Response in Frontal Cortex of Demented and Nondemented Individuals with Alzheimer’s Neuropathology. J Neurosci 2021, 41:538–554.

60. Palop JJ, Mucke L: Amyloid-beta-induced neuronal dysfunction in Alzheimer’s disease: from synapses toward neural networks. Nat Neurosci 2010, 13:812–818.

61. Louneva N, Cohen JW, Han LY, Talbot K, Wilson RS, Bennett DA, Trojanowski JQ, Arnold SE: Caspase-3 is enriched in postsynaptic densities and increased in Alzheimer’s disease. Am J Pathol 2008, 173:1488–1495.

62. Stadelmann C, Deckwerth TL, Srinivasan A, Bancher C, Bruck W, Jellinger K, Lassmann H: Activation of caspase-3 in single neurons and autophagic granules of granulovacuolar degeneration in Alzheimer’s disease. Evidence for apoptotic cell death. Am J Pathol 1999, 155:1459–1466.

63. Goel P, Chakrabarti S, Goel K, Bhutani K, Chopra T, Bali S: Neuronal cell death mechanisms in Alzheimer’s disease: An insight. Front Mol Neurosci 2022, 15:937133.

64. Hansen DV, Hanson JE, Sheng M: Microglia in Alzheimer’s disease. J Cell Biol 2018, 217:459–472.

65. Leng F, Edison P: Neuroinflammation and microglial activation in Alzheimer disease: where do we go from here? Nat Rev Neurol 2021, 17:157–172.

66. Fan Z, Brooks DJ, Okello A, Edison P: An early and late peak in microglial activation in Alzheimer’s disease trajectory. Brain 2017, 140:792–803.

67. Solito E, Sastre M: Microglia function in Alzheimer’s disease. Front Pharmacol 2012, 3:14.

68. Wyss-Coray T: Inflammation in Alzheimer disease: driving force, bystander or beneficial response? Nat Med 2006, 12:1005–1015.

69. Long HZ, Zhou ZW, Cheng Y, Luo HY, Li FJ, Xu SG, Gao LC: The Role of Microglia in Alzheimer’s Disease From the Perspective of Immune Inflammation and Iron Metabolism. Front Aging Neurosci 2022, 14:888989.

70. Dolan MJ, Therrien M, Jereb S, Kamath T, Gazestani V, Atkeson T, Marsh SE, Goeva A, Lojek NM, Murphy S, et al: Exposure of iPSC-derived human microglia to brain substrates enables the generation and manipulation of diverse transcriptional states in vitro. Nat Immunol 2023, 24:1382–1390.

71. Abud EM, Ramirez RN, Martinez ES, Healy LM, Nguyen CHH, Newman SA, Yeromin AV, Scarfone VM, Marsh SE, Fimbres C, et al: iPSC-Derived Human Microglia-like Cells to Study Neurological Diseases. Neuron 2017, 94:278–293 e279.

72. Rossol M, Heine H, Meusch U, Quandt D, Klein C, Sweet MJ, Hauschildt S: LPS-induced cytokine production in human monocytes and macrophages. Crit Rev Immunol 2011, 31:379–446.

73. Erta M, Quintana A, Hidalgo J: Interleukin-6, a major cytokine in the central nervous system. Int J Biol Sci 2012, 8:1254–1266.

74. Feng Q, Wang YI, Yang Y: Neuroprotective effect of interleukin-6 in a rat model of cerebral ischemia. Exp Ther Med 2015, 9:1695–1701.

75. Andreadou M, Ingelfinger F, De Feo D, Cramer TLM, Tuzlak S, Friebel E, Schreiner B, Eede P, Schneeberger S, Geesdorf M, et al: IL-12 sensing in neurons induces neuroprotective CNS tissue adaptation and attenuates neuroinflammation in mice. Nat Neurosci 2023, 26:1701–1712.

76. Yang HS, Zhang C, Carlyle BC, Zhen SY, Trombetta BA, Schultz AP, Pruzin JJ, Fitzpatrick CD, Yau WW, Kirn DR, et al: Plasma IL-12/IFN-gamma axis predicts cognitive trajectories in cognitively unimpaired older adults. Alzheimers Dement 2022, 18:645–653.

77. Vom Berg J, Prokop S, Miller KR, Obst J, Kalin RE, Lopategui-Cabezas I, Wegner A, Mair F, Schipke CG, Peters O, et al: Inhibition of IL-12/IL-23 signaling reduces Alzheimer’s disease-like pathology and cognitive decline. Nat Med 2012, 18:1812–1819.

78. Takata M, Nishimura K, Harada K, Iwasaki R, Ando M, Yamada S, Ginhoux F, Takata K: Analysis of Abeta-induced neurotoxicity and microglial responses in simple two- and three-dimensional human iPSC-derived cortical culture systems. Tissue Cell 2023, 81:102023.

79. Alkhalifa AE, Al-Ghraiybah NF, Odum J, Shunnarah JG, Austin N, Kaddoumi A: Blood-Brain Barrier Breakdown in Alzheimer’s Disease: Mechanisms and Targeted Strategies. Int J Mol Sci 2023, 24.

80. Sun M, You H, Hu X, Luo Y, Zhang Z, Song Y, An J, Lu H: Microglia-Astrocyte Interaction in Neural Development and Neural Pathogenesis. Cells 2023, 12.

81. Matejuk A, Ransohoff RM: Crosstalk Between Astrocytes and Microglia: An Overview. Front Immunol 2020, 11:1416.

82. Chun H, Lee CJ: Reactive astrocytes in Alzheimer’s disease: A double-edged sword. Neurosci Res 2018, 126:44–52.

83. Shinohara M, Tachibana M, Kanekiyo T, Bu G: Role of LRP1 in the pathogenesis of Alzheimer’s disease: evidence from clinical and preclinical studies. J Lipid Res 2017, 58:1267–1281.

84. Liu CC, Hu J, Zhao N, Wang J, Wang N, Cirrito JR, Kanekiyo T, Holtzman DM, Bu G: Astrocytic LRP1 Mediates Brain Abeta Clearance and Impacts Amyloid Deposition. J Neurosci 2017, 37:4023–4031.

85. Habib N, McCabe C, Medina S, Varshavsky M, Kitsberg D, Dvir-Szternfeld R, Green G, Dionne D, Nguyen L, Marshall JL, et al: Disease-associated astrocytes in Alzheimer’s disease and aging. Nat Neurosci 2020, 23:701–706.

86. Foster EM, Dangla-Valls A, Lovestone S, Ribe EM, Buckley NJ: Clusterin in Alzheimer’s Disease: Mechanisms, Genetics, and Lessons From Other Pathologies. Front Neurosci 2019, 13:164.

87. Kamphuis W, Kooijman L, Orre M, Stassen O, Pekny M, Hol EM: GFAP and vimentin deficiency alters gene expression in astrocytes and microglia in wild-type mice and changes the transcriptional response of reactive glia in mouse model for Alzheimer’s disease. Glia 2015, 63:1036–1056.

88. Drummond E, Nayak S, Faustin A, Pires G, Hickman RA, Askenazi M, Cohen M, Haldiman T, Kim C, Han X, et al: Proteomic differences in amyloid plaques in rapidly progressive and sporadic Alzheimer’s disease. Acta Neuropathol 2017, 133:933–954.

89. Kim J, Yoo ID, Lim J, Moon JS: Pathological phenotypes of astrocytes in Alzheimer’s disease. Exp Mol Med 2024, 56:95–99.

90. Bakken TE, Hodge RD, Miller JA, Yao Z, Nguyen TN, Aevermann B, Barkan E, Bertagnolli D, Casper T, Dee N, et al: Single-nucleus and single-cell transcriptomes compared in matched cortical cell types. PLoS One 2018, 13:e0209648.

91. Mathys H, Davila-Velderrain J, Peng Z, Gao F, Mohammadi S, Young JZ, Menon M, He L, Abdurrob F, Jiang X, et al: Single-cell transcriptomic analysis of Alzheimer’s disease. Nature 2019, 570:332–337.

92. Kamphuis W, Middeldorp J, Kooijman L, Sluijs JA, Kooi EJ, Moeton M, Freriks M, Mizee MR, Hol EM: Glial fibrillary acidic protein isoform expression in plaque related astrogliosis in Alzheimer’s disease. Neurobiol Aging 2014, 35:492–510.

93. Jiwaji Z, Tiwari SS, Aviles-Reyes RX, Hooley M, Hampton D, Torvell M, Johnson DA, McQueen J, Baxter P, Sabari-Sankar K, et al: Reactive astrocytes acquire neuroprotective as well as deleterious signatures in response to Tau and Ass pathology. Nat Commun 2022, 13:135.

94. Ganne A, Balasubramaniam M, Griffin WST, Shmookler Reis RJ, Ayyadevara S: Glial Fibrillary Acidic Protein: A Biomarker and Drug Target for Alzheimer’s Disease. Pharmaceutics 2022, 14.

95. Dai DL, Li M, Lee EB: Human Alzheimer’s disease reactive astrocytes exhibit a loss of homeostastic gene expression. Acta Neuropathol Commun 2023, 11:127.

96. D’Ezio V, Colasanti M, Persichini T: Amyloid-beta 25-35 Induces Neurotoxicity through the Up-Regulation of Astrocytic System X(c)(). Antioxidants (Basel) 2021, 10.

97. Cui X, Zong S, Song W, Wang C, Liu Y, Zhang L, Xia P, Wang X, Zhao H, Wang L, Lu Z: Omaveloxolone ameliorates cognitive dysfunction in APP/PS1 mice by stabilizing the STAT3 pathway. Life Sci 2023, 335:122261.

98. Soni P, Ammal Kaidery N, Sharma SM, Gazaryan I, Nikulin SV, Hushpulian DM, Thomas B: A critical appraisal of ferroptosis in Alzheimer’s and Parkinson’s disease: new insights into emerging mechanisms and therapeutic targets. Front Pharmacol 2024, 15:1390798.

99. Joly-Amado A, Hunter J, Quadri Z, Zamudio F, Rocha-Rangel PV, Chan D, Kesarwani A, Nash K, Lee DC, Morgan D, et al: CCL2 Overexpression in the Brain Promotes Glial Activation and Accelerates Tau Pathology in a Mouse Model of Tauopathy. Front Immunol 2020, 11:997.

100. Bose S, Cho J: Role of chemokine CCL2 and its receptor CCR2 in neurodegenerative diseases. Arch Pharm Res 2013, 36:1039–1050.

101. Wan J, Fu AK, Ip FC, Ng HK, Hugon J, Page G, Wang JH, Lai KO, Wu Z, Ip NY: Tyk2/STAT3 signaling mediates beta-amyloid-induced neuronal cell death: implications in Alzheimer’s disease. J Neurosci 2010, 30:6873–6881.

102. Toral-Rios D, Patino-Lopez G, Gomez-Lira G, Gutierrez R, Becerril-Perez F, Rosales-Cordova A, Leon-Contreras JC, Hernandez-Pando R, Leon-Rivera I, Soto-Cruz I, et al: Activation of STAT3 Regulates Reactive Astrogliosis and Neuronal Death Induced by AbetaO Neurotoxicity. Int J Mol Sci 2020, 21.

103. Levin EC, Acharya NK, Sedeyn JC, Venkataraman V, D’Andrea MR, Wang HY, Nagele RG: Neuronal expression of vimentin in the Alzheimer’s disease brain may be part of a generalized dendritic damage-response mechanism. Brain Res 2009, 1298:194–207.

104. Zhang H, Wu LM, Wu J: Cross-talk between apolipoprotein E and cytokines. Mediators Inflamm 2011, 2011:949072.

105. Yamazaki Y, Zhao N, Caulfield TR, Liu CC, Bu G: Apolipoprotein E and Alzheimer disease: pathobiology and targeting strategies. Nat Rev Neurol 2019, 15:501–518.

106. Chen Y, Strickland MR, Soranno A, Holtzman DM: Apolipoprotein E: Structural Insights and Links to Alzheimer Disease Pathogenesis. Neuron 2021, 109:205–221.

107. Zhou Y, Song WM, Andhey PS, Swain A, Levy T, Miller KR, Poliani PL, Cominelli M, Grover S, Gilfillan S, et al: Human and mouse single-nucleus transcriptomics reveal TREM2-dependent and TREM2-independent cellular responses in Alzheimer’s disease. Nat Med 2020, 26:131–142.

108. Fernandez-Calle R, Konings SC, Frontinan-Rubio J, Garcia-Revilla J, Camprubi-Ferrer L, Svensson M, Martinson I, Boza-Serrano A, Venero JL, Nielsen HM, et al: APOE in the bullseye of neurodegenerative diseases: impact of the APOE genotype in Alzheimer’s disease pathology and brain diseases. Mol Neurodegener 2022, 17:62.

109. Jackson RJ, Hyman BT, Serrano-Pozo A: Multifaceted roles of APOE in Alzheimer disease. Nat Rev Neurol 2024, 20:457–474.

110. Verghese PB, Castellano JM, Holtzman DM: Apolipoprotein E in Alzheimer’s disease and other neurological disorders. Lancet Neurol 2011, 10:241–252.

111. Verghese PB, Castellano JM, Garai K, Wang Y, Jiang H, Shah A, Bu G, Frieden C, Holtzman DM: ApoE influences amyloid-beta (Abeta) clearance despite minimal apoE/Abeta association in physiological conditions. Proc Natl Acad Sci U S A 2013, 110:E1807–1816.

112. Castellano JM, Kim J, Stewart FR, Jiang H, DeMattos RB, Patterson BW, Fagan AM, Morris JC, Mawuenyega KG, Cruchaga C, et al: Human apoE isoforms differentially regulate brain amyloid-beta peptide clearance. Sci Transl Med 2011, 3:89ra57.

113. Holtzman DM, Bales KR, Wu S, Bhat P, Parsadanian M, Fagan AM, Chang LK, Sun Y, Paul SM: Expression of human apolipoprotein E reduces amyloid-beta deposition in a mouse model of Alzheimer’s disease. J Clin Invest 1999, 103:R15–R21.

114. Musiek ES, Holtzman DM: Three dimensions of the amyloid hypothesis: time, space and ’wingmen’. Nat Neurosci 2015, 18:800–806.

115. Rajmohan R, Reddy PH: Amyloid-Beta and Phosphorylated Tau Accumulations Cause Abnormalities at Synapses of Alzheimer’s disease Neurons. J Alzheimers Dis 2017, 57:975–999.

116. Bloom GS: Amyloid-beta and tau: the trigger and bullet in Alzheimer disease pathogenesis. JAMA Neurol 2014, 71:505–508.

117. Hefti MM, Kim S, Bell AJ, Betters RK, Fiock KL, Iida MA, Smalley ME, Farrell K, Fowkes ME, Crary JF: Tau Phosphorylation and Aggregation in the Developing Human Brain. J Neuropathol Exp Neurol 2019, 78:930–938.

118. Oakley DH, Chung M, Abrha S, Hyman BT, Frosch MP: beta-Amyloid species production and tau phosphorylation in iPSC-neurons with reference to neuropathologically characterized matched donor brains. J Neuropathol Exp Neurol 2024, 83:772–782.

119. Ochalek A, Mihalik B, Avci HX, Chandrasekaran A, Teglasi A, Bock I, Giudice ML, Tancos Z, Molnar K, Laszlo L, et al: Neurons derived from sporadic Alzheimer’s disease iPSCs reveal elevated TAU hyperphosphorylation, increased amyloid levels, and GSK3B activation. Alzheimers Res Ther 2017, 9:90.

120. Choi SH, Kim YH, Hebisch M, Sliwinski C, Lee S, D’Avanzo C, Chen H, Hooli B, Asselin C, Muffat J, et al: A three-dimensional human neural cell culture model of Alzheimer’s disease. Nature 2014, 515:274–278.

121. Gonzalez C, Armijo E, Bravo-Alegria J, Becerra-Calixto A, Mays CE, Soto C: Modeling amyloid beta and tau pathology in human cerebral organoids. Mol Psychiatry 2018, 23:2363–2374.

122. Knupp A, Mishra S, Martinez R, Braggin JE, Szabo M, Kinoshita C, Hailey DW, Small SA, Jayadev S, Young JE: Depletion of the AD Risk Gene SORL1 Selectively Impairs Neuronal Endosomal Traffic Independent of Amyloidogenic APP Processing. Cell Rep 2020, 31:107719.

123. Lee H, Aylward AJ, Pearse RV, 2nd, Lish AM, Hsieh YC, Augur ZM, Benoit CR, Chou V, Knupp A, Pan C, et al: Cell-type-specific regulation of APOE and CLU levels in human neurons by the Alzheimer’s disease risk gene SORL1. Cell Rep 2023, 42:112994.

124. Avey DR, Ng B, Vialle RA, Kearns NA, de Paiva Lopes K, Iatrou A, De Tissera S, Vyas H, Saunders DM, Flood DJ, et al: Uncovering Plaque-Glia Niches in Human Alzheimer’s Disease Brains Using Spatial Transcriptomics. bioRxiv 2024.

125. Liu T, Zhu B, Liu Y, Zhang X, Yin J, Li X, Jiang L, Hodges AP, Rosenthal SB, Zhou L, et al: Multi-omic comparison of Alzheimer’s variants in human ESC-derived microglia reveals convergence at APOE. J Exp Med 2020, 217.

126. Takano-Kawabe K, Matoba K, Nakamura Y, Moriyama M: Low Density Lipoprotein Receptor-related Protein 2 Expression and Function in Cultured Astrocytes and Microglia. Neurochem Res 2024, 49:199–211.

127. Quesnel MJ, Labonte A, Picard C, Bowie DC, Zetterberg H, Blennow K, Brinkmalm A, Villeneuve S, Poirier J, Alzheimer’s Disease Neuroimaging I, Group P-AR: Osteopontin: A novel marker of pre-symptomatic sporadic Alzheimer’s disease. Alzheimers Dement 2024, 20:6008–6031.

128. Qiu Y, Shen X, Ravid O, Atrakchi D, Rand D, Wight AE, Kim HJ, Liraz-Zaltsman S, Cooper I, Schnaider Beeri M, Cantor H: Definition of the contribution of an Osteopontin-producing CD11c(+) microglial subset to Alzheimer’s disease. Proc Natl Acad Sci U S A 2023, 120:e2218915120.

129. Akiyama H, Tooyama I, Kawamata T, Ikeda K, McGeer PL: Morphological diversities of CD44 positive astrocytes in the cerebral cortex of normal subjects and patients with Alzheimer’s disease. Brain Res 1993, 632:249–259.

130. Wang Q, Klyubin I, Wright S, Griswold-Prenner I, Rowan MJ, Anwyl R: Alpha v integrins mediate beta-amyloid induced inhibition of long-term potentiation. Neurobiol Aging 2008, 29:1485–1493.

131. Opanashuk LA, Mark RJ, Porter J, Damm D, Mattson MP, Seroogy KB: Heparin-binding epidermal growth factor-like growth factor in hippocampus: modulation of expression by seizures and anti-excitotoxic action. J Neurosci 1999, 19:133–146.

132. Maurya SK, Mishra J, Abbas S, Bandyopadhyay S: Cypermethrin Stimulates GSK3beta-Dependent Abeta and p-tau Proteins and Cognitive Loss in Young Rats: Reduced HB-EGF Signaling and Downstream Neuroinflammation as Critical Regulators. Mol Neurobiol 2016, 53:968–982.

133. Mansour HM, Fawzy HM, El-Khatib AS, Khattab MM: Repurposed anti-cancer epidermal growth factor receptor inhibitors: mechanisms of neuroprotective effects in Alzheimer’s disease. Neural Regen Res 2022, 17:1913–1918.

134. Wang L, Chiang HC, Wu W, Liang B, Xie Z, Yao X, Ma W, Du S, Zhong Y: Epidermal growth factor receptor is a preferred target for treating amyloid-beta-induced memory loss. Proc Natl Acad Sci U S A 2012, 109:16743–16748.

135. Jayaswamy PK, Vijaykrishnaraj M, Patil P, Alexander LM, Kellarai A, Shetty P: Implicative role of epidermal growth factor receptor and its associated signaling partners in the pathogenesis of Alzheimer’s disease. Ageing Res Rev 2023, 83:101791.

136. Woo RS, Lee JH, Yu HN, Song DY, Baik TK: Expression of ErbB4 in the neurons of Alzheimer’s disease brain and APP/PS1 mice, a model of Alzheimer’s disease. Anat Cell Biol 2011, 44:116–127.

137. Zhang H, Zhang L, Zhou D, Li H, Xu Y: ErbB4 mediates amyloid beta-induced neurotoxicity through JNK/tau pathway activation: Implications for Alzheimer’s disease. J Comp Neurol 2021, 529:3497–3512.

138. Shaftel SS, Griffin WS, O’Banion MK: The role of interleukin-1 in neuroinflammation and Alzheimer disease: an evolving perspective. J Neuroinflammation 2008, 5:7.

139. Kitazawa M, Cheng D, Tsukamoto MR, Koike MA, Wes PD, Vasilevko V, Cribbs DH, LaFerla FM: Blocking IL-1 signaling rescues cognition, attenuates tau pathology, and restores neuronal beta-catenin pathway function in an Alzheimer’s disease model. J Immunol 2011, 187:6539–6549.

140. Lopez-Rodriguez AB, Hennessy E, Murray CL, Nazmi A, Delaney HJ, Healy D, Fagan SG, Rooney M, Stewart E, Lewis A, et al: Acute systemic inflammation exacerbates neuroinflammation in Alzheimer’s disease: IL-1beta drives amplified responses in primed astrocytes and neuronal network dysfunction. Alzheimers Dement 2021, 17:1735–1755.

141. Labandeira-Garcia JL, Costa-Besada MA, Labandeira CM, Villar-Cheda B, Rodriguez-Perez AI: Insulin-Like Growth Factor-1 and Neuroinflammation. Front Aging Neurosci 2017, 9:365.

142. Sohrabi M, Floden AM, Manocha GD, Klug MG, Combs CK: IGF-1R Inhibitor Ameliorates Neuroinflammation in an Alzheimer’s Disease Transgenic Mouse Model. Front Cell Neurosci 2020, 14:200.

143. Zhou Y, Chen Y, Xu C, Zhang H, Lin C: TLR4 Targeting as a Promising Therapeutic Strategy for Alzheimer Disease Treatment. Front Neurosci 2020, 14:602508.

144. Zhang X, Sun D, Zhou X, Zhang C, Yin Q, Chen L, Tang Y, Liu Y, Morozova-Roche LA: Proinflammatory S100A9 stimulates TLR4/NF-kappaB signaling pathways causing enhanced phagocytic capacity of microglial cells. Immunol Lett 2023, 255:54–61.

145. Frew J, Baradaran-Heravi A, Balgi AD, Wu X, Yan TD, Arns S, Shidmoossavee FS, Tan J, Jaquith JB, Jansen-West KR, et al: Premature termination codon readthrough upregulates progranulin expression and improves lysosomal function in preclinical models of GRN deficiency. Mol Neurodegener 2020, 15:21.

146. Lee C, Frew J, Weilinger NL, Wendt S, Cai W, Sorrentino S, Wu X, MacVicar BA, Willerth SM, Nygaard HB: hiPSC-derived GRN-deficient astrocytes delay spiking activity of developing neurons. Neurobiol Dis 2023, 181:106124.

147. Rose SE, Frankowski H, Knupp A, Berry BJ, Martinez R, Dinh SQ, Bruner LT, Willis SL, Crane PK, Larson EB, et al: Leptomeninges-Derived Induced Pluripotent Stem Cells and Directly Converted Neurons From Autopsy Cases With Varying Neuropathologic Backgrounds. J Neuropathol Exp Neurol 2018, 77:353–360.

148. Stine WB, Jungbauer L, Yu C, LaDu MJ: Preparing synthetic Abeta in different aggregation states. Methods Mol Biol 2011, 670:13–32.

149. Dana H, Mohar B, Sun Y, Narayan S, Gordus A, Hasseman JP, Tsegaye G, Holt GT, Hu A, Walpita D, et al: Sensitive red protein calcium indicators for imaging neural activity. Elife 2016, 5.

150. Wang Y, DelRosso NV, Vaidyanathan TV, Cahill MK, Reitman ME, Pittolo S, Mi X, Yu G, Poskanzer KE: Accurate quantification of astrocyte and neurotransmitter fluorescence dynamics for single-cell and population-level physiology. Nat Neurosci 2019, 22:1936–1944.

151. Ku T, Swaney J, Park JY, Albanese A, Murray E, Cho JH, Park YG, Mangena V, Chen J, Chung K: Multiplexed and scalable super-resolution imaging of three-dimensional protein localization in size-adjustable tissues. Nat Biotechnol 2016, 34:973–981.

152. Raudvere U, Kolberg L, Kuzmin I, Arak T, Adler P, Peterson H, Vilo J: g:Profiler: a web server for functional enrichment analysis and conversions of gene lists (2019 update). Nucleic Acids Res 2019, 47:W191–W198.

